# A neofunctionalized flowering antagonist channeled Solanaceae evolution

**DOI:** 10.64898/2026.02.20.707063

**Authors:** Hagai Shohat, Danielle Ciren, Andrea Arrones, Iacopo Gentile, Srividya Ramakrishnan, Anat Hendelman, Katharine M. Jenike, Nicole L. Brown, Luiz Carlos Machado, Nurdan Kuru, Jose Luna-Ramos, Michael J. Passalacqua, James W. Satterlee, Blaine Fitzgerald, Virginia Baraja-Fonseca, Gina M. Robitaille, Brooke M. Seman, Jesse Gillis, Joyce Van Eck, Adam Siepel, Jaime Prohens, Michael C. Schatz, Zachary B. Lippman

## Abstract

Neofunctionalization is a fate of gene duplication, classically defined as the acquisition of a new molecular function^1,2^. Rather than evolving to function autonomously, neofunctionalized genes may also remain embedded within their ancestral regulatory networks, potentially reshaping the genetic trajectories through which phenotypic change occurs. Testing this hypothesis within the polygenic architecture of flowering time in plants, we leveraged a pan-genetic platform comprising ten Solanaceae species and show that a paralogue of the flowering hormone *florigen* neofunctionalized into a flowering antagonist and was repeatedly mutated to accelerate flowering in wild plants and during crop domestication across 50 million years of evolution. Independent *cis*-regulatory and coding mutations in *SELF-PRUNING 5G* (*SP5G*) enabled rapid flowering in the wild progenitor of domesticated tomato from the Americas, as well as three eggplant species independently domesticated in Asia and Africa. We further found that degradation of conserved *cis*-regulatory sequences reduced *SP5G* expression and flowering time in wild species, including plants endemic to the arid Australian desert. Together with similar examples across diverse species, regulatory networks, and traits^3–6^, our findings suggest the emergence of opposing functions among paralogues through antagonistic neofunctionalization can create evolutionary contingencies that channel future phenotypic change and adaptation through parallel genetic paths.

## INTRODUCTION

Two fundamental questions in evolutionary genetics are whether similar phenotypes among related lineages arise through parallel or independent genetic paths, and how the architectures of gene-regulatory networks bias evolution toward one trajectory over another^7–9^. Diversification of gene-regulatory networks, particularly those controlling quantitative traits, can promote evolvability by expanding the phenotypic space available for adaptation^10^. Gene-regulatory networks evolve through mutations in existing network components, as well as through gene duplication^11^. Following duplication, redundant paralogues initially confer network robustness by tolerating the accumulation of deleterious mutations^12^. If one or the other redundant paralogue does not pseudogenize or get eliminated from the genome, fortuitous *cis*-regulatory or coding mutations may partition ancestral functions (subfunctionalization) or alter expression patterns or protein function to generate novel activities (neofunctionalization)^1,2^. These fates of surviving paralogues may not be discrete, and instead represent a continuum of functional divergence, with gradual transition between subfunctionalization to neofunctionalization^13^.

Paralogue neofunctionalization has long been viewed as a source of new gene family member function that facilitates the emergence of new traits^2^. However, rather than becoming functionally decoupled from their ancestral regulatory networks, neofunctionalized paralogues can rewire interactions within the networks in which they remain embedded, thereby expanding the regulatory capacity available for phenotypic changes. Neofunctionalized genes may also escape ancestral redundancy and introduce evolutionary contingencies that redirect the genetic paths of future evolutionary change^14^. However, testing this concept requires expanding the phenotypic, organismal, and taxonomic scopes in which neofunctionalized paralogues operate within their ancestral regulatory network.

The *florigen-antiflorigen* hormone network is conserved across flowering plants (angiosperms) and controls the timing of the transition to reproductive growth (flowering time), a critical polygenic adaptive trait in both natural and agricultural environments^15^. This network is composed of mobile peptides encoded by flowering-promoting homologs of *FLOWERING LOCUS T* (*FT*, encoding *florigen*) and antagonistic members of the same gene family, *TERMINAL FLOWER 1* (*TFL1*, encoding *antiflorigen*)^16,17^. Across angiosperms, the *florigen-antiflorigen* network has diversified through gene duplication followed by lineage-specific paralogue diversification, including neofunctionalization^18^. Mutations of core and peripheral components within *florigen/antiflorigen* networks or their surrounding transcriptional regulators have been repeatedly selected during angiosperm and crop evolution^19,20^. Thus, flowering time, with its underlying polygenic control, is an opportune system to dissect whether parallel phenotypic change among related species arose from diverse genetic paths or was channeled by evolutionary contingencies introduced by the emergence of new paralogues.

The nightshade (Solanaceae) family comprises nearly 3,000 species, including numerous globally and locally cultivated crops, including tomatoes, potatoes, peppers, and eggplants^21^. Solanaceous crops and their wild relatives evolved across diverse ecological and geographical habitats^22^, many of which now extend far beyond their centers of origin through human selection and migration (**Fig. 1a**), owing to extensive variation in flowering time that enhanced adaptability to contrasting environments (**Fig. 1b,c**). Here, leveraging ten species into a pan-genetic platform spanning diverse geographical origins and evolutionary timescales, we use flowering time as a tractable phenotypic system to reveal how an evolutionary contingency repeatedly shaped phenotypic evolution across 50 million years of Solanaceae diversification.

**Figure 1.**
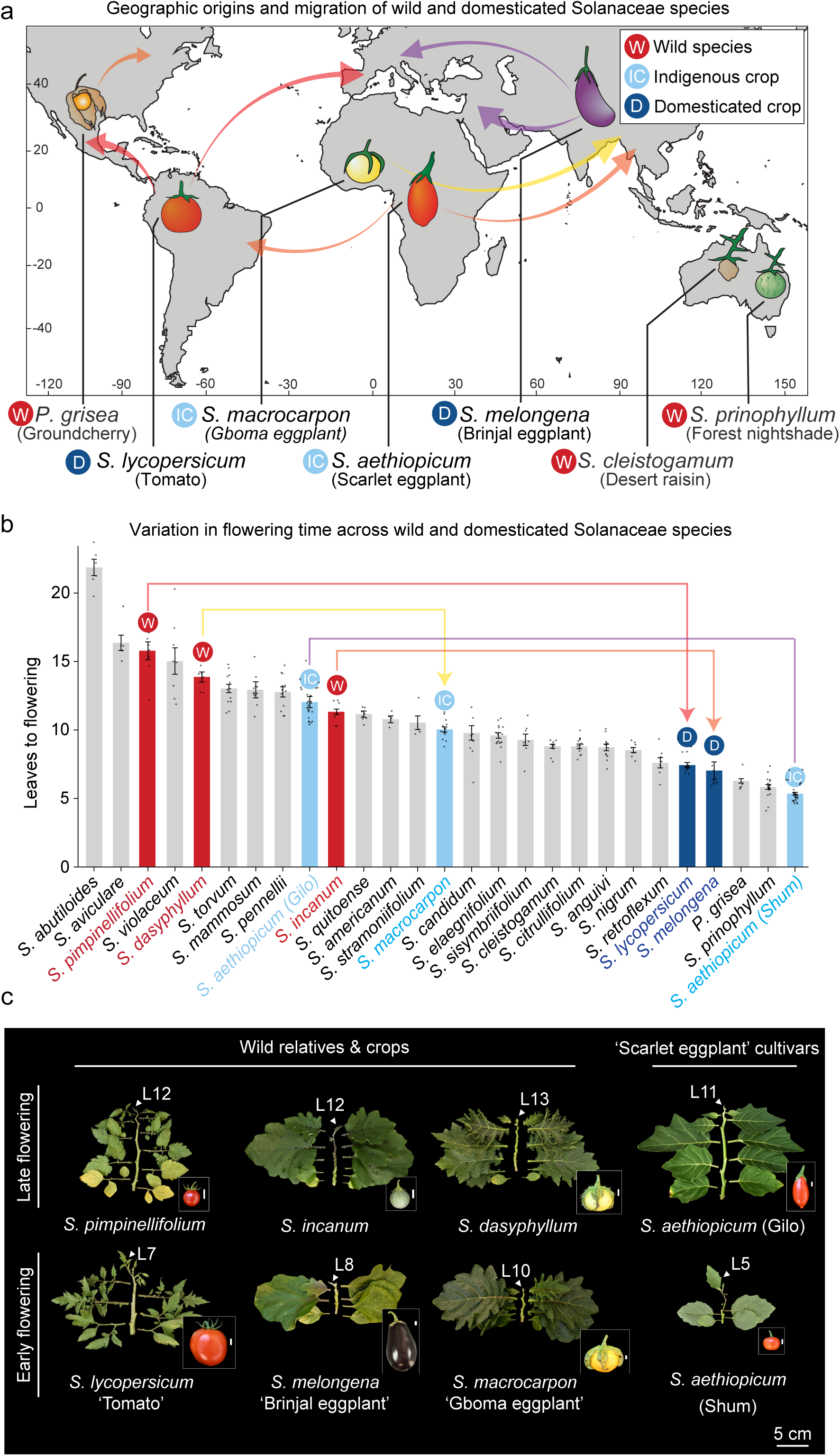
Recurrent variation in flowering time underlies independent Solanaceae adaptations. **(a)** Approximate geographic origins and migration paths of selected wild and domesticated Solanaceae species and their uses: wild species (W), indigenous crop (IC), and domesticated crop (D). **(b)** Flowering-time variation across Solanaceae species. Bar charts present means ±s.e. Sample size (n) is presented in **Supplementary Table 3**, representing the number of plants quantified per species. **(c)** Images of shoots and fruits of *Solanum* wild relatives and crop cultivars. White arrowheads indicate leaf number (L) before the first inflorescence. Scale bars: 5 cm (shoots) and 1 cm (fruits).

## RESULTS

### *SP5G* mutations in tomato domestication

In many angiosperms, flowering time is regulated by seasonal photoperiod cues. Modification of day-length sensitivity, including day-neutral flowering, was a key step in multiple crop domestications, enabling broader geographical cultivation^23^. In the wild progenitor of domesticated tomato, *Solanum pimpinellifolium*, this agricultural adaptation was achieved through the *antiflorigen* gene *SELF-PRUNING 5G* (*SP5G,* Solyc05g053850), which is upregulated under long days to repress the transition to flowering on primary and side shoots^24^. *SP5G* is a *florigen*-derived paralogue that evolved antiflorigenic activity through still uncharacterized missense mutations in its external loop domain^24^. Two conspicuous substitutions within this domain (Y139 and W143) represent a plausible recurrent target by which *florigens* neofunctionalize into *antiflorigens*, supported by parallel changes in homologous positions in neofunctionalized antiflorigens of sugar beet, onion, and eelgrass^25–27^. In addition, still unresolved modification of *SP5G cis*-regulation conferred a diurnal expression pattern, peaking in cotyledons to prevent precocious flowering^24^.

Our prior work found that a downregulated *SP5G* allele enabled rapid and day-neutral flowering in domesticated tomato (*S. lycopersicum*)^24^. This phenotypic change was suggested to result from a 52-bp deletion in the 3’-UTR (*SlycSP5G^Δ^*^52^) disrupting a chromatin loop between the promoter and the intact 52-bp sequence^28^. Because this deletion and its sufficiency was never validated, we used CRISPR/Cas9 to target the intact 52-bp region in an introgression line carrying a functional *SP5G* allele from the wild tomato relative *S. pennellii* (*SpenSP5G*) in a standard domesticated genotype (M82)^29^. We isolated an 87-bp deletion allele (*SpenSP5G^cr-Δ87^*) that removed 43 of the 52-bp putative causal domestication variant (**Fig. 2a, Supplementary Table 1**). This deletion eliminated a predicted CYCLING DOF FACTOR (CDF) binding site, a circadian clock transcriptional regulator controlling photoperiodic flowering in tomato^30^. However, the engineered allele *SpenSP5G^cr-Δ^*^87^ showed a weaker effect on flowering time (mean 11.1 leaves produced before the first inflorescence) and *SP5G* expression compared with the natural domestication allele (mean 6.9 leaves in *SlycSP5G^Δ52^*) (**Fig. 2b,c**). This indicated that the 52-bp deletion does not account for the full effect of the true domestication allele.

**Figure 2.**
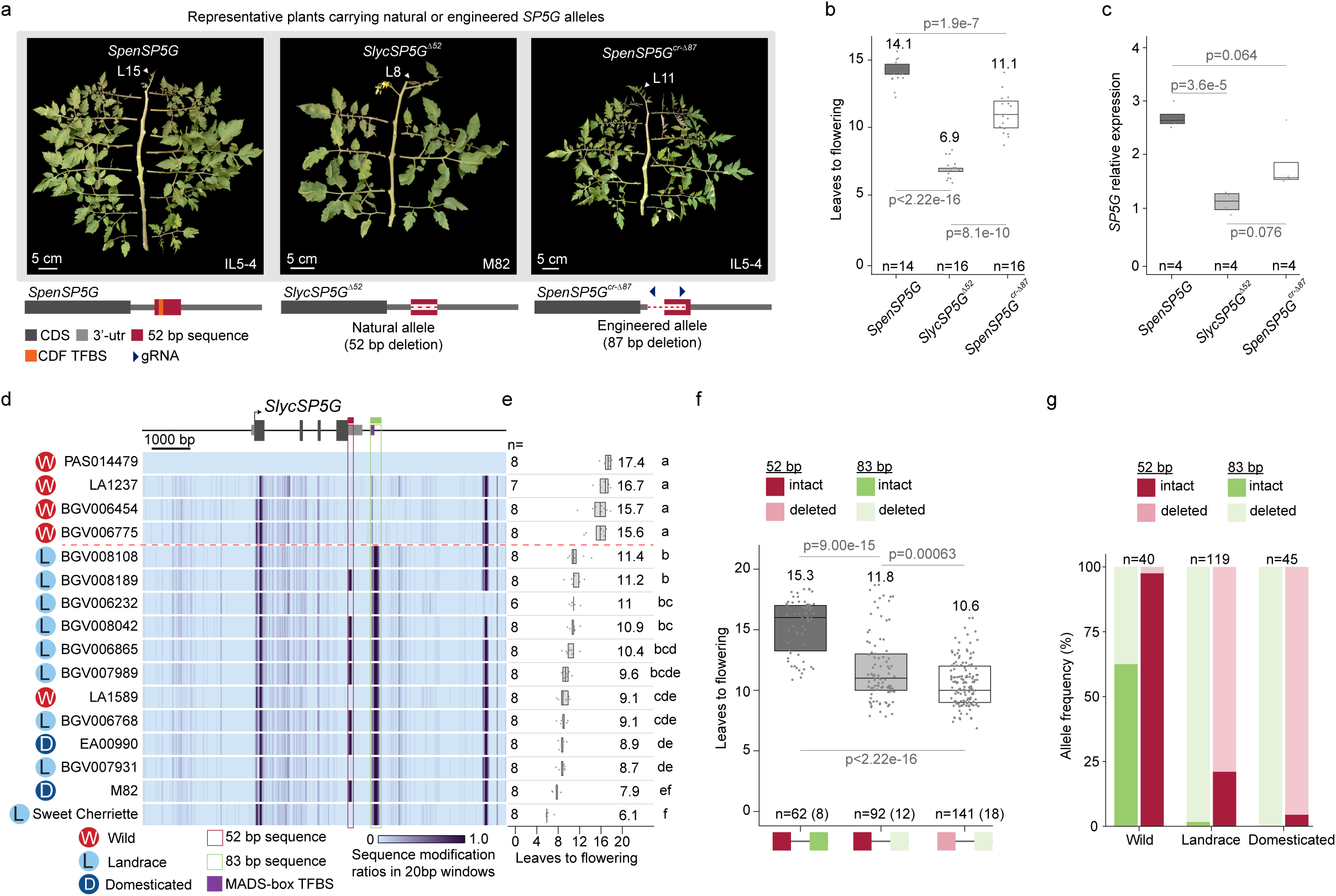
Stepwise *cis*-regulatory deletions in *SP5G* accelerated flowering in tomato domestication. **(a)** Representative plants carrying functional and mutated tomato *SP5G* alleles and their sequence description. White arrowheads indicate leaf number (L) before the first inflorescence. Dashed red lines indicate deletions. **(b)** Effects on flowering time of tomato *SP5G* alleles. **(c)** Expression analysis (RT-qPCR) of tomato *SP5G* alleles 4h after sunrise. Sample size (n) represents the number of cotyledons taken from different plants. A two-tailed, two-sample t-test was performed to compare means. **(d)** Heatmaps of *SP5G* haplotypes from accessions of *S. pimpinellifolium* (W; Wild), early (L; Landrace) and modern (D; Domesticated) varieties. The haplotypes were encoded; perturbations (purple intensity) are represented as the degree of sequence modification within 20-bp windows relative to *SpimSP5G* (accession PAS014479). **(e)** Flowering time of the corresponding accessions in (d). **(f)** Flowering time of accessions carrying the intact or deleted 83-bp and 52-bp sequences. Red (52-bp) or green (83-bp) squares denote intact sequence, lighter colors denote deletion. Numbers in parentheses represent the number of accessions per genotype. Means in (b), (c), (e) are shown above box plots. Sample size (n) in (b), (e), (f) represents the number of quantified plants per genotype. Lowercase letters in (b), (e), (f) indicate significant difference (*p* < 0.05) based on one-way ANOVA with post-hoc Tukey-Kramer HSD test. Box plots in (b), (c), (e), (f) show the 25th, 50th (median) and 75th percentiles. (**g**) Allele frequency of the 83-bp and 52-bp deletions across the tomato pan-genome^32,34^. Sample size (n) represents the number of tested accessions per genotype.

Given our previous *cis*-regulatory editing results demonstrating widespread redundancy among *cis*-regulatory sequences^31^, we reasoned that additional *SP5G cis*-regulatory variants were likely involved. Using our tomato pan-genome spanning 100 wild (*S. pimpinellifolium*), early-domesticated (landraces), and modern genotypes^32^, we aligned *cis*-regulatory sequences surrounding *SP5G* from 16 genotypes spanning these major subclades and identified a previously overlooked 83-bp deletion located 370 bp downstream of the 52-bp deletion. The 83-bp sequence was broadly conserved across the *Solanum* (**Extended Data Fig. 1a**), and its loss eliminated a MADS-box transcription factor binding site associated with flowering^33^ (**Fig. 2d-f**). Tellingly, several Peruvian and Mexican *S. pimpinellifolium* and landrace accessions, which are the most likely direct progenitors of cultivated tomato^34^, carry the 83-bp deletion and show reduced *SP5G* expression and earlier flowering compared with accessions carrying an intact allele (**Extended Data Fig. 1b,c, Fig. 2d,e, Supplementary Table 2 and 3**).

This suggests the 83-bp deletion arose initially as a standing variant in progenitor *S. pimpinellifolium* accessions. In support, allele-frequency analysis across 204 accessions^32,34^ showed that 37% of *S. pimpinellifolium* accessions carry the deletion, increasing to near (98%) and complete fixation in landrace and domesticated varieties, respectively. The 52-bp deletion allele is highly enriched (79%) in landraces already carrying the 83-bp deletion, and the allele with both *cis*-regulatory deletions exists in 95% of tested domesticated genotypes (**Fig. 2g, Supplementary Table 2**). The near fixation of the 83-bp deletion allele in landraces is consistent with reduced nucleotide diversity (*π*) at this locus, previously interpreted as evidence of a domestication-associated selective sweep^35^. We extended this analysis to include 106 accessions from our dataset and a recently assembled collection of 452 accessions^36^. Both datasets revealed elevated population differentiation (*F_ST_*) at *SP5G* between wild and landrace accessions relative to the genomic background (**Supplementary Notes**).

To further investigate the evolutionary forces acting on *SP5G*, we carried out genome-wide scans for selective sweeps using nucleotide diversity (*π*, Tajima’s *D*) and haplotype-based statistics (*nS*L, XP-*nS*L). These exhaustive tests, based on all available tomato pan-genome data, produced mixed results which did not provide consistent evidence of positive selection at *SP5G* (**Supplementary Notes**). Importantly, simulations under the tomato demographic model^34^ indicated that, even with the larger dataset, we are near the limits of statistical power for detecting sweeps, with particularly low power for soft sweeps from standing variation at high initial frequency, as found in *SP5G* (**Supplementary Fig. 1 and 2**). To further assess these limitations, we applied the same tests to eight established domestication loci and similarly found that most tests were inconclusive (**Supplementary Notes**). Given these power limitations, particularly for soft sweeps from standing variation^37^, we do not interpret the absence of consistent statistical evidence for a sweep at *SP5G* or other established domestication loci as evidence against selection.

To further assess phenotypic effects and the evolutionary importance of the *cis*-regulatory deletions at *SP5G*, we performed co-segregation analyses in F2 populations segregating for each variant individually. A population derived from a cross between late-and early-flowering *S. pimpinellifolium* accessions segregating for the 83-bp deletion showed that plants carrying the 83-bp deletion flowered earlier than those with the intact sequence (mean 10.3 leaves versus 13.2) (**Extended Data Fig. 1d**). Though attempts to edit the 83-bp region were unsuccessful due to poor gRNA accessibility, our functional analyses support causality. We also evaluated the effect of the 52-bp deletion in a fixed 83-bp deletion background using an F2 population. Notably, the combined deletions further accelerated flowering, nearly recapitulating the early-flowering phenotype of cultivated tomato (mean 10.4 leaves versus 8.1) (**Extended Data Fig. 1e, Fig. 2d**). Although contributions from other *cis*-regulatory variants cannot be excluded, our findings strongly support that flowering-time adaptation during tomato domestication proceeded through accumulation of two *cis*-regulatory deletions that stepwise mitigated *SP5G* activity.

### *SP5G* is a neofunctionalized antiflorigen

An important question raised by our findings is when *SP5G* arose and acquired antiflorigenic activity, and whether its function is unique more broadly among its *FT-like* paralogues *SFT/FT* (Solyc03g063100), *SP6A* (Solyc05g055660), *FTL1a* (Solyc11g008650), and *FTL1b* (Solyc11g008640)^24^. Phylogenetic analysis of protein sequences revealed that the closest homologs in species from the sister family Convolvulaceae, including sweet potato (*Ipomoea batatas*) and Japanese morning glory (*I. nil*) clustered outside the Solanaceae SP5G sub-clade (**Fig. 3a, Extended Data Fig. 2a**). Multiple sequence alignment of 22 Solanaceae SP5G orthologues with homologous FT proteins showed that the characteristic external loop domain of SP5G is conserved across Solanaceae, whereas the closest Convolvulaceae homologs retained a canonical florigen sequence (**Fig. 3b, Extended Data Fig. 2b, Supplementary Fig. 3a**). Moreover, *SP5G* orthologues are consistently expressed in vegetative tissues (**Extended Data Fig. 2c-g**), indicating a conserved regulatory role across Solanaceae. Together, these results indicate that *SP5G* is a Solanaceae gene that diverged following duplication of an ancestral *florigen*.

**Figure 3.**
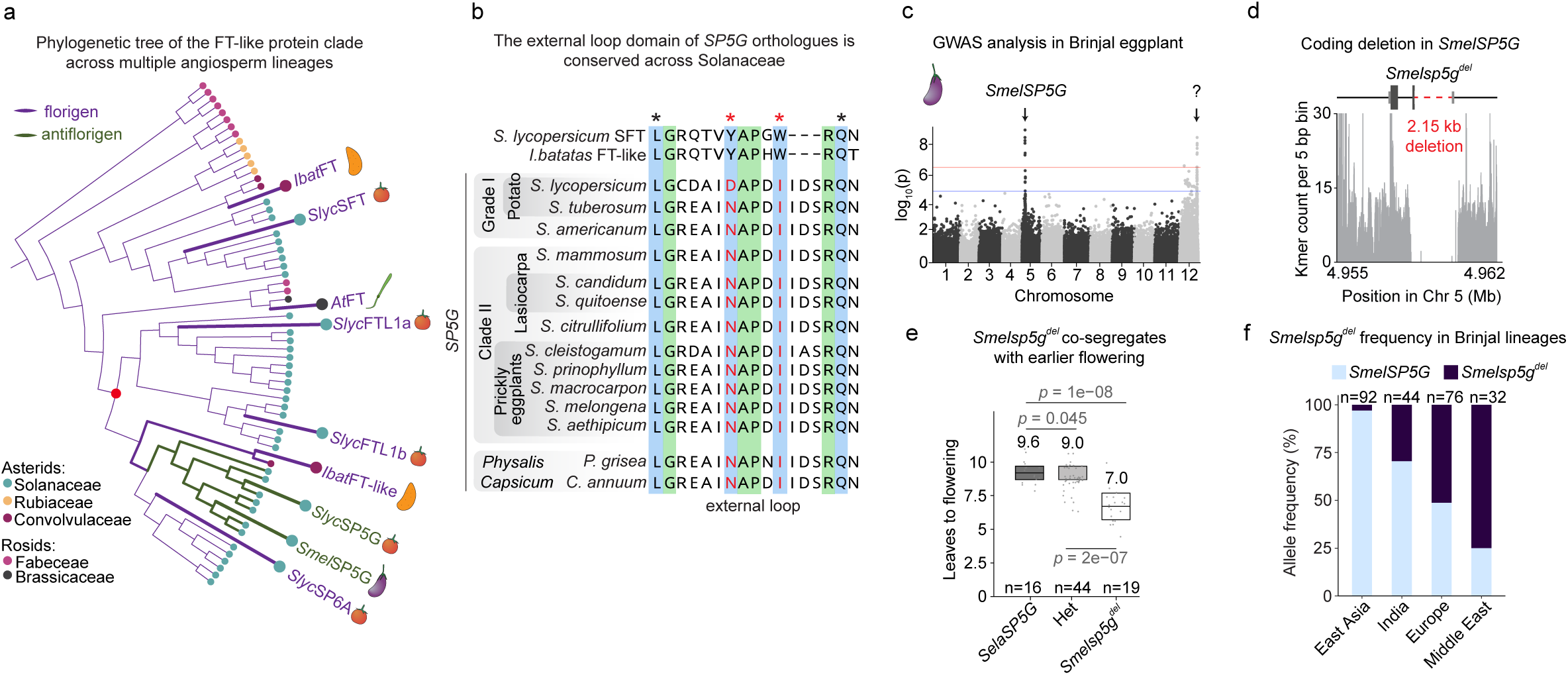
*SP5G* is a neofunctionalized antiflorigen and its deletion accelerates flowering in Brinjal eggplants. **(a)** Protein-based phylogenetic tree of the FLOWERING LOCUS T (FT) clade of the *florigen* gene family across angiosperm families. Green lines denote the Solanaceae SP5G sub-clade. **(b)** Partial alignment of the external loop domain of Solanaceae SP5G orthologues. The tomato FT (SFT, SINGLE FLOWER TRUSS) and the closest sweet potato (*Ipomoea batatas*) homolog retain the canonical florigen motif in the external loop domain. Asterisks and blue color mark key florigenic amino acid residues with red asterisks marking critical changes in SP5G orthologues relative to florigens. Green color marks identical residues. (**c**) Manhattan plot of the GWAS in Brinjal eggplant showing a peak on chromosome 5 overlapping *SmelSP5G*. The red and blue horizontal lines represent a multiple-testing correction using the Bonferroni and FDR significance thresholds at *p* = 0.05, respectively. **(d)** Kmer pile-up at *SmelSP5G* identifies a coding deletion, and *Smelsp5g^del^* schematic above shows loss of third and fourth exons. **(e)** Flowering time of homozygous *Smelsp5g^del^* plants compared to *SelaSP5G* homozygotes and heterozygotes from a BC4S1 segregating population. Sample size (n) represents the number of plants quantified. Means are shown above box plots. Box plots show the 25th, 50th (median) and 75th percentiles. A two-tailed, two-sample t-test was performed to compare means. Box plots show the 25th, 50th (median) and 75th percentiles. **(f)** Allele frequency of the deletion allele across Brinjal eggplants from different geographical regions^45,46^. Geographical origins were assigned based on passport data from Gaccione et al., (2025)^46^ and summarized in **Supplementary Table 5**. Sample size (n) represents the number of accessions per geographic region.

A remaining question is whether *SP5G* antiflorigenic activity is unique or has a more ancestral property shared with other Solanaceae *FT-like* paralogues. While *SP6A* is a florigenic *FT* homolog that promotes tuberization in potato and retains flowering-promoting activity^38^, previous studies have suggested that the tomato paralogues *FTL1a* and *FTL1b* may also act as *antiflorigens*^39,40^. However, more recent studies support a florigenic role for *FTL1a* in tomato and potato^41,42^, consistent with the presence of florigen-associated residues^43^ (**Fig. 3b, Supplementary Fig. 3b,c**).

To further validate the function of these paralogues, we generated CRISPR *FTL1a* knockouts in *S. prinophyllum* (Forest nightshade), a wild species belonging to the prickly eggplant clade^44^. Homozygous *FTL1a* mutants (*Spriftl1a^cr^*, Solpri1_04g026960), flowered later than WT plants (7.0 versus 5.7 leaves) (**Extended Data Fig. 2h)**, supporting a conserved florigenic role across multiple Solanaceae lineages. Moreover, CRISPR knockouts of *FTL1b* in tomato (*Slycftlb^cr^*) had no effect on flowering time (**Extended Data Fig. 2i)**. This is consistent with a plausible negligible evolutionary role of this paralogue in flowering-time regulation based on frequent gene truncation or pseudogenization across at least half of 22 sequenced *Solanum* species^22^ (**Supplementary Fig. 3c)**. Our genome-wide scans likewise showed mixed selection signals across these paralogues due to power limitations, although loss-of-function allele of *FTL1a*, which was reported to delay flowering under short days^41^, arose in landraces and showed a strong signal, consistent with a more readily detectable hard sweep (**Supplementary Notes**). These combined phylogenetic and functional-genetic analyses demonstrate that *SP5G* is a Solanaceae-specific paralogue that acquired antiflorigenic function following gene duplication, positioning *SP5G* as a potent antagonist within the Solanaceae *florigen/antiflorigen* network.

### Recurrent mutation of *SP5G* in eggplants

We hypothesized that *SP5G* neofunctionalization could have created an evolutionary contingency that channels parallel flowering-time evolution across solanaceous crops. Brinjal eggplant (*S. melongena*)^45^, a global major crop like tomato, flowers earlier than its wild relative (mean 7 leaves versus 12 leaves in *S. incanum*) (**Fig. 1b,c**). Brinjal eggplant originated in Asia and belongs to the prickly eggplant clade^46^. To dissect the genetic basis of early flowering in Brinjal eggplant, we conducted a genome-wide association study (GWAS) on 283 lines derived from a multiparent population (MAGIC) composed of seven cultivated accessions and a wild relative *S. incanum*^47^. Because genomic resources were lacking for the founder accessions, we generated a chromosome-scale genome assembly and gene annotation for an early-flowering accession H15 [post-contamination screened contig N50 (average weighted contig length) = 95.7 Mb, BUSCO Completeness = 98.7, **Supplementary Table 4**]. Strikingly, we identified association peaks on chromosomes 12 and 5, the latter aligned with the *SP5G* locus of Brinjal (*SmelSP5G,* SMEL5_05g005030) (**Fig. 3c**), along with a 2.15-kb coding deletion that eliminates nearly half of the *SmelSP5G* gene body in the early-flowering commercial genotypes H15 and DH ECAVI (**Extended Data Fig. 3a**)^47^. We validated the earlier flowering effect of the deletion allele (*Smelsp5g^del^*) using genotype-phenotype association analyses across the 283 MAGIC lines in two independent environments (**Extended Data Fig. 3b,c**), and by co-segregation analysis using a population derived from a cross between a Brinjal accession (MEL3) and the wild relative *S. elaeagnifolium* (mean 7 versus 9.6 leaves) (**Fig. 3e**).

Unlike the single domestication origin of tomato^34^, which likely facilitated fixation of the *cis*-regulatory *SP5G* deletions in modern cultivars, Brinjal experienced two independent domestications in Southeast Asia and India^45^. European accessions, including the two MAGIC founders carrying the 2.15-kb deletion, were proposed to derive from the Indian lineage^45^. Using *Panagram*, our alignment-free pan-genome browser (see **Methods**), we visualized *SmelSP5G* haplotypes across 341 accessions (**Supplementary Fig. 4, Supplementary Table 5**)^45,46^. This analysis revealed enrichment of the 2.15-kb deletion allele in genotypes of Indian origin compared with Southeast Asian accessions (30% versus 1%) (**Fig. 3e, Supplementary Fig. 4, Supplementary Table 5**). The increase in frequency of the 2.15-kb deletion from Indian to Middle Eastern (75%) and European (51%) accessions, along with its near absence in Southeast Asian genotypes, align with the independent domestication and migration routes of Brinjal eggplant^45^. Applying the same genome-wide scan pipeline to 324 accessions in our eggplant dataset, neither *SP5G* nor its paralogues showed consistent evidence of selective sweeps across tests, likely due to high initial frequency of *Smelsp5g^del^* which suggests that sweep detection at this locus may face similar power limitations as the tomato *SP5G* (**Supplementary Notes**). These findings show that at least two independent loss-of-function mutations of *SP5G* accelerated flowering time in tomato and Brinjal eggplant, two distinct lineages of major solanaceous crops.

These results prompted us to ask whether *SP5G* alleles underlie parallel flowering-time evolution more broadly across solanaceous crops. We leveraged our pan-genetic platforms of two regionally important indigenous crops from Africa: the Gboma (*S. macrocarpon*) and Scarlet (*S. aethiopicum*) eggplants^22,48^ (**Fig. 1a**). The Gboma eggplant flowers substantially earlier than its wild progenitor *S. dasyphyllum* (mean 10 versus 12.9 leaves) (**Fig. 1b,c**). We generated an F2 mapping population between a cultivated Gboma (accession PI 441914)^44^ and *S. dasyphyllum*, and performed quantitative trait locus-sequencing (QTL-seq). This analysis identified two peaks on chromosomes 4 and 12, the latter coinciding with the orthologous position of the Gboma *SP5G* (*SmacSP5G,* Solmac3_12g004790) (**Fig. 4a, Extended Data Fig. 4a**). Using existing reference Gboma genome^44^ (accession PI 441914) together with newly generated chromosome-scale assemblies for two other accessions (PI 441915 and SC129) and *S. dasyphyllum* [post-contamination screened contig N50 = 107.5, 111.1 and 105.7 Mb, BUSCO Completeness = 98.3 and 98.7, respectively, **Supplementary Table 4**], we identified a 3.7-kb transposable-element (TE) insertion located 580 bp upstream of *SmacSP5G*, which was absent from *SdasSP5G* (**Fig. 4b**). Notably, all three Gboma accessions carrying the TE insertion(*SmacSP5G^TE^*) showed only residual cotyledon expression relative to *SdasSP5G* (**Fig. 4c**), and this insertion co-segregated with earlier flowering (**Fig. 4d, Extended Data Fig. 4b, Supplementary Table 6**). These results suggest that TE-mediated loss of *SmacSP5G* expression contributes to rapid flowering in Gboma eggplant.

**Figure 4.**
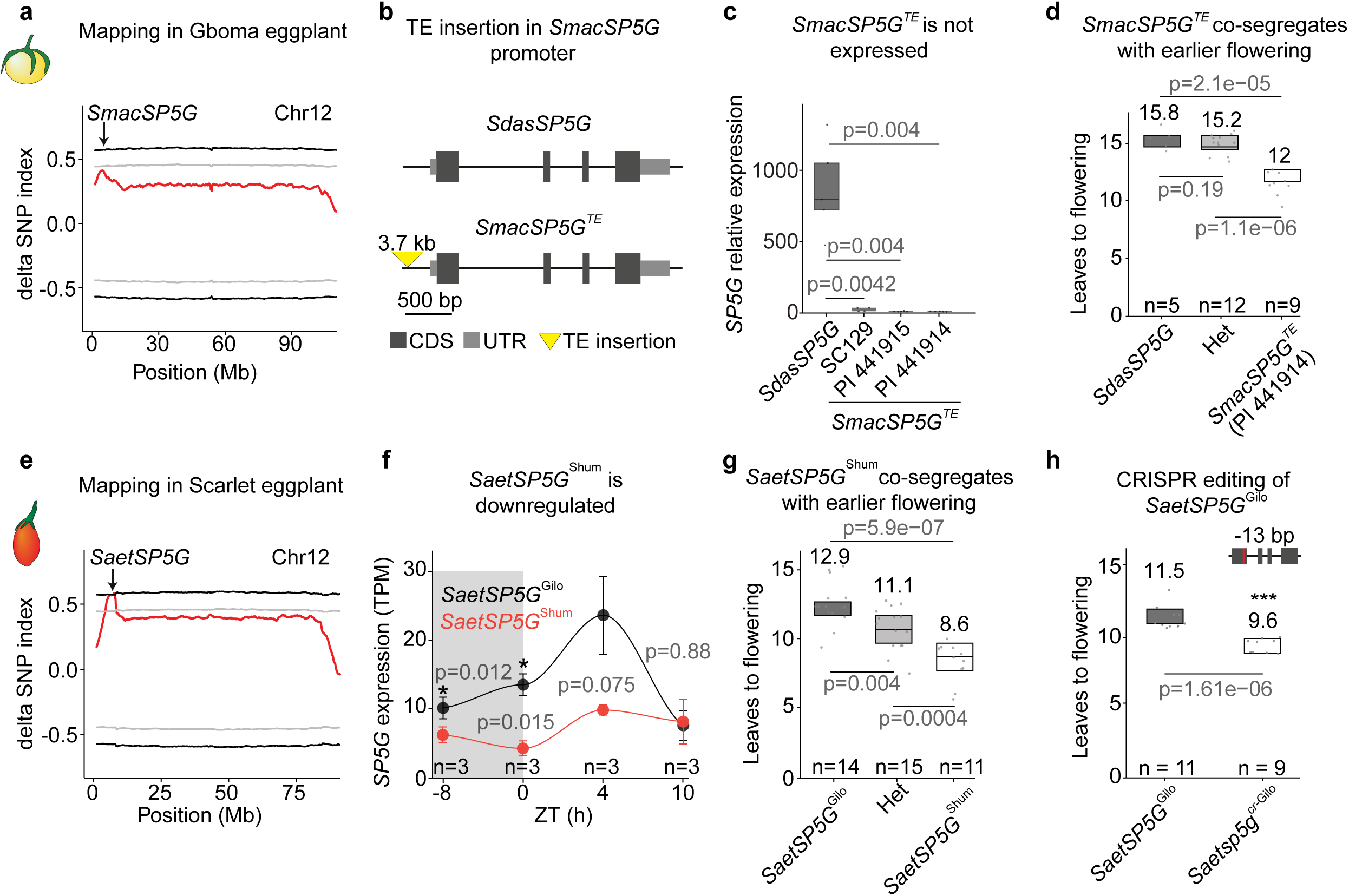
Recurrent *SP5G* mutation drove flowering-time change in African eggplants. (**a**) Quantitative trait locus mapping-by-sequencing (QTL-seq) identifies a major-effect flowering-time QTL on chromosome 12 in Gboma eggplant overlapping with *SP5G*. **(b)** Schematic of *SdasSP5G* and the *PIF/Harbinger* transposon-insertion allele *SmacSP5G^TE^*. **(c)** RT-qPCR expression analysis of *SP5G* in *S. dasyphyllum* compared to three accessions of Gboma eggplant. Five cotyledons per genotype taken from different plants were used as biological replicates. **(d)** Flowering time of plants homozygous and heterozygous for *SdasSP5G* and *SmacSP5G^TE^* from an F2 segregating population. Significant differences (*p* < 0.05) were determined by one-way ANOVA with post-hoc Tukey–Kramer HSD test. **(e)** QTL-seq identifies a flowering-time QTL on chromosome 12 in Scarlet eggplant overlapping with *SP5G*. **(f)** Diurnal expression analysis of *SaetSP5G* in Scarlet eggplants showing reduced expression of *SaelSP5G^Shum^.* Grey marks night periods. Cotyledons from three different (n=3) plants were used as biological replicates at each time point. TPM, Transcript Per Million. Data points represent means ±s.e. ZT 0 (Zeitgeber time) denotes the beginning of the light phase. **(g)** *SaetSP5G*^Shum^ co-segregates with early flowering in an F2 population. Significance (*p* < 0.05) determined by one-way ANOVA with post-hoc Tukey–Kramer HSD test. **(h)** Flowering time of *SaetSP5G*^Gilo^ and CRISPR-Cas9 engineered *Saetsp5g^cr^*^-Gilo^ alleles. Schematic of *SaetSP5G* is shown above. Red rectangle on the first exon denotes the deletion mutation (−13 bp). Means in (d), (g), (h) are shown above box plots. Sample size (n) in (d), (g), (h) represents the number of plants quantified per genotype. A two-tailed, two-sample t-test was performed to compare means in (c), (f), (h), asterisks indicate significant difference (\**p* < 0.05, \*\*\**p* < 0.01). Box plots in (c), (d), (g), (h) show the 25th, 50th (median) and 75th percentiles.

The Scarlet eggplant comprises diverse cultivar groups selected for edible fruit (Gilo), edible leaves (Shum), or ornamentation (Aculeatum)^48^. Gilo flowers substantially later than the Shum (mean 11.3 versus 5.3 leaves) (**Fig. 1b,c**), suggesting that accelerated flowering and fruiting were not selected in Gilo, possibly reflecting limited systematic breeding. We performed QTL-seq analysis on an F2 population between Gilo and Shum (**Supplementary Table 6**). As in Gboma, we identified two QTLs on chromosomes 4 and 12, the latter overlapping the orthologous *SP5G* locus(**Fig. 4e, Extended Data Fig. 4c**). Due to the absence of conspicuous causal variants, we performed diurnal expression analysis and found that the Shum *SP5G* allele (*SaetSP5G*^Shum^) was substantially downregulated relative to Gilo (*SaetSP5G*^Gilo^, Solaet3_12g005050) (**Fig. 4f**). Co-segregation analysis supported the association between *SaetSP5G*^Shum^ and early flowering (mean 8.6 versus 12.9 leaves in *SaetSP5G*^Gilo^) (**Fig. 4g**). Importantly, the *SaetSP5G*^Shum^ allele could not fully account for the Shum phenotype (mean 8.6 leaves in F2 segregants versus 5.3 in parental accession), suggesting that the as-yet undefined QTL on chromosome 4 contributes to extremely early-flowering. To validate causality, we generated a CRISPR knockout of *SaetSP5G*^Gilo^ (*Saetsp5g^cr^*^-Gilo^) (**Supplementary Table 1)** and found that homozygous mutants flowered earlier than WT (mean 9.6 versus 11.5 leaves) (**Fig. 4h**). Allele-frequency analysis across 70 Scarlet eggplant accessions showed that *SaetSP5G*^Shum^ is enriched among early-flowering ornamental Aculeatum genotypes, where rapid flowering was likely favored for synchronous blooming (**Extended Data Fig. 4d**). QTL-seq analyses on F2 populations between Gilo and four Aculeatum accessions revealed, again, the same peak above the *SP5G* locus (**Extended Data Fig. 4e**). Co-segregation analyses supported the association between *SaetSP5G*^Shum^ and early flowering in all four Aculeatum accessions (**Extended Data Fig. 4f**).

### Natural *SP5G* variation in wild species

Our results thus far demonstrate that recurrent loss-of-function mutations in the neofunctionalized flowering antagonist *SP5G* underlie parallel flowering-time change across crops from South America (tomato), Asia (Brinjal eggplant), and Africa (Gboma eggplant), and strongly associated with rapid flowering in Scarlet eggplant. However, early flowering is not restricted to agricultural adaptations, having also evolved repeatedly in wild species to escape deleterious environmental conditions^49^. Consistently, several wild Solanaceae flower rapidly, including the *S. prinophyllum*, which grows in arid regions of the Australian desert^44^ (**Fig. 1a,b**). We therefore asked whether allelic variation in *SP5G* contributed to flowering-time variation more broadly, possibly shaping wild adaptation. We focused on four wild species from three lineages varying in flowering time, together capturing 50 million years of evolution^50^. This panel included the late-flowering progenitor of tomato, *S. pimpinellifolium* (mean 14.6 leaves), as reference; the Australian nightshades, *S. cleistogamum* (desert raisin) and its close relative *S. prinophyllum* (mean 8.8 and 5.8 leaves, respectively) from the prickly eggplant lineage^21^ and *Physalis grisea* (groundcherry), a rapid-flowering species from genus *Physalis* (mean 6.1 leaves)^51^ (**Fig. 5a**).

**Figure 5.**
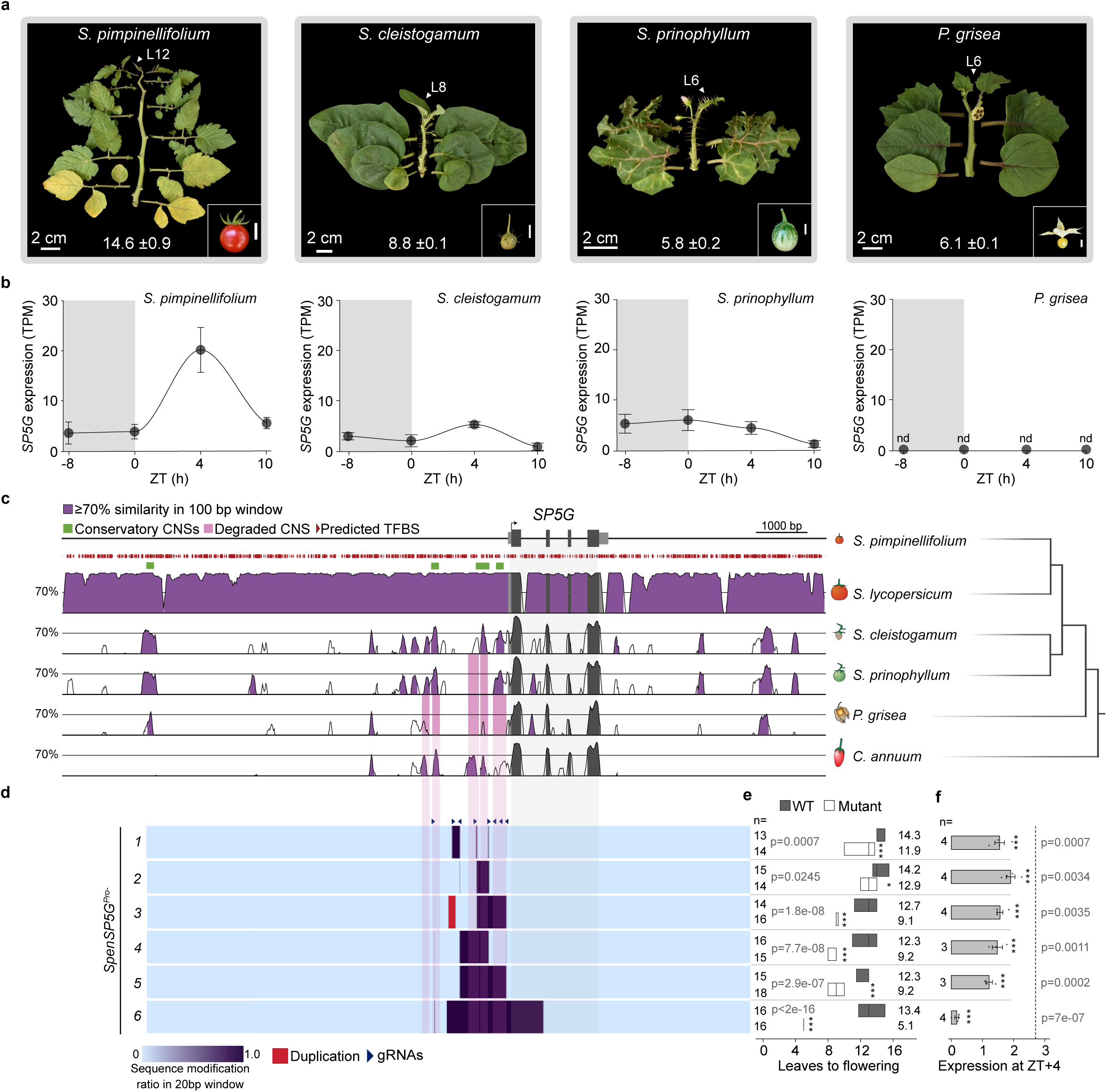
*SP5G cis*-regulatory variation is associated with early-flowering wild Solanaceae species. **(a)** Shoots and fruits of wild Solanaceae and their flowering time. Means ±s.e are presented. Scale bars: 2 cm (shoots) 1 cm (fruits). **(b)** Diurnal expression of *SP5G* orthologues in wild species. Data points present means ±s.e. Cotyledons from different plants (n=3) were used as biological replicates for each time point. nd, not detected. TPM, Transcript Per Million. **(c)** mVISTA alignments of *SP5G* orthologues and flanking *cis*-regulatory sequences. Above is the *SpimSP5G* gene model and 8.7-kb upstream and 4-kb downstream regions. Schematic phylogenetic tree is shown to the right. Conservation (purple and dark grey) was calculated with >70% similarity in 100-bp windows. TFBSs were predicted by FIMO^67^. Green rectangles denote conserved non-coding sequences (CNSs) found by Conservatory^53^. Light and gray regions denote UTRs and CDS, respectively. Pink vertical windows mark degraded CNSs. **(d)** Heatmaps of CRISPR-Cas9 *SpenSP5G^Pro^* alleles. Alleles were encoded; perturbations (purple intensity) are represented as the degree of sequence modification relative to *SpenSP5G* within 20-bp windows. Grey and pink windows match the syntenic regions shown in (d). **(e)** Flowering times of six engineered *SpenSP5G^Pro^ cis*-regulatory mutants relative to WT alleles from F2 segregating populations. Means are shown to the right. Box plots show the 25th, 50th (median) and 75th percentiles. **(f)** RT-qPCR expression analysis of the *SpenSP5G^Pro^* mutants in cotyledons 4h after sunrise. Vertical dashed line marks WT expression (*SpenSP5G*). 3-4 cotyledons from different plants were used as biological replicates. Bar charts are presented as means ±s.e. A two-tailed, two-sample t-test was performed to compare means of WT and mutant plants in (e), (f), asterisks indicate significant difference (\**p* < 0.05, \*\*\**p* < 0.01). ns, not significant. Sample size (n) in (e) represents the number of plants quantified per genotype.

Due to limited available intraspecific diversity, conventional forward genetics was not feasible. Importantly, none of these wild Solanaceae carry obvious deleterious coding mutations in *SP5G*^22^ (**Supplementary Fig. 3a**), suggesting that if *SP5G* natural variants are involved in flowering-time change, this variation likely arose through expression change mediated by *cis*-regulatory modifications. We therefore performed a comparative diurnal expression analysis. Consistent with previous findings, *SP5G* exhibited a strong diurnal pattern in cotyledons of *S. pimpinellifolium* (*SpimSP5G*, Solyc05g053850) (**Fig. 5b**)^24^. In contrast, diurnal behavior was partially retained in *S. cleistogamum* (*ScleSP5G*, Solcle2_12g004780), but a daytime expression peak was substantially reduced (**Fig. 5b**), mirroring the reduced expression in domesticated tomato carrying the *cis*-regulatory deletions (**Fig. 2c**). Strikingly, *SP5G* expression was low and diurnal behavior was lost in *S. prinophyllum* (*SpriSP5G*, Solpri1_12g005000), and *SP5G* expression was not detected in *P. grisea* (*PgriSP5G*, Phygri_05g017570) (**Fig. 5b**).

The reduced expression peak of *SP5G* and accelerated flowering in wild species implied functional association. We further tested this possibility by engineering CRISPR knockouts of *SP5G* in each species (**Extended Data Fig. 5a, Supplementary Table. 1**), reasoning that null mutations would have a mild effect in *S. cleistogamum* and weak or effect in *S. prinophyllum* and *P. grisea*. As predicted, *SP5G* knockout in *S. pimpinellifolium* substantially accelerated flowering (mean 7.9 versus 14.6 leaves in WT), whereas knockouts in *S. cleistogamum*, *S. prinophyllum*, and *P. grisea* had little to no effect (**Extended Data Fig. 5b,c**). Together, the progressive reduction in *SP5G* expression levels and corresponding rapid flowering, along with diminishing effects of *SP5G* knockouts in rapid-flowering species, supported a plausible broader mechanism by which *cis*-regulatory changes in *SP5G* shape flowering-time change across both wild and domesticated Solanaceae. Whether *SP5G* loss-of-function alleles are adaptive in wild taxa deserves further investigation through establishing population-scale genomics for formal tests of selection^37^.

To further search for *cis*-regulatory modifications that may link *SP5G* expression changes and accelerated flowering in wild species, we aligned the *cis*-regulatory regions surrounding *SP5G* across 22 Solanaceae with mVISTA^52^, using the functional *SpimSP5G* as a reference. We identified several upstream and downstream regions that were deeply conserved from a database of conserved non-coding sequences (CNSs) identified by our algorithm, Conservatory^53^ (**Fig. 5c, Extended Data Fig. 6**). Several CNSs located in the *SP5G* promoter region were degraded in the early-flowering species *S. prinophyllum* and *P. grisea* (**Fig. 5c, Extended Data Fig. 6**). Because the CNSs within this region, which spans ∼1.6 kb beginning ∼50-bp upstream of the TSS, were naturally degraded or lost in these early-flowering species, a direct *in vivo* assessment of their control of *SP5G* expression and flowering time was not possible. We therefore again leveraged the tomato *SpenSP5G* introgression line, in which these CNSs are intact, to test their functional relevance. Using a multiplex 8-guide RNA (gRNA) editing construct, we isolated six alleles carrying either small or large deletions (**Supplementary Table 1**) that partially or fully eliminated CNSs naturally degraded in *S. prinophyllum* and *P. grisea*.

Overall, homozygous mutants of all six *SpenSP5G^Pro^* alleles exhibited reduced *SP5G* expression and accelerated flowering, indicating these CNSs within the promoter region harbor functional *cis*-regulatory elements (CREs). The strongest allele (*SpenSP5G^Pro-6^*; 2.45-kb deletion) flowered after five leaves and showed only residual *SP5G* expression due to loss of the first exon (**Fig. 5d,e**). Three promoter alleles carrying large deletions (*SpenSP5G^Pro-3^*; 754 bp, *SpenSP5G^Pro-4^*; 734 bp, *SpenSP5G^Pro-5^*; 1178 bp), which eliminated multiple proximal CNSs and introduced indels in a more distal CNS (**Fig. 5f, Extended Data Fig. 6**), showed substantial reductions in both *SP5G* expression and flowering time (mean 9.1, 9.2, and 9.2 leaves versus 12.7, 12.3, and 12.3 in WT) (**Fig. 5e,f, Extended Data Fig. 7**). These observations are consistent with previous findings that large or cumulative smaller CNS perturbations are often required for substantial phenotypic effects^31^. In support, two smaller deletion alleles (*SpenSP5G^Pro-1^*and *SpenSP5G^Pro-2^*), which disrupted overlapping but fewer CNSs, caused much less substantial reductions in flowering time (mean 11.9 and 12.9 leaves versus 14.3 and 14.2 in WT, respectively) (**Fig. 5d-f, Extended Data Fig. 7**).

Notably, none of the engineered promoter alleles fully recapitulated the early-flowering phenotypes observed in *S. prinophyllum*, *P. grisea* (**Fig. 5a**), suggesting additional genetic changes outside this labile region contributed additively or synergistically to accelerated flowering in each species. These observations suggest that substantial phenotypic changes from *cis*-regulatory variation of *SP5G* depends on more than one variant affecting CNSs. This parallels our findings in tomato (**Fig. 2**) and aligns with previous *cis*-regulatory editing results^31^ of multi-step accumulation of cryptic or weak *cis*-regulatory mutations as a common path to elicit substantial phenotypic change. In contrast, in Brinjal and Gboma eggplants single large structural variants drove similar phenotypic change, raising an old debate in evolutionary genetics: how similar selective pressures acting on distinct regulatory-network architectures influence whether phenotypic evolution proceeds through multiple small-effect versus few large-effect mutations^54^.

## DISCUSSION

Here, through the power of pan-genetics in the Solanaceae, we show that recurrent mutation of a neofunctionalized flowering antagonist contributed repeatedly to phenotypic change across 50 million years of evolution (**Extended Data Fig. 8**). Although limited statistical power restricts our ability to definitively infer positive selection from independent population-genetic tests (**Supplementary Notes, Supplementary Fig. 1 and 2**)^37^, multiple lines of evidence from forward and reverse genetics across crops and wild species provide compelling support for adaptive significance of *SP5G* loss-of-function alleles. Our findings extend classical neofunctionalization theory^2^ by suggesting that the emergence of network-embedded neofunctionalized antagonistic paralogues can bias evolutionary trajectories by channeling repeated evolution through parallel genetic routes. This framework reveals an apparent paradox: while regulatory network capacity expands through gene duplication, selection may preferentially target the same emerging paralogue for future adaptive change. This, however, is consistent with natural selection acting along genetic paths of least resistance^55^, among which the accessibility of loss-of-function mutations may represent a primary route^56^. Accordingly, the recurrent attenuation of the flowering repressor *SP5G* may exemplify how adaptive evolution can proceed through loss-of-function mutations in repressive rather than promoting counterparts of regulatory networks.

However, genes and their variants, including those derived from modifications of *SP5G*, do not act in isolation. Uncovering the broader regulatory landscape of quantitative trait variation and evolution is a central objective of genotype-phenotype prediction and crop breeding^57^. Flowering time is an inherently complex trait regulated by polygenic interactions^19^. In tomato, mutation in the ancestral *antiflorigen SP* was discovered a century ago and revolutionized the breeding and productivity of tomatoes^24,58^. While *SP5G* represses the initial flowering event of primary and side shoots, the transformative *sp-classic* mutation accelerated flowering on later forming sympodial shoots, resulting in compact plants with synchronous flowering and fruiting that enabled large-scale field production. Importantly, the agricultural benefit from *sp-classic* is effective only in the background of the compromised *SlycSP5G* allele^24^ (**Fig. 2**), highlighting how founder contingency mutations can influence the effects of subsequent variation in paralogous network components that can themselves become contingencies in later steps of crop improvement. It is noteworthy in this respect that the yet to be characterized QTLs on chromosomes 4 and 12 in African and Brinjal eggplants (**Fig. 3c, Extended Data Fig. 4a,c**) can reveal how evolutionary changes in flowering time across Solanaceae reflect both repeated targeting of *SP5G* and species-specific subsidiary contingencies whose emergence and effects are likely shaped by broader constraints of network-level architecture. Identifying such network-level contingencies will improve our understanding of polygenic adaptation and accelerate the translation of such knowledge into predictable engineering of agricultural traits, particularly in understudied indigenous crops.

More broadly, an enduring question in evolutionary genetics concerns what enables duplicated genes to persist once a paralogue emerges, and how natural selection shapes the functional relationships of retained paralogues^1,59^. An extension of this long-standing question emerging from our study is how frequently emerging paralogues neofunctionalize and evolve opposing functions from their ancestral genes, and whether such antagonistic configurations affect subsequent adaptive evolution. To assess the generality of this concept, we revisited the literature and found examples with similar patterns of paralogue neofunctionalization and subsequent channeling of phenotypic change and adaptation across diverse biological systems. Beyond *SP5G* in the Solanaceae, equivalent cases of neofunctionalized *antiflorigens* shaped flowering time in sugar beet, onion, and eelgrass^25–27^. Extending beyond the *florigen-antiflorigen* network, a striking parallel example is found in the Brassicaceae family. A subclade of the flowering-promoting MADS-box gene family evolved flowering-repressing activity early in Brassicaceae diversification^60^, and allelic variation in a specific member of this clade (*FLOWERING LOCUS C*) underlies repeated flowering-time change across lineages, including ecotypes of *Arabidopsis thaliana*^3^ (**Supplementary Discussion**). Similar patterns of antagonistic neofunctionalization appear beyond flowering; independently evolved opposing paralogues of the classical maize domestication gene (*TEOSINTE BRANCHED1*) drove changes in plant architecture in potato and rice, spanning 120 million years^61,62^. These and multiple other compelling examples stretching across kingdoms, including leaf complexity in plants, wing development in insects, and evolution of the human neocortex (**Supplementary Discussion**), support the view that the emergence of opposing functions among paralogues, by giving rise to antagonistic and quantitatively tunable architectures in gene regulatory networks^63^, may represent a broader principle in phenotypic and adaptive evolution. This framework is now becoming an empirically tractable question: to what extent have neofunctionalized paralogues, and antagonizing paralogues in particular, created evolutionary contingencies that channeled subsequent phenotypic and adaptive evolution? As illustrated in this study, continued integration of comparative and population genomics, phylogenetic reconstructions, and functional genetic analyses across diverse traits and taxa will reveal the broader scope of this synthesis.

## Supporting information

SI Guide

## METHODS

### Plant material, growth conditions and phenotyping

Seeds of the introgression line IL5-4 (cv. M82) which carries a functional *SP5G* variant from *S. pennellii* (*SpenSP5G*) were obtained from the Charles M. Rick Tomato Genetics Resource Center (TGRC) at the University of California, Davis. IL5-4 and *S. pimpinellifolium* (accession PAS14479) were used as WT background for CRISPR-Cas9 tomato mutagenesis experiments. For mutagenesis in related Solanaceae plants, we used reference backgrounds as described in Satterlee et al., (2024)^44^. Seeds were sown and grown in 96-well flats for 3-4 weeks before transplanting to 4L pots and grown under greenhouse conditions. The greenhouse operates under long-day conditions (16h light, 8h dark) with natural and artificial light (high pressure sodium bulbs ∼250 µmol/m2), at a temperature between 26-28 C° (day) and 18-20 C° (night), and relative humidity 40-60%. Field-grown plants were drip-irrigated and fertilized using standard regime. Quantification of flowering time in WT and homozygous mutant plants was done under long-day conditions (16h light, 8h dark) and was defined as the number of leaves produced before the emergence of the first inflorescence on the main shoot, following a standard approach as a proxy for measuring flowering time. Raw counts of flowering time are available in **Supplementary Table 3.**

### Distribution maps and species status

Species were categorized into wild, indigenous, and domesticated, based on taxonomic literature and expert opinion^21^ (PBI Solanum Project (2024), Solanaceae Source; http://www.solanaceaesource.org/). The distribution map was generated using the open-source osm-liberty package (http://github.com/maputnik/osm-liberty/). Native ranges were derived from the same taxonomic literature and approximate centroids of the ranges were used for the mapping. The map is from osm-liberty, designed for open-source maps.

### CRISPR-Cas9 mutagenesis, plant transformation, and selection of mutant alleles

Generation of transgenic tomato and Solanaceae plants with CRISPR-Cas9 mutagenesis was performed as previously described^64^. Briefly, gRNAs were designed with Geneious Prime (https://www.geneious.com). The Golden Gate assembly method was used to clone gRNAs into a binary vector with Cas9 and the neomycinphosphotransferase II selectable marker gene^65^. Binary vectors were introduced into tomato cotyledons using *Agrobacterium*-mediated transformation^66^. Transgenic plants were screened for mutations using PCR primers surrounding the gRNA target sites. PCR products were screened for obvious shifts in size by gel electrophoresis, and mutations were defined by Sanger sequencing. First or second generation transgenics (T0 or T1) were backcrossed to WT to eliminate the Cas9 transgene and potential off-target mutations. F2 populations derived from these crosses were used for phenotypic analyses. All gRNA and primer sequences are listed in **Supplementary Table 7.**

### *Cis*-regulatory sequence conservation analysis, identification of open chromatin regions, and prediction of TFBSs

The Solanaceae family conservation analysis was performed using the ‘Conservatory’ algorithm^53^ to predict conserved non-coding sequences within the 5’ and 3’ regions of *SpimSP5G* that were shared among other Solanaceae species. The closest *SP5G* orthologue from each species was determined based on micro-synteny and highest protein sequence similarity to *SlycSP5G* (cv. M82). The non-coding sequence of each *SP5G* orthologue was extracted between its flanking genes and aligned to *SpimSP5G* (PAS014779) using mVISTA LAGAN (http://genome.lbl.gov/vista/mvista/submit.shtml)^52^. Conservation was calculated in 100-bp windows, with a 70% similarity threshold. TFBSs were predicted by scanning the *SpenSP5G* (IL5-4) 5’ and 3’ regions for motifs using FIMO in the MEME suite (http://meme-suite.org/doc/fimo.html)^67^. Position frequency matrices for known plant transcription factors were obtained from the JASPAR CORE PFMs of plants collection 2022^68^ using a *p-value* cutoff of 0.00001 to predict TFBSs.

### Genome assemblies and gene annotations

We assembled and annotated the genomes following protocols we developed for our *Solanum* pan-genome^22^. Genome size and heterozygosity were estimated from 21-bp k-mer profiles derived from PacBio HiFi sequencing reads using KMC3 v3.2.1^69^, followed by model fitting in GenomeScope 2.0^70^. De novo assemblies were generated from PacBio HiFi reads using hifiasm^71^ with parameter −l=3. Assembled contigs were screened to remove potential bacterial, fungal, mitochondrial and chloroplast sequences. Screened contigs were subsequently scaffolded using RagTag^32^, guided by the closest available reference genome to establish chromosomal ordering and orientation. Assembly completeness was evaluated using BUSCO v5^72^ against the *Solanales_odb10* dataset. Assembly continuity was assessed through N-chart analyses implemented in R using ggplot2, adapted from the Nchart framework. (https://github.com/MariaNattestad/Nchart). For genome annotation, orthologous gene models were transferred from a reference genome of *S. macrocarpon* (accession PI 441914)^44^ and from the reference *S. melongena* (accession GPE001970)^46^ using LiftOn^73^, retaining models with ≥50% alignment coverage and ≥75% sequence identity. Gene space completeness was independently assessed using BUSCO v5^72^.

### Phylogenetic analyses

Homologous proteins were identified using blastp and a profile HMM–based approach followed by phylogenetic analysis. Known orthologues were aligned with MAFFT (v7.505)^74^ and used to build an HMM profile with HMMER (v3.3.2). This profile was searched against target proteomes using hmmsearch, and significant hits (E < 1 × 10^-5^) were extracted and retrieved with seqtk. Candidate homologs were realigned with MAFFT^74^ using the default parameters, and maximum-likelihood phylogenetic trees were inferred using IQ-TREE (v2.2.2)^75^ with automatic model selection and 1,000 ultrafast bootstrap replicates. The trees were visualized in R with the ggtree package (v4.0.3)^76^.

### Variant calling and selective sweep analyses

For tomato, the dataset was obtained from the authors as a VCF file^35^. For eggplant variant calling, paired-end reads from 374 accessions were trimmed for poly-G/poly-A sequence and adapter contamination using Trimmomatic v0.41^77^ and aligned to the *S. melongena* reference genome GPE001970^46^ using miniBWA v0.3^78^. Alignments with median depth ≥10 were retained for variant calling, excluding four low-depth accessions. Variants were jointly called with FreeBayes v1.3.9^79^ using a minimum base quality of 30, normalized relative to the reference, split into biallelic records, and filtered to remove variants with QUAL < 30 using BCFtools v1.23.1^80^. Mapping and variant-calling quality were assessed with MultiQC^81^ and BCFtools stats^80^.

To test for signatures of selection at *SP5G*, related paralogues, and other domestication loci, statistics were calculated in 100-kb windows with 10-kb steps and included nucleotide diversity (π), diversity ratios, Tajima’s D, Weir–Cockerham and Hudson *F_ST_*, and haplotype-based nSL and XP-nSL. Focal windows were ranked against the corresponding genome-wide distribution for each statistic, dataset, and population comparison. We also performed simulations under published tomato demographic models^34^ to evaluate power to detect sweeps under different starting allele frequencies, selection coefficients, recombination rates, sample panels, and sweep ages. Full analysis details are provided in the **Supplementary Notes**.

### Panagram analysis

Panagram (https://github.com/kjenike/panagram), an alignment-free pan-genome analysis tool, was used to identify deletions in *SmelSP5G* across a panel of 341 Brinjal eggplant accessions. Briefly, the software constructs a pan-genome-wide k-mer index by counting k-mers in each genome using KMC3^69^ and encoding their presence or absence in bit vectors. These vectors are then merged into a unified bitmap that tracks which genomes contain each k-mer. The resulting pan-k-mer bitmap is anchored to a reference genome to enable visualization of k-mer presence or absence relative to the reference coordinates. For this study, the bitmap was anchored on the reference accession GPE001970^46^. For each genome, k-mer similarity to the reference was calculated in 1-Mbp bins across each chromosome by dividing the number of k-mers matching the reference by the total number of k-mers in the bin, yielding values between 0 and 1 for each bin, for each accession.

Analogous to the widely used algorithm Mash^82^, the binned k-mer similarities were used to construct a dendrogram estimating the phylogenetic relationships among the 346 eggplants. Pairwise Euclidean distances between accessions were calculated using the binned k-mer similarities from all chromosomes for each accession as input. These distances were used to group the accessions using complete-linkage hierarchical clustering. Finally, to examine variation at finer resolution, a zoomed-in pan-k-mer bitmap using 500-bp bins was generated around the *SmelSP5G* locus. At this scale, deletions were visible as regions with reduced similarity or a complete lack of k-mers shared with the reference genome. Specifically, bins with less than 50% k-mer similarity were found in 10 accessions, indicating candidate deletion events at the locus.

### GWAS in Brinjal eggplant

The GWAS was performed on 283 lines derived from the first multiparent advanced generation intercross (MAGIC) population that were generated from seven *S. melongena* accessions and one wild *S. incanum*, as previously described^47^. They were genotyped through low-coverage whole-genome sequencing (lcWGS) at 3X depth (3.6 Gb of clean data per sample) using the DNBseq platform at Beijing Genomics Institute (BGI Genomics, Hong Kong, China) as part of a previous project^83^. High-quality reads, as determined by the fastq-mcf software v. 1.04.676^84^, were aligned to the *S. melongena* genome (accession H15) using the BWA-MEM algorithm v. 0.7.17–r1188^85^ with default parameters. Plants were grown under two different environmental conditions: (1) under natural conditions in an open field during the spring-summer season in Alcasser, Valencia, Spain (39°23’40.2”N, 0°26’54.4”W), where temperatures ranged from 11 to 32 °C, with a photoperiod of approximately 11-15 hours of daylight, and (2) under conventional greenhouse conditions during the autumn-winter season in Ruescas, Almería, Spain (36°49’14.7”N, 2°14’08.1”W), with temperatures ranging from 10 to 28 °C, and a photoperiod of approximately 9-11 hours of daylight. In both experiments, a randomized complete block design was used, with three blocks and one plant per block for each line. Genotypic and flowering time phenotypic data were used to perform a GWAS analysis using GAPIT software v.3^86^ under the mixed linear model (MLM). The result was visualized as a Manhattan plot using the qqman R package^87^, and multiple testing correction was applied using the Bonferroni method^88^ with a significance threshold of 0.05.

### Mapping of loci underlying flowering time variation in African eggplants

The late-flowering parents of Gboma (*S. dasyphyllum*, wild progenitor) and Scarlet (Gilo, accession PI 424860) eggplants were crossed to early flowering Gboma (*S. macrocarpon,* accessions PI 441914 and SC129) and Scarlet (Shum, accession 804750187; Aculeatum, accessions SC102, PI 636107, PI 666076, PI 666078) parents, respectively, to map the QTLs and the causative variants affecting flowering time. The resulting F1 progenies were selfed to generate F2 mapping populations, which were grown in a field site at Lloyd Harbor, New York, USA. Flowering time was quantified by counting the number of leaves produced before the emergence of the first inflorescence in F2 individuals. Individual plants were phenotyped from each segregating F2 populations (**Supplementary Table 6**). For each population, DNA from 30 random individuals at the low and high ends of the phenotypic distribution were pooled for bulk-segregant QTL-seq analysis. DNA was extracted from young leaf tissue using the DNeasy Plant Pro Kit (Qiagen) according to the manufacturer’s instructions. Briefly, leaf tissue was ground using a SPEX SamplePrep 2010 Geno/Grinder^TM^ (Cole-Parmer, NJ, USA) for 2 min at 1,440 rpm. The sample DNA (1 µl assay volume) concentrations were assayed using Qubit 1× dsDNA HS buffer (Fisher Scientific, MA, USA) on the Qubit 4 fluorometer (Fisher Scientific, MA, USA) according to the manufacturer’s instructions. The bulked late-flowering F2 individuals and the bulked early-flowering F2 individuals, with an equivalent mass of DNA pooled from each individual plant, were combined to yield a final pooled mass of 3 µg in each bulk. Pooled DNA was purified using x1.8 volume of AMPure XP beads (Beckman Coulter) and DNA concentration was analyzed using NanoDrop One spectrophotometer (Fisher Scientific, MA, USA). Paired-end sequencing libraries for QTL-seq analysis were prepared with >1 µg of DNA using the KAPA HyperPrep PCR-free kit (Roche) according to the manufacturer’s instructions. Indexed libraries were pooled for sequencing on a NextSeq 2000 P3 chip (Illumina). Mapping was performed using the end-to-end pipeline implemented in the QTL-seq software package^89,90^ (v.2.2.4, https://github.com/YuSugihara/QTL-seq) with reads aligned against the *S. macrocarpon* (accession PI 441914) or *S. aethiopicum* (accession PI 424860) reference genome assemblies^44^.

### Co-segregation analyses

Co-segregation analysis of the *SP5G* 83-bp deletion allele in tomato was performed on an F2 population derived from an intraspecific cross between the *S. pimpinellifolium* accessions LA1237 (intact 83 bp) and LA1589 (deleted 83 bp). Cotyledon tissue was collected and DNA was extracted using the cetrimonium bromide (CTAB) method^91^. Primers were designed to amplify the 83-bp element and its flanking sequences. PCR reactions were prepared in a 10 µL reaction using KOD One PCR Master Mix (Toyobo, Osaka, Japan). Genotypes were determined by separation of the visualized PCR products using gel electrophoresis. Co-segregation analysis of the *SP5G* 52-bp deletion allele in tomato was performed on an F2 population derived from an interspecific cross between the *S. pimpinellifolium* accession LA1589 (intact 52 bp) and cultivated tomato cv. M82 (deleted 52 bp), as described above for the 83-bp allele.

Co-segregation analysis of the coding deletion allele in Brinjal eggplant was performed on the selfed progeny of an advanced backcross individual (BC4S1) derived from an interspecific cross between the wild relative *S. elaeagnifolium* and a domesticated eggplant (accession MEL3), which carry the intact and the deletion allele, respectively. Genotyping was based on sequencing data generated using the 5k eggplant Single Primer Enrichment Technology (SPET) platform^92^. The selected BC4 individual carried a heterozygous *S. elaeagnifolium* introgression within the Brinjal genetic background, which specifically spanned the *SP5G* genomic region. Co-segregation analysis was subsequently conducted in the BC4S1 segregating population obtained by self-pollination of this individual. Leaf tissue was collected and DNA was extracted using the silica matrix extraction (SILEX) method^93^. Primers were designed to amplify the 2.15-kb coding deletion and its flanking sequences. PCR reactions were prepared in a 10 µL reaction using 2x PCRBIO Taq mix (PCR Biosystems Ltd, London, England). Genotypes were determined by separation of the visualized PCR products using gel electrophoresis.

Co-segregation analysis of the transposon-inserted *SmacSP5G* allele in Gboma eggplant was performed on an F2 population derived from interspecific crosses between the wild progenitor *S. dasyphyllum* and the Gboma accessions PI 441914 and SC129, as described above for the tomato 83-bp allele.

Co-segregation analysis of the *SP5G* alleles in Scarlet eggplant was performed on F2 populations derived from intraspecific crosses between the Gilo accession PI 424860 and Shum (accession 804750187) or Aculeatum varieties (accessions SC102, PI 636107, PI 666076, PI 666078), using Cleaved Amplified Polymorphic Sequence (CAPS) located 400-bp upstream of *SaetSP5G* in a T-to-C SNP modified Ndel restriction site (New England BioLabs, MA, USA). The same CAPS marker was used to analyze *SaetSP5G* allele frequency across our Scarlet eggplant germplasm. All primers used for co-segregation analyses are available in **Supplementary Table 7**.

### Tissue collection and RNA extraction and quantification

Fully expanded cotyledons were collected for 3-4 biological replicates from different plants grown under long-day conditions (16h light, 8h dark) and were flash-frozen in liquid nitrogen in 2 ml microfuge tubes containing two 1/8 inch (∼3.175 mm) 440 stainless steel ball bearings (BC Precision, TN, USA). Tubes containing tissue were placed in a −80°C tube rack and ground using a SPEX^TM^ SamplePrep 2010 Geno/Grinder^TM^ (Cole-Parmer, NJ, USA) for 30 seconds at 1,440 rpm. Total RNA was extracted using Quick-RNA MicroPrep Kit (Zymo Research). RNA was treated with DNase I (Zymo Research, CA, USA) according to the manufacturer’s instructions. RNA concentration and quality was assessed using a NanoDrop One spectrophotometer (Fisher Scientific, MA, USA).

### Gene expression analysis using RT-qPCR

Between 500 and 2000 ng of RNA were used for complementary DNA synthesis with the SuperScript IV VILO Master Mix (Invitrogen) according to the manufacturer’s instructions. Quantitative RT-PCR (RT-qPCR) in Gboma eggplant (*SmacSP5G* and *SdasSP5G*), *S. pimpinellifolium*, and landrace or cultivated tomato was performed with gene-specific primers using Fast SYBR^TM^ Green Master Mix reaction (Applied Biosystems by Fisher Scientific, MA, USA). As endogenous control, the Gboma and tomato housekeeping gene Ubiquitin (Solmac3_01g019950 and Solyc01g056940, respectively) was used for normalization. RT-qPCR in tomato (*SpenSP5G* and *SlycSP5G*) was performed using Taqman^TM^ technology with allele-specific custom probes and TaqMan™ Fast Advanced Master Mix (Fisher Scientific, MA, USA). Both analyses were performed on the QuantStudio 7 system (Fisher Scientific, MA, USA). Primers and Taqman^TM^ probe information are available in **Supplementary Table 7**.

### Diurnal expression analysis of *SP5G* in wild Solanaceae using 3’ RNA-sequencing

Tissue collection and RNA extraction were performed as described above. The 3′ RNA-seq QuantSeq FWD V2 kit (Lexogen, Vienna, Austria) was used to generate RNA libraries. Libraries were sequenced on an Illumina NovaSeq X (Illumina, CA, USA), 10B lane, generating paired-end 150-bp reads. The read corresponding to the 3′ end of transcripts was used for downstream analyses. Transcriptomic alignment was performed using STAR v2.7.11b^94^. *S. pimpinellifolium* samples were aligned to the PAS014479_MAS2.0 genome^32^, while all other samples were aligned to their respective reference genomes previously described^22,44^. Following alignment, expression values were normalized using RNAnorm v2.1.0^95^. To assess RNA library quality and alignment consistency across species, genes with one-to-one orthologues present in all six species were used for principal component analysis (PCA). PCA was performed using Python v3.9 and scikit-learn v1.7.1^96^. RNA-sequencing was performed at Cornell Biotechnology Resource Center (RRID:SCR_021727).

### Statistical methods

Statistical tests for phenotypic and expression analyses were performed in R. Statistical significance was determined by using two-tailed t-test for a single comparison between two groups of treatments or genotypes, and one-way ANOVA for multiple comparisons with post-hoc Tukey–Kramer HSD test. *P-value* cutoff of <0.05 was used to indicate significant differences (* for *P* < 0.05, ** for *P* < 0.01, *** for *P* < 0.001). Different lowercase letters and asterisks indicate significant differences (*P* < 0.05). P-values of all phenotypic and expression statistical tests performed in this study are available in **Supplementary Table 8**.

## ONLINE CONTENT

### Reporting summary

Further information on research design is available in the Nature Portfolio Reporting Summary linked to this article.

### Data availability

All data are available within this manuscript and its Supplementary Information. Raw sequencing data are available at the SRA under BioProject PRJNA1457893. Genomes and gene annotations of *S. lycopersicum* (M82), *S. pimpinellifolium* (PAS014479_MAS2.0), *S. prinophyllum*, *S. cleistogamum*, *P. grisea*, *S. aethiopicum* (PI 424860 and 804750187), *S. macrocarpon* (accessions PI 441915 and SC129), *S. dasyphyllum*, *S. melongena* (accession H15), eggplant VCF files, including images of species and accessions are open access and available at the solpangenomics website (www.solpangenomics.com) and the Solanaceae Genomics Network (SGN: https://solgenomics.net/ftp/genomes/Solanum_pangenomics/). All source data for flowering-time quantifications and associated summary of statistical tests (p-values) are available in **Supplementary Tables 3 and 8**.

### Code availability

Panagram analysis scripts are available at GitHub (https://github.com/kjenike/panagram). Other analysis scripts were adapted from GitHub (https://github.com/pan-sol/pan-sol-pipelines).

## Acknowledgments

We thank present and former members of the Lippman laboratory and Yuval Eshed for discussions and input on the manuscript; Denise Tieman, Xiang Li, and Harry Klee for providing tomato VCF files; the Tomato Genetics Resource Center at UC-Davis for providing tomato seed stocks; S. Qiao, K. Schlecht, A. Harrison, and R. Ratchford for assistance with greenhouse and field plant care; This work was performed with assistance from CSHL Sequencing Technologies and Analysis Shared Resource. We thank S. Goodwin, S. Muller, R. Wappel, S. Mavruk-Eskipehlivan, and E. Ghiban for sequencing support. First and corresponding authors acknowledge using ChatGPT solely to improve grammar and readability of specific phrases and sentences. All authors reviewed and edited later drafts.

## Author contributions

Conceptualization: HS, MCS, ZBL

Data curation: HS, DC, AA, IG, AH, NLB, JLR, JWS, BF

Formal analysis: HS, DC, AA, IG, SR, AH, KMJ, NLB, LCM, NK, JLR, MJP, JWS, BF, VBF, JG, AS, JP, MCS, ZBL

Funding acquisition: JG, JVE, AS, JP, MCS, ZBL

Investigation: HS, DC, AA, IG, SR, AH, KMJ, NLB, JLR, MJP, JWS, BF, VBF, JG, AS, JP, MCS, ZBL

Methodology: HS, DC, AA, IG, SR, AH, KMJ, NLB, LCM, NK JLR, MJP, JWS, BF, VBF, GMR, BMS, JG, JVE, AS, JP, MCS, ZBL

Project administration: HS, JP, MCS, ZBL

Software: SR, AH, KMJ, NLB, LCM, NK, AS, MCS Resources: HS, DC, AA, AH, KMJ, NLB, JP, MCS, ZBL

Supervision: JG, JVE, AS, JP, MCS, ZBL

Validation: HS, AA, IG, SR, AH, JG, JVE, AS, JP, MCS, ZBL

Visualization: HS, DC, AA, IG, AH, KMJ, NLB, LCM, NK, VBF, MCS, ZBL

Writing-original draft: HS, MCS, ZBL

Writing-review and editing: HS, DC, AA, IG, AH, GMR, JG, JVE, AS, JP, MCS, ZBL

## Contact details

Correspondence and requests for materials should be addressed to Michael C. Schatz or Zachary B. Lippman.

## Additional information

The online version contains supplementary material available at https://doi.org/xxxxxx.

## EXTENDED DATA FIGURE LEGENDS

**Extended Data Figure 1.**
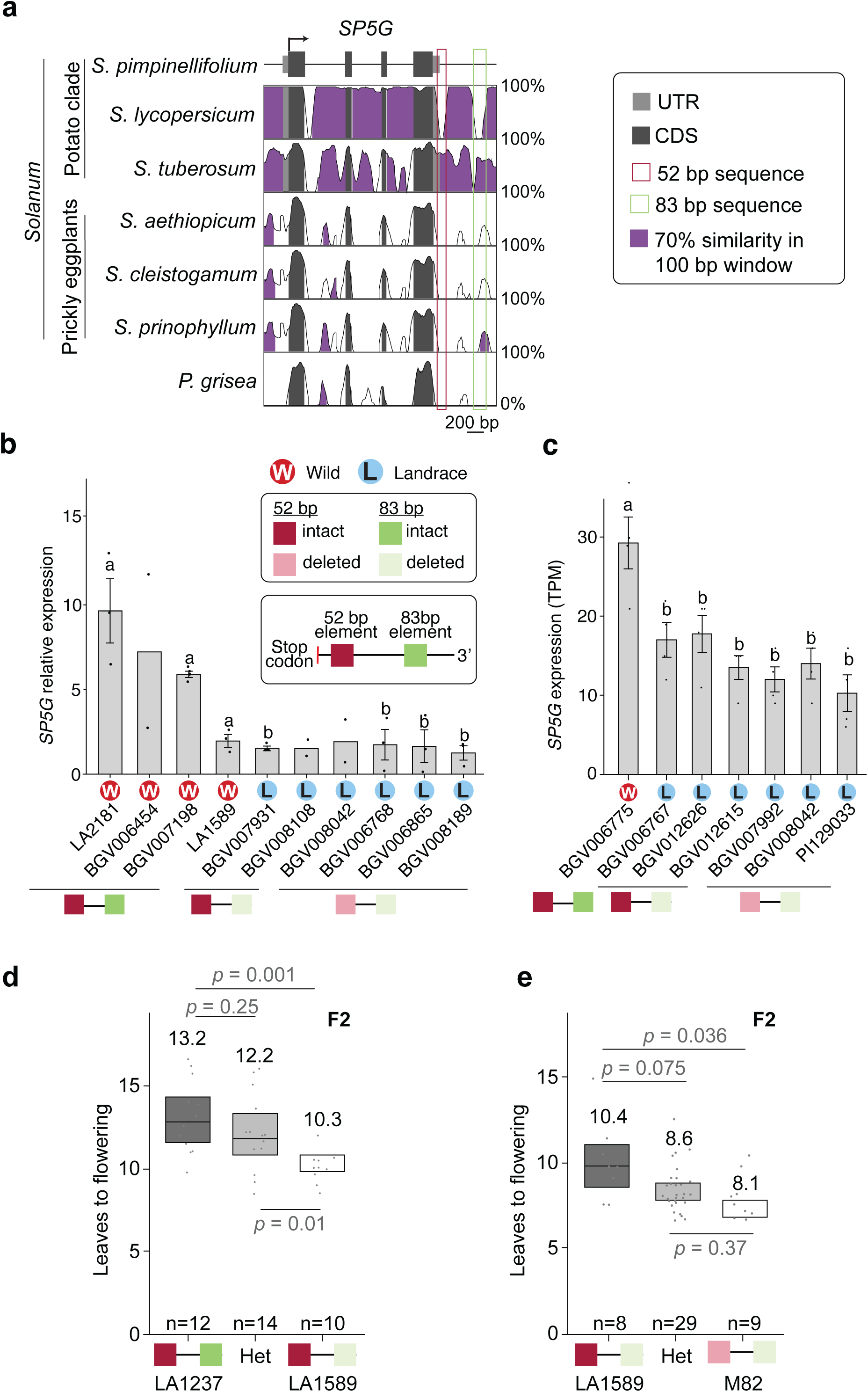
C*is*-regulatory deletions downstream of *SP5G* drove stepwise acceleration of flowering time during tomato domestication. **(a)** *SpimSP5G* gene model and proximal regulatory regions. mVISTA DNA sequence alignments of six *SP5G* orthologues using the *SpimSP5G* gene as reference. Conservation (purple) was calculated as sequences with 70% similarity in 100-bp windows. **(b)** RT-qPCR expression analysis of *SP5G* in cotyledons of *S. pimpinellifolium* and landrace accessions homozygous for either the intact or deleted 83-bp and 52-bp sequences. **(c)** *SP5G* expression in cotyledons of *S. pimpinellifolium* (accession BGV006775) and landrace accessions homozygous for either the intact or deleted 83-bp and 52-bp sequences. TPM, Transcript Per Million. Cotyledons (n=3) from different plants in (b) and (c) were used as biological replicates. Bar charts present means ±s.e. Expression values were obtained from transcriptome data published in Alonge et al., (2020)^32^. **(d)** Flowering time of tomato plants homozygous and heterozygous for either the intact or deleted 83-bp sequence from an F2 segregating population. **(e)** Flowering time of tomato plants homozygous and heterozygous for either the intact or deleted 52-bp sequence in an F2 segregating population. Dark red and green squares in (b), (c), (d), (e), denote intact sequence, light red and green squares denote deletion. Sample size (n) in (d), (e) represents the number of plants quantified per genotype. Lowercase letters in (b), (c) indicate significant difference (*p* < 0.05) based on one-way ANOVA with post-hoc Tukey-Kramer HSD test. Means in (d), (e) are shown above box plots. Box plots in (d), (e) show the 25th, 50th (median) and 75th percentiles.

**Extended Data Figure 2.**
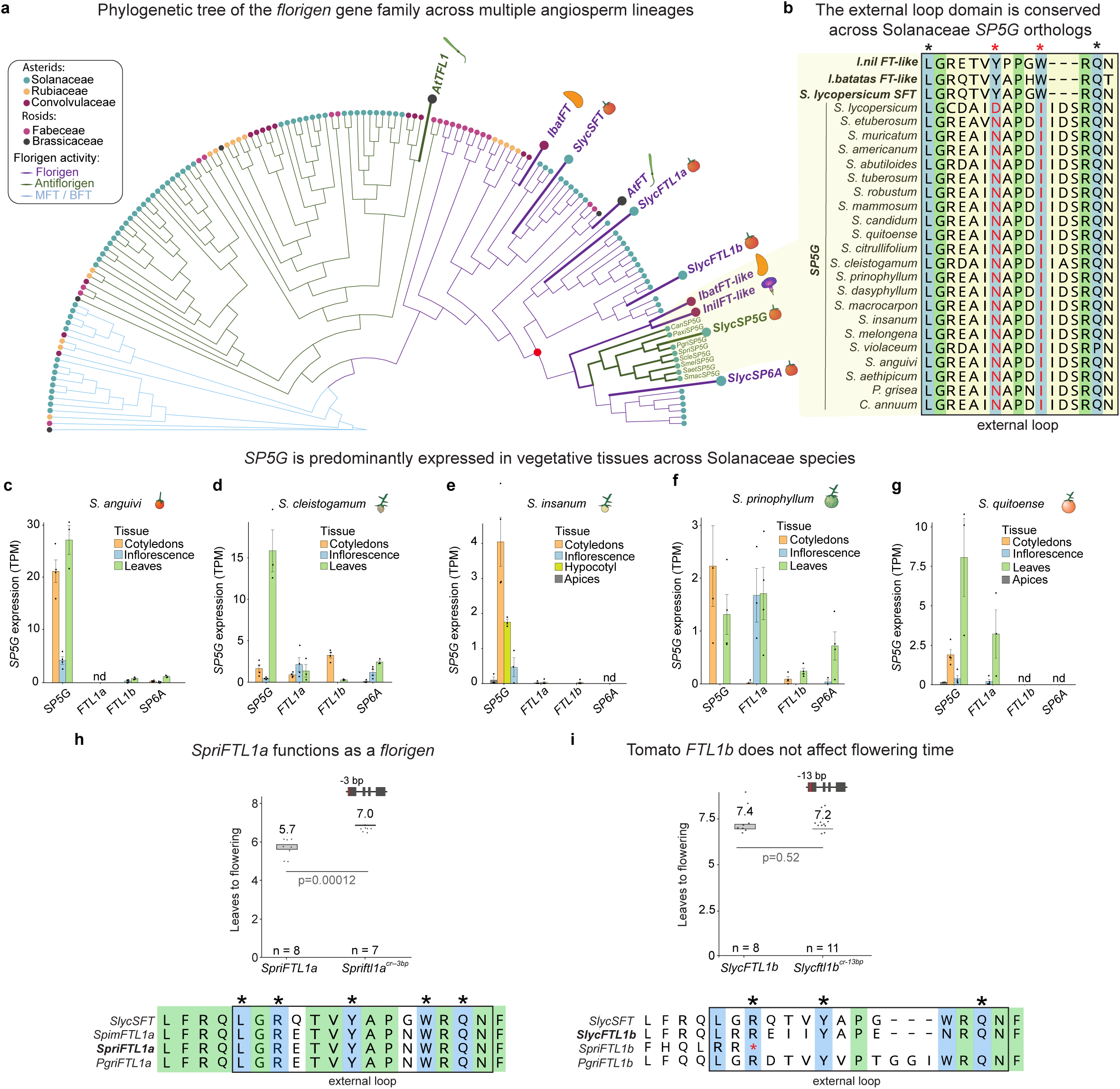
Phylogenetic analysis of the *florigen* gene family across angiosperms showing *SP5G* is a Solanaceae neofunctionalized antiflorigen. **(a)** Protein-based phylogenetic tree of the *florigen* gene family across angiosperm families showing three clades: FLOWERING LOCUS T (FT), TERMINAL FLOWER 1 (TFL1), and BROTHER OF FT (BFT) or MOTHER OF FT (MFT). **(b)** Alignment of the 14 amino acid external loop domain of Solanaceae SP5G orthologues. The tomato FT (SFT) and the closest sweet potato (*Ipomoea batatas*) homolog retain the canonical florigen motif in the external loop domain. Asterisks and blue color mark key florigenic amino acid residues with red asterisks marking critical changes in SP5G orthologues relative to florigens. Green color marks identical residues. **(c-g)** Expression of *SP5G* and its paralogues in different tissues of the indicated Solanaceae species. Bar charts present means ±s.e. TPM, Transcript Per Million. nd, not detected. Expression values were obtained from transcriptome data published in Benoit et al., (2025)^22^. Sample size (n) for each tested tissue is presented in **Supplementary Table 3**, representing the number of biological replicates collected from separate plants. **(h)** Flowering time of plants homozygous for functional *SpriFTL1a* compared to CRISPR-Cas9-engineered *SpriFTL1a^cr-3bp^* mutants. **(i)** Flowering time of plants homozygous for functional *SlycFTL1b* and CRISPR-Cas9-engineered *SlycFTL1b^cr-13bp^* mutants. Alignment of the external loop domain of *FTL1a* (h) and *FTL1b* (i) orthologues are shown below the plots. Means and a scheme of the *SpriFTL1a* (h) and *SlycFTL1b* (i) genes are shown above each plot. The red rectangles on the first exon denote the indicated deletion sizes (noted above). Sample size (n) in (h), (i) represents the number of plants quantified per genotype. A two-tailed, two-sample t-test was performed to compare means in (h), (i). Box plots in (h), (i) show the 25th, 50th (median) and 75th percentiles.

**Extended Data Figure 3.**
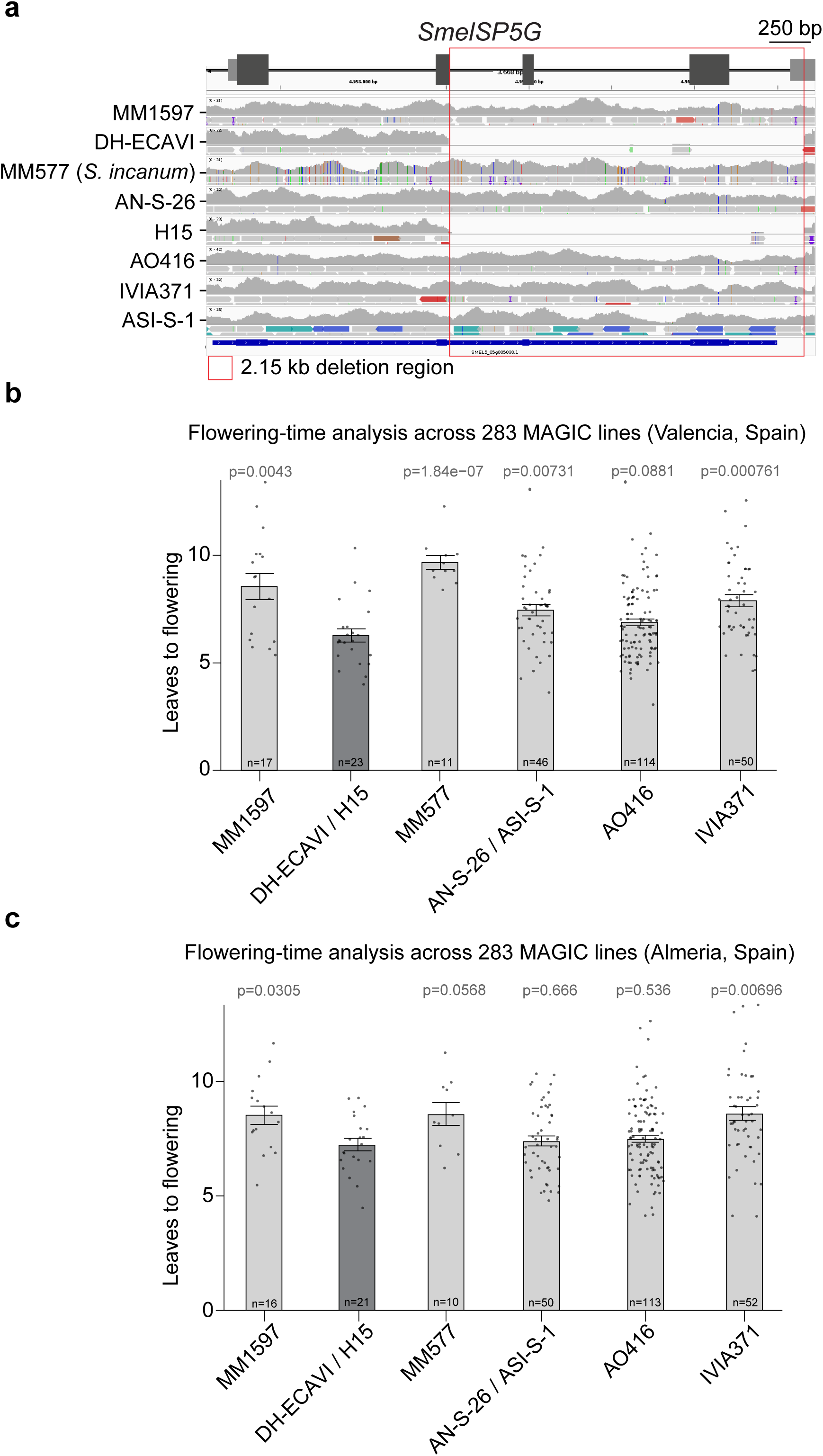
The *SmelSP5G* coding deletion allele drives early flowering in Brinjal eggplant. **(a)** Mapping of short-reads of the MAGIC genotypes to the wild relative *S. incanum* (accession MM577) identifies a 2.15-kb deletion in *SmelSP5G* in the varieties H15 and DH-ECAVI. **(b and c)** Flowering times of 283 MAGIC lines in two different environments in Spain [Valencia (b) and Almeria (c)]. Bar charts present means ±s.e. Sample size (n) in (b) and (c) represents the number of quantified plants. A two-tailed, two-sample t-test was performed to compare all means to the H15/DH-ECAVI genotypes, which are homozygous for the *Smelsp5g^del^* deletion allele.

**Extended Data Figure 4.**
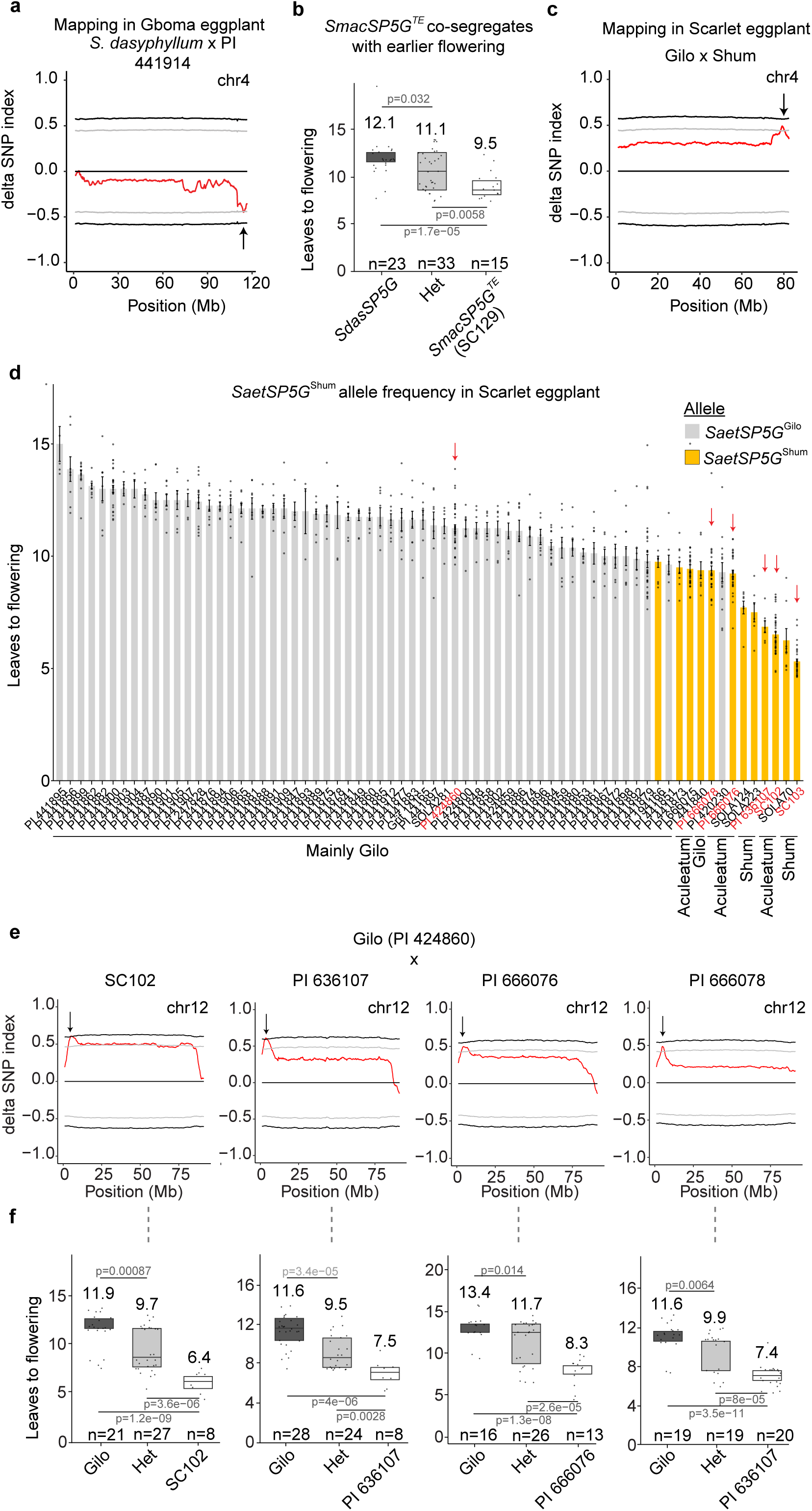
A shared flowering-time QTL on chromosome 4 in African eggplants. **(a)** QTL-seq identifies a locus on chromosome 4 affecting flowering time in Gboma eggplant. Black arrow marks the position of the peak exceeding the significance threshold. **(b)** *SmacSP5G^TE^* co-segregates with early flowering in an F2 population. **(c)** QTL-seq identifies a locus on chromosome 4 affecting flowering time in Scarlet eggplant. Black arrow marks the position of the peak exceeding the significance threshold. (**d**) Allele-frequency analysis of *SaetSP5G*^Shum^ across 70 Scarlet eggplant accessions. Accessions marked in red denote the parental Gilo and Shum parents used for QTL-seq. Bar charts present means ±s.e. Sample size (n) in (d) represents the number of plants quantified per species and is presented in **Supplementary Table 3**. **(e)** QTL-seq analyses identify a shared flowering-time QTL on chromosome 12 in Aculeatum ornamental accessions of Scarlet eggplant. Black arrow marks the position of the peak exceeding the significance threshold. Accession codes for each parental early flowering accessions are indicated. **(f)** A *SaetSP5G*^Shum^ molecular marker in the QTL-seq peak co-segregates with early flowering in four independent F2 populations. Means in (b), (f) are shown above box plots. Significance (p < 0.05) in (b), (f) were determined by one-way ANOVA with post-hoc Tukey–Kramer HSD test. Box plots in (b), (f) show the 25th, 50th (median) and 75th percentiles.

**Extended Data Figure 5.**
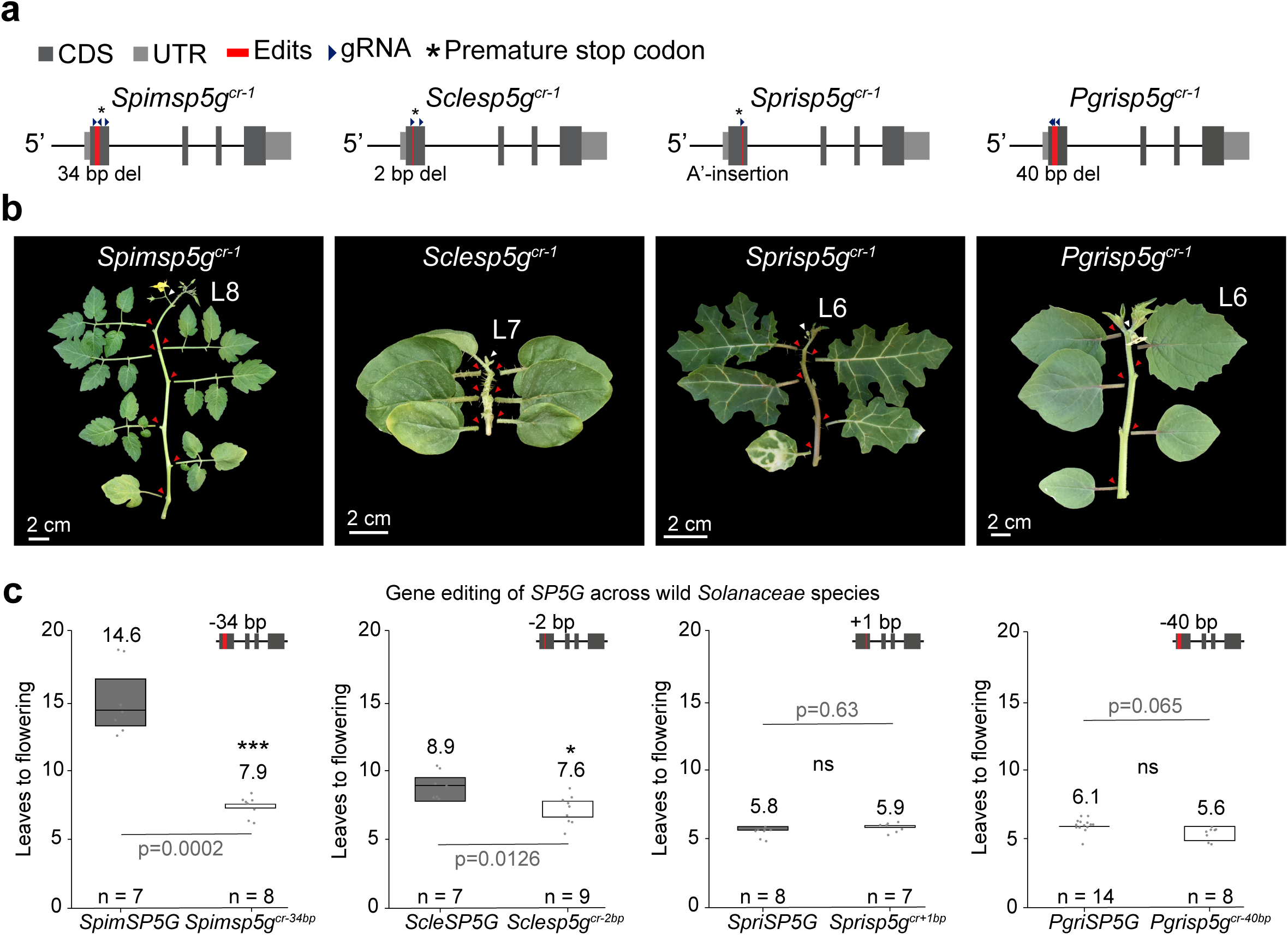
Pan-Solanaceae gene editing of *SP5G* reveals its functional diversification. **(a)** Schematics of *SP5G* gene and CRISPR-Cas9 loss-of-function mutations in the indicated species. **(b)** Representative plants and flowering times, measured in leaves (L) before the first inflorescence (white arrowheads), of gene-edited *SP5G* mutants in the indicated species. **(c)** Flowering time of WT and CRISPR-Cas9 engineered mutants of *SP5G* in the indicated species. Means are shown above box plots. Red rectangles in gene schematics denote mutations with sequence changes indicated above. A two-tailed, two-sample t-test was performed to compare means of WT and mutant plants in (c), asterisks indicate significant difference (\**p* < 0.05, \*\*\**p* < 0.01). ns, not significant. Sample size (n) in (c) represents the number of plants quantified per genotype. Box plots in (c) show the 25th, 50th (median) and 75th percentiles.

**Extended Data Figure 6.**
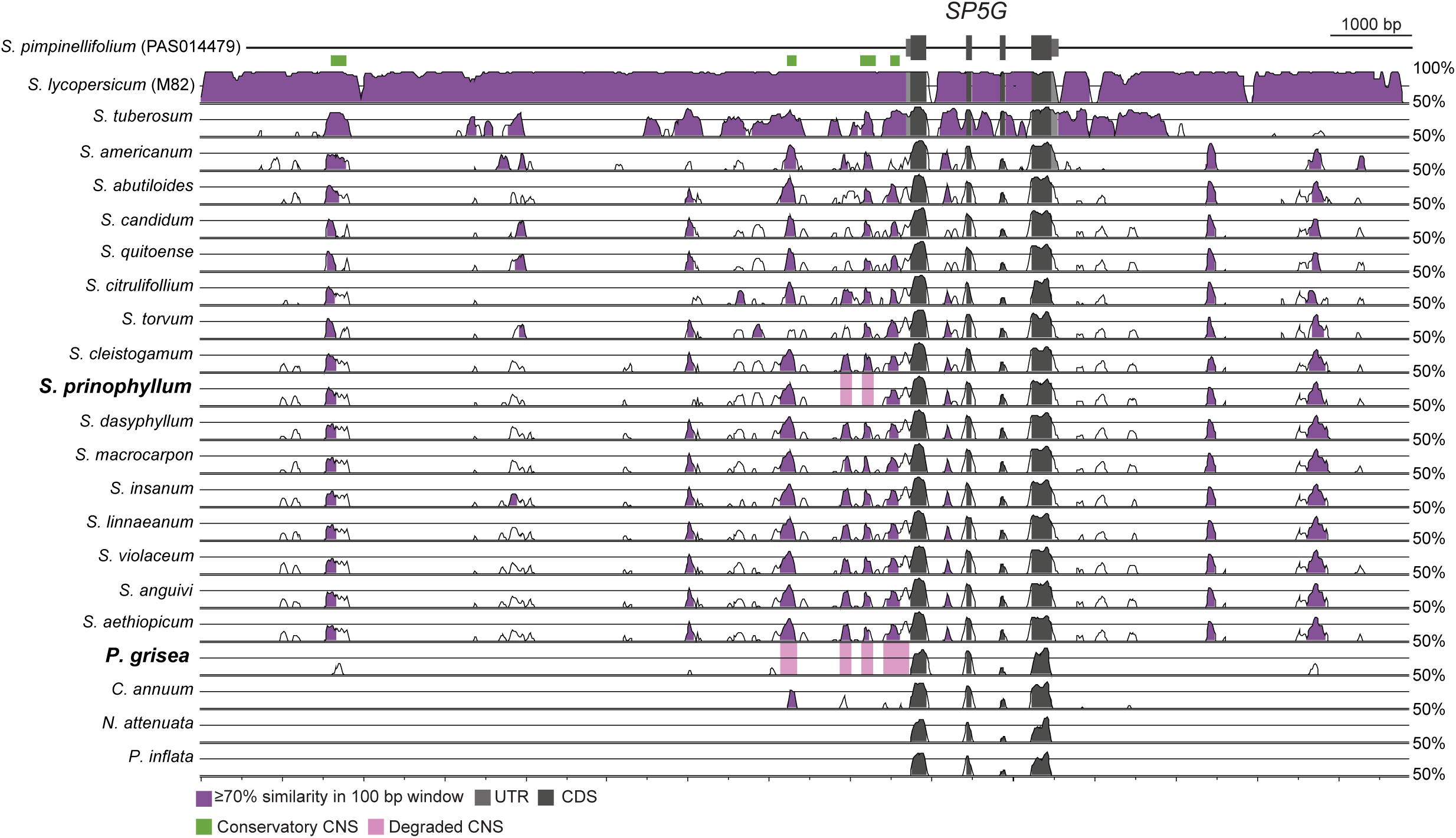
mVISTA plots showing *SP5G cis*-regulatory conservation and its overlap with conserved non-coding sequences (CNSs) defined by Conservatory. Shown are sequence alignments of 21 *SP5G* orthologues using *SpimSP5G* (accession PAS014479) and its surrounding upstream (∼8.7 kb) and downstream (∼4 kb) *cis*-regulatory regions as the reference. Conservation (purple and dark grey) was calculated as sequences with 70% similarity in 100-bp windows. Green rectangles below the *SpimSP5G* reference are positions of Conservatory CNSs^53^. Light gray regions in gene model denote UTRs and dark gray denote CDS in gene model and conservation. Pink vertical windows within the alignment map mark degraded CNSs.

**Extended Data Figure 7.**
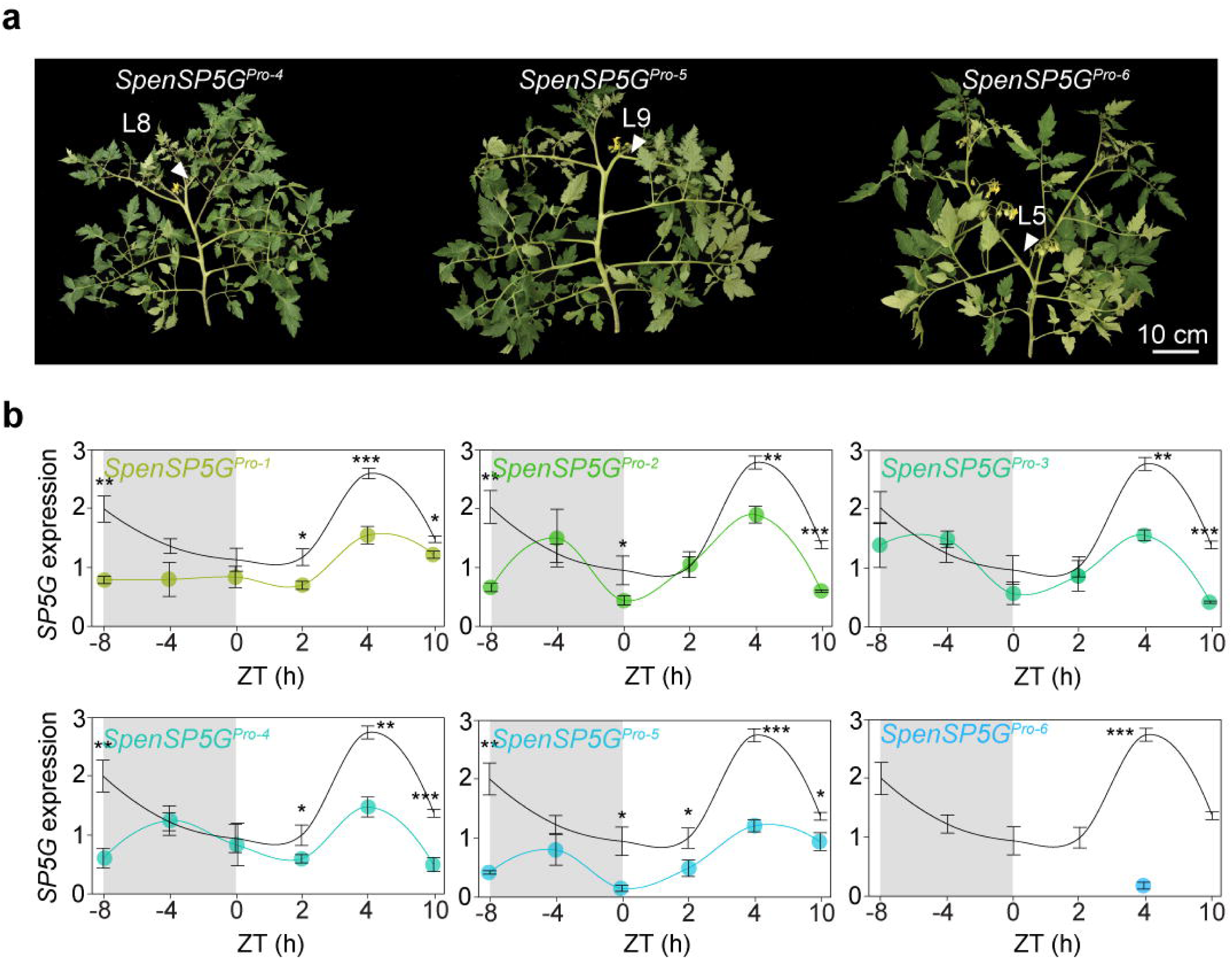
CRISPR-Cas9 engineered *SpenSP5G^Pro^* alleles reduce *SP5G* expression and flowering time. **(a)** Representative images of a subset of CRISPR-Cas9 *SpenSP5G^Pro^* alleles. White arrowheads indicate leaf number above the first inflorescence. **(b)** RT-qPCR diurnal expression analysis of the engineered *SpenSP5G^pro^* alleles in 6 time-points throughout 24 hours. Black line in each plot denotes the expression of *SpenSP5G*, the colored line denotes the engineered *SpenSP5G^Pro^*allele. *SpenSP5G^Pro-6^* was analyzed at a single time point (ZT+4). Grey areas mark dark periods. Data points are presented as means ±s.e. Sample size (n) is presented in **Supplementary Table 3**, representing the number of biological replicates (cotyledons) collected from separate plants at each time point. A two-tailed, two-sample t-test was performed to compare means, asterisks indicate significant difference (* for *P* < 0.05, ** for *P* < 0.01, *** for *P* < 0.001).

**Extended Data Figure 8.**
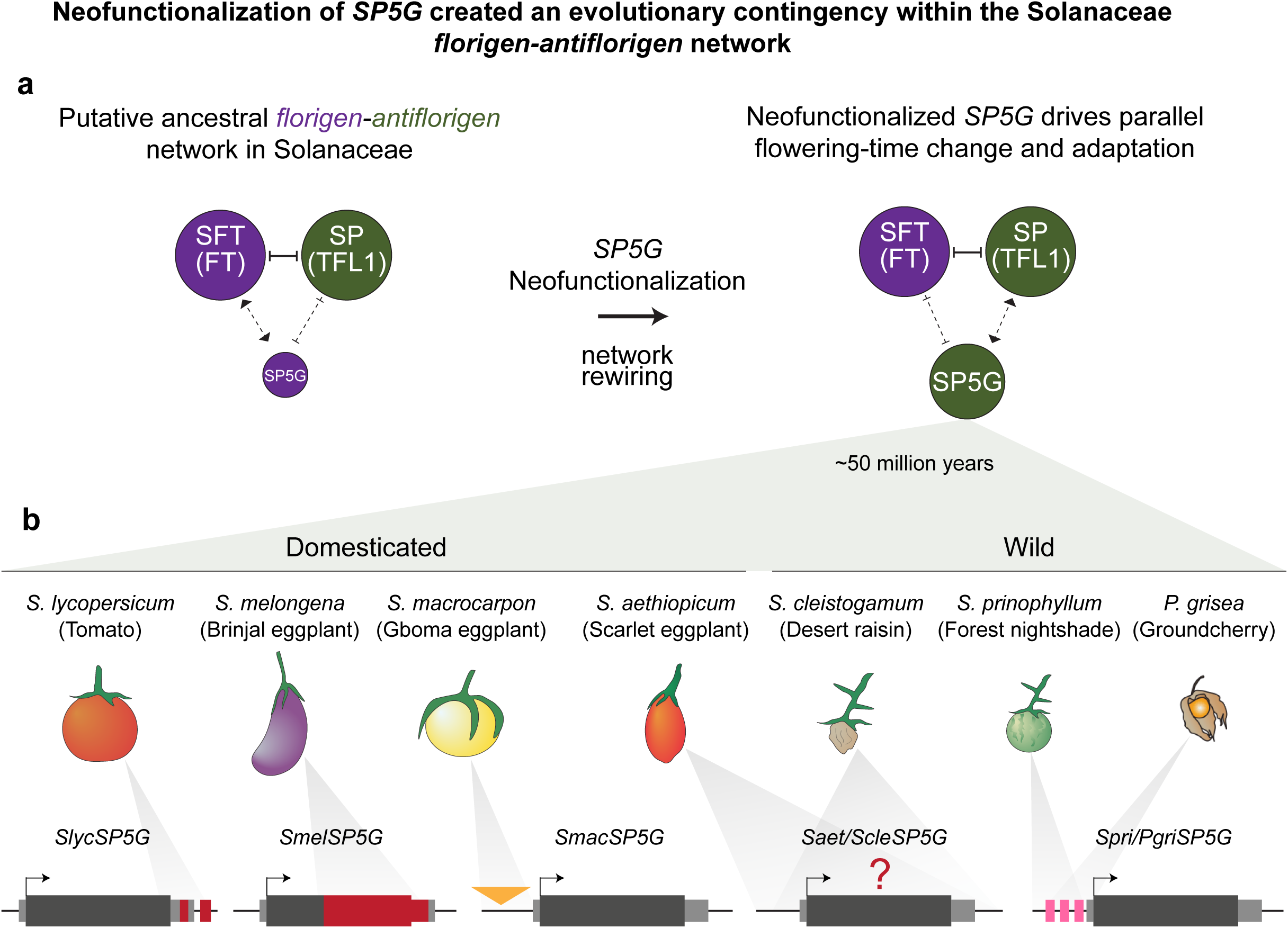
Neofunctionalization of *SP5G* created an evolutionary contingency within the Solanaceae *florigen-antiflorigen* network. **(a)** Simplified schematic of the ancestral *florigen-antiflorigen* network in Solanaceae and the influence of *SP5G* neofunctionalization on network interactions. Solid lines denote functional interaction, dashed lines denote putative positive (arrowheads) or negative (vertical line) interactions among paralogues. **(b)** Recurrent mutation of *SP5G* underlie parallel flowering-time change in domesticated and wild Solanaceae species. Schematics of *SP5G* gene bodies and types of alleles marked in different colors and associated with their corresponding species. Red denotes deletions in tomato and Brinjal eggplant, yellow denotes transposon insertion in Gboma eggplant, and pink denotes *cis*-regulatory degradation in *S. prinophyllum* and *P. grisea*. Question marks denote uncharacterized causative mutations.

