## Supplementary material for "A neofunctionalized flowering antagonist channeled Solanaceae evolution": SI Guide

**Supplementary Notes - Genome-wide Tests for Selective Sweeps**

**Methods**

Pages 2-7.

**Results**

Pages 8-12.

**Supplementary Tables**

Pages 13-30.

Supplementary Table 1. Accessions analyzed, by population and dataset.

Supplementary Table 2. Gene coordinates and focal windows.

Supplementary Table 3. Published sweep evidence at known domestication loci.

Supplementary Table 4. Simulation parameters.

Supplementary Table 5. Detection power, within-population statistics, within SLC.

Supplementary Table 6. Detection power, cross-population statistics, SLC against SP.

Supplementary Table 7. Detection power for both deletions.

Supplementary Table 8. Within-population statistics at target loci.

Supplementary Table 9. Cross-population statistics at target loci, SLC against SP.

Supplementary Table 10. Cross-population statistics at target loci, SLL against SP and SLC.

Supplementary Table 11. Within-population statistics at target loci, eggplant.

Supplementary Table 12. Cross-population statistics at target loci, eggplant.

Supplementary Table 13. Within-population statistics at known domestication loci.

Supplementary Table 14. Cross-population statistics at known domestication loci, SLC against

SP.

Supplementary Table 15. Cross-population statistics at known domestication loci, SLL against

SP and SLC.

**Supplementary Figures**

Pages 31-34.

Supplementary Figure 1. Detection power.

Supplementary Figure 2. Detection power against end frequency and sweep age.

Supplementary Figure 3. Key florigenic residues are largely conserved across Solanaceae FT-like paralogues.

Supplementary Figure 4. The frequency of the *SmelSP5G* coding deletion allele (*Smelsp5g^del^*)

is widespread in European and Middle Eastern accessions.

**Supplementary Discussion**

Pages 35-36

**References**

Pages 37-39.
