## Supplementary Information for "A neofunctionalized flowering antagonist channeled Solanaceae evolution"

### Supplementary Notes

#### Genome-wide Tests for Selective Sweeps

**Datasets.** We analyzed three datasets (**Supplementary Table 1**). The first, referred to below as the CSHL dataset, was assembled at Cold Spring Harbor Laboratory and consisted of over 100 accessions[32,34]. It is in coordinates of the wild *S. pimpinellifolium* (SP) assembly PAS014479_MAS2.0[32]. The second is the 667-sample dataset, a survey of flavor variation across tomato domestication and improvement[36]. In this case, we obtained a VCF file from the authors and analyzed it in the coordinate frame of BGV1.3, their reference genome. Coordinates and gene models differ between the assemblies, so we cross-referenced by Gene ID for all comparisons (**Supplementary Table 2**). The third dataset consists of 324 brinjal eggplant (*S. melongena*) accessions, in the coordinate frame of assembly GPE001970[45,46].

**Population assignment.** Population labels follow Supplementary Table 3 of Razifard et al., (2014)[34] which gives an assigned population for 294 accessions. Accessions were matched by name. Besides SP, the other two groups are the semidomesticated *S. lycopersicum* var. *cerasiforme* (SLC) and the domesticated *S. lycopersicum* var. *lycopersicum* (SLL). We analyzed all three, with SP–SLC as the primary comparison and SLL considered as well because the 52-bp deletion approaches fixation there. For the CSHL dataset we used Razifard et al., (2014)[34] assignments directly. For the Li et al., (2026)[36] dataset we used the assignments of Li et al., (2026)[36] except where the two sources were inconsistent in their SLC assignments. We excluded those accessions rather than reassigning them [G528/TS-75 SLC in Li et al., (2026)[36], SLL in Razifard et al., (2014)[34] and G285/LA2260 SLL in Li et al., (2026)[36] SLC PER, the Peruvian SLC subgroup, in Razifard et al., (2014)[34]]. We also excluded G35 (BGV006775), the accession used as the BGV1.3 reference.

For Brinjal eggplant, we considered three groupings of the 324 accessions: Indian accessions (IND, $n$ = 47) against Middle Eastern and European accessions (MEU, $n$ = 121), excluding Indian *S. insanum*; IND and MEU combined ($n$ = 168) against all remaining accessions (COMP, $n$ = 156); and all 324 pooled.

**Individual quality control.** We removed individuals with excess missing genotypes. Within each population we excluded any individual whose F_MISS exceeded the median plus three unscaled median absolute deviations. Thresholds were estimated separately for each population and dataset, excluding chromosome 5. This left 15 SP and 81 SLC accessions in the CSHL dataset, from 15 and 84, and 30 SP and 186 SLC in the Li et al. dataset, from 34 and 222 (**Supplementary Table 1**). The same filters applied to SLL left 10 of 11 accessions in the CSHL dataset and 236 of 299 in the Li et al., (2026)[36] dataset. Notably, for the Li et al., (2026)[36] dataset, we had to convert haploid-missing genotypes (.) to diploid-missing (./.) for proper downstream processing. For eggplant, we found that similar filtering rules led to a more complicated situation where thresholds varied across chromosomes, so we opted for an alternative approach where all 324 accessions were retained but missingness was controlled at the site level.

**Loci.** We analyzed the same florigen genes in every dataset: *SP5G*, *SP6A*, *FTL1a*, *FTL1b* and *SFT* (**Supplementary Table 2**). In tomato, however, *FTL1a* and *FTL1b* lie about 13 kb apart and fall in the same focal window, so were treated as one locus, leaving four. *SFT* retains the ancestral function and serves as a control. For tomato only, we also included eight loci reported in the literature as targets of domestication sweeps (**Supplementary Table 3**). *LNK2* and *FAS*/*SlCLV3* were omitted from the Li et al., (2026)[36] analysis because they lack validated gene models in BGV1.3.

### Methods

All statistics were calculated in 100-kb windows with a 10-kb step. For each gene the focal window is the one whose center lies closest to the gene center, with ties resolved to the lower start position.

**Site filters.** We calculated $\pi$, $F_{ST}$ and Tajima's $D$ from biallelic SNPs with QUAL $\geq$ 30 and at least 70% of alleles called across the combined panel (AN70). We repeated the analysis at AN50, AN60 and AN80. Variants were reduced to primitive SNP coordinates. Positions that could not be mapped uniquely were removed. Filters were applied once to the combined panel, not within populations, so both populations are scored at the same sites. The AN70 requirement does not carry into the haplotype analyses.

**Nucleotide diversity.** Per-site diversity is

$$\pi_{\text{site}} = \frac{2\,\text{AC}\,(\text{AN}-\text{AC})}{\text{AN}\,(\text{AN}-1)}, \tag{S1}$$

We summed these values over each window and divided by the nominal window length. Uncallable bases cannot be identified from the VCFs, so they are not masked and absolute $\pi$ may be slightly depressed. We report $\pi$ within each population and their ratio, with the population closer to the ancestral state in the numerator: $\pi_{\text{SP}}/\pi_{\text{SLC}}$, $\pi_{\text{SP}}/\pi_{\text{SLL}}$ and $\pi_{\text{SLC}}/\pi_{\text{SLL}}$ in tomato, and $\pi_{\text{IND}}/\pi_{\text{MEU}}$ in eggplant. The denominator cancels in the ratio, since both populations are scored over the same window.

**Population differentiation.** We report two $F_{\text{ST}}$ estimators. Weir–Cockerham[97] is the estimator used by Lin et al., (2014)[35] and Li et al., (2026)[36]. Hudson's ratio-of-sums estimator[98] is better behaved at unequal sample sizes[99], which matters at $n$ = 15 against 81. Each is ranked against its own genome-wide background.

**Tajima's *D*.** We calculated Tajima's $D$ separately within each population and dataset. The classical estimator assumes a fixed number of sampled chromosomes. We therefore required complete calls within the population: a site contributed only where $\text{AN}_{\text{pop}} = 2N_{\text{pop}}$. Two compared populations therefore retain different sites within the same window. Within a window, $S$ is the number of complete-case segregating sites. The pairwise-difference term is $\sum_i 2\,\text{AC}_i(n-\text{AC}_i)/[n(n-1)]$ over those sites, with $n$ the fixed chromosome count. Watterson's estimator is $\theta_W = S/a_1$ with $a_1 = \sum_{i=1}^{n-1} 1/i$. We combined these under the Tajima normalization[100], with variance constants determined by $n$. Windows with no segregating sites or with non-positive variance are unrankable. The pairwise-difference term is a window sum over segregating sites, not the per-base $\pi$ above.

**Haplotype statistics.** We used $nS_{\text{L}}$[101] and XP-$nS_{\text{L}}$[102] rather than iHS or XP-EHH, since no recombination map is available for these assemblies. Both have been shown to have good power for soft sweeps. Both were computed with selscan v2.1.2[103] in --unphased mode, which encodes each genotype as a multilocus genotype (MLG) state (0/0 → 0, heterozygous → 1, 1/1 → 2). From v2.1.3 onward, segregating sites can be dropped if individuals differ only in the homozygous state, which was not the desired behavior. We therefore used v2.1.2.

**Sites considered for $nS_{\text{L}}$.** The set of sites considered was built once on the combined panel of a pair of compared populations, before splitting. We required biallelic SNPs, QUAL ≥ 30, complete genotypes across the combined panel, polymorphism in the combined panel, and a combined-panel minor allele frequency ≥ 0.05. All records at any duplicated CHROM+POS coordinate were dropped. The same coordinates were supplied to both populations. We ran selscan --nsl --unphased --maf 0.05 without --keep-low-freq. The frequency threshold is therefore applied again within each population, and the sites scored differ between them. Scores were normalized genome-wide across all twelve chromosomes, separately within each population and dataset, in 100 frequency bins. The window statistic is the fraction of finite normalized scores with $|nS_L| > 2$.

**Sites considered for XP-$nS_L$.** The set of sites considered was built jointly on the query and reference panels. We restricted to biallelic SNPs. We deduplicated records identical in identity and genotype and removed positions that could not be mapped uniquely. We required complete diploid genotypes at every site in every individual of the pair, applied MLG encoding, and kept sites with at least two distinct MLG states across the pair. Query and reference VCFs were split from this set and checked to carry the same sites in the same order. Polymorphism was not required within each population separately, so sites fixed for different alleles between populations remain eligible. The command was selscan v2.1.2 --xpnsl --unphased --maf 0 --keep-low-freq --max-extend-nsl 100, with the default --max-gap of 200,000 bp and --gap-scale of 20,000 bp. Scores were normalized genome-wide across all twelve chromosomes, separately within each comparison and dataset, using norm --xpnsl --files. In tomato the derived population is the query: SLC against SP, and SLL against either SP or SLC. In eggplant MEU is the query and IND the reference. Positive scores therefore indicate longer haplotypes in the query population. We report the fraction of scores above +2 and the fraction below −2.

**Percentile backgrounds.** Each focal value is ranked against the genome-wide distribution of rankable windows for the same statistic, panel and population, using ascending average ranks with midranks for ties. For $\pi$, the $\pi$ ratio and $F_{ST}$ the background is the set of windows on all twelve chromosomes at which all three are finite, so they share one denominator: $N = 78{,}640$ in the CSHL panel, 78,639 in its SLL comparisons, and 78,737 in the Li et al., (2026)[36] panel. Tajima's $D$ has its own denominators because of its complete-case requirement: 78,624 for SP, 78,595 for SLC and 78,622 for SLL in the CSHL panel; 78,675, 78,408 and 78,318 in the Li et al., (2026)[36] panel. A low Tajima's $D$ percentile therefore denotes an unusually negative value. In eggplant they are 112,927, and for Tajima's $D$ 112,712 for IND and 112,582 for MEU. For $nS_L$ and XP-$nS_L$ the background is windows with at least 20 scored sites. In tomato it excludes chromosome 5, which carries *SP5G*, so *SP5G* and *SP6A* are ranked against a background that does not contain them; in eggplant all twelve chromosomes are used.

**Validation.** Window aggregation was reimplemented independently, with a maximum $|\Delta F_{ST}|$ of $8.2 \times 10^{-12}$. Hudson $F_{ST}$ components were revalidated on 5,000 sites with no mismatches. Weir–Cockerham $F_{ST}$ agreed with VCFtools[104] to within $5.0 \times 10^{-7}$, against a pre-specified tolerance of $5 \times 10^{-6}$, and our Tajima's $D$ agreed with vcftools --TajimaD to within $3.2 \times 10^{-7}$ on a synthetic dataset with complete diploid genotypes. Varying the allele-number threshold (AN50/60/70/80) and duplicate handling moved percentiles by less than 0.4 points. Re-running Li et al., (2026)[36]'s own command on their VCF reproduced the $F_{ST}$ landscape of their published table (Spearman $\rho = 0.994$ across 770 windows, means 0.249 against 0.259). On one chromosome, selscan v2.1.2 and v3.0.1 returned identical scores at every shared locus, differing only in which positions survive. Notably, Lin et al., (2014)[35] analyzed a different panel aligned against a different assembly and reported genome-wide diversity of $3.23 \times 10^{-3}$ in SP (their PIM) and $1.74 \times 10^{-3}$ in SLC (their CER), a ratio of 1.856. Our medians are comparable at 0.003761 and 0.002228, a ratio of 1.688.

**Simulation design.** All simulations are of tomato, where we have two datasets and a published demographic history to draw on. No comparable demographic model is available for eggplant, so the power estimates below are not transferred to it.

**Demographic model.** All simulations used demographic histories inferred by Razifard et al., (2014)[34] with dadi[105], Supplementary Table 7, under two scenarios. The origin of SLC (model 5) has a population split followed by isolation and then secondary contact with a size change, fitted to SP against SLC and SLL pooled. Redomestication (model 3) is the later step, fitted to SLC MEX against SLL, with secondary contact but no size change. We simulated the 83-bp deletion in the context of the first scenario and the 52-bp deletion in the context of the second (discussed further below).

Scaled parameters were converted using $T_{gen} = 2N_{ref}T_{scaled}$, $N_e = vN_{ref}$ and $m_{gen} = m_{dadi}/(2N_{ref})$. We obtained $N_{ref} = T_{div}/[2(T_1 + T_2)]$, giving 42,560 for model 5 and 5,694 for model 3, assuming one generation per year. We preserved dadi's forward-time migration orientation. The fitted $\theta$ was not used, since mutation rate and sequence length were set directly for each experiment.

Two effective sizes were recalibrated. In model 5, the fitted descendant size describes SLC and SLL pooled and gives simulated diversity about 2.2× the value reported for SLC, so we replaced it with the size that brings simulated diversity close to that value, leaving SP about 10% below its own reported diversity (**Supplementary Table 4**). The second recalibration, for the single-population sweep approximation, is described below.

**Simulators.** $\pi$, Tajima's $D$ and $nS_L$ were simulated in msprime 1.3.3[106] using SweepGenicSelection, in the selected population alone, since that model cannot run with a second active population. Because this single-population approximation changes neutral diversity, the present-day size in it was set to 31,500 to restore diversity to the two-population value. The two-population statistics, $\pi$ ratio and XP-$nS_L$, were simulated in discoal[107], which permits sampling several subpopulations while restricting the sweep to one.

**Sweep simulations.** Samples were simulated as one haplotype per accession, matching the unphased empirical analyses. The $\pi$ ratio is the exception and was computed on $2n$ allele counts, as in the empirical scan it is compared against. In the main set of simulations, sweeps were placed at the center of the simulated region and conditioned on a final frequency of 0.99, with three selection coefficients and eight starting frequencies ranging from a single copy, $1/(2N_e)$, up to 0.37, the estimated starting frequency of the 83-bp deletion in SP. For the two-population statistics we also varied sweep age, using the present day as the most favorable case and $T_2 = 45{,}956$ generations as the age a sweep at the origin of SLC would have under this model. The 52-bp deletion was treated as a hard sweep in SLL against SLC and scored at 0, 500 and 711 generations, the last of these being the estimated age of the re-domestication event.

The window scales in **Supplementary Table 4** apply to $\pi$ and Tajima's $D$. $nS_L$ and XP-$nS_L$ were instead computed on fixed simulated regions, also given there. $nS_L$ was standardized within derived-allele-count bins against the unswept flanks of the same replicate, which is why the simulated region is wider than the scored window. XP-$nS_L$ was scored against matched neutral replicates rather than against flanks, since flank-based normalization fails for the cross-population statistic.

A second set of simulations resolved how detection power is affected by the final frequency of the allele and the time since selection ceased. Under the same demography and scenario, this narrower grid varied two parameters the main grid holds fixed. Sweeps were conditioned on final frequencies of 0.05, 0.20, 0.50, 0.70 and 0.99 rather than 0.99 alone, and sweep age was resolved at 0, 711,

2,500, 10,000, 25,000 and 45,956 generations rather than at its two endpoints. Cells whose final frequency was at or below the starting frequency describe no trajectory and were not simulated. This grid was run for XP-$nS_L$ only, in the CSHL panel, at $s = 0.01$ and 0.05 and the 95% selfing rate, so it pertains to haplotypic signal only.

**Recombination and selfing.** Neither simulator implements selfing, so selfing enters only through effective recombination, $r_{eff} = r(1 - F)$ with $F = \sigma/(2 - \sigma)$. Because selfing is common in the sampled tomato lineages, the primary results use a rate of $9.5 \times 10^{-10}$ per bp per generation, the effective rate under 95% selfing. The outcrossed rate of $1 \times 10^{-8}$ and the rate under 99% selfing, $2.0 \times 10^{-10}$, are reported as sensitivity analyses (**Supplementary Table 4**). No genetic map was used, since none is available for this assembly. Effective sizes were not rescaled for selfing, because they were calibrated against observed diversity and therefore already absorb its effect.

**Detection threshold and power.** For every parameter combination we simulated a matched neutral set under the same demography, window size, sample size and recombination rate, differing only in the absence of the sweep. Power was the fraction of sweep replicates falling beyond the 5% tail of that matched neutral set. We used the lower tail for $\pi$ and Tajima's $D$, and the upper tail for the $\pi$ ratio, $nS_L$ and XP-$nS_L$. This is a fixed 5% false-positive rate per window, not a false discovery rate. The 5% tail matches the window-level outlier criterion used in the empirical scan these simulations are intended to inform. The two-population statistics were run both with and without migration between populations. Reported power uses migration on for the 83-bp scenario, since model 5 includes secondary contact, and migration off for the 52-bp scenario, since discoal cannot implement a timed migration change.

**Limitations.** Recombination is uniform with no genetic map, selfing is modeled only through effective recombination rather than an explicit mating model, and no callable-site mask was applied. discoal seeds the standing variation in the sweeping population only, so the reference population does not carry the variant before the sweep. For XP-$nS_L$ this makes the simulated standing-variation comparison optimistic, since a private selected allele gives a cleaner haplotype comparison than a shared standing variant. Except for the $\pi$-ratio zero-diversity floor, replicates in which a statistic could not be computed were dropped rather than imputed.

### Results

#### Power detection

To measure our detection power for sweeps, we simulated datasets under the demographic models inferred by Razifard et al., (2014)[34], assuming a grid spanning eight starting frequencies from a single copy to 0.37, three selection coefficients, three recombination rates, two sample panels and two sweep ages for the 83-bp scenario, three for the 52 bp. Unless stated otherwise, values below are for 100-kb windows, 95% selfing and migration between populations.

Not surprisingly, we found that the best case for detection was a recent hard sweep. For example, a hard sweep that has just fixed is recovered at rates of 0.99 to 1.00 by $\pi$, 0.98 to 1.00 by Tajima's $D$, 0.91 to 0.97 by the $\pi$ ratio and 0.92 to 1.00 by XP-$nS_L$ (**Supplementary Fig. 1a**, **Supplementary Tables 5 and 6**). The weakest discriminator in this case was $nS_L$, at 0.47 to 0.59. Among the variables we considered, the starting frequency $f_0$ mattered most, with power declining rapidly with $f_0$. For example, at $s = 0.05$, power for $\pi$ drops from 0.999 for a hard sweep to 0.238 by $f_0 = 0.005$, and for Tajima's $D$ from 0.991 to 0.002. The $\pi$ ratio and XP-$nS_L$ maintained reasonable power somewhat longer, with rates of 0.832 and 0.728 at $f_0 = 0.02$, falling to 0.494 and 0.100 by $f_0 = 0.1$. By $f_0 = 0.37$, the frequency observed for the 83-bp deletion in SP, no scenario with migration between populations gave power above 0.116, and none without it gave power above 0.278. Sweep age behaved similarly, with reasonable power at young sweeps ($\tau = 0$) substantially declining for older sweeps ($\tau = T_2 = 45{,}956$ generations, the age expected for a sweep at the origin of SLC) as recombination breaks up haplotypes and mutation and migration restore diversity (**Supplementary Tables 6**).

Selfing matters only for $nS_L$ (**Supplementary Fig. 1b**), which recovers a hard sweep at 0.742 outcrossed but 0.489 under 95% selfing and 0.414 under 99%. Power for $\pi$ and XP-$nS_L$ is essentially unchanged. At $f_0 = 0.37$ power spans 0.002 to 0.084 outcrossed, 0.000 to 0.116 under 95% selfing and 0.000 to 0.156 under 99%, so the recombination assumption does not rescue a high-frequency start.

Larger panels help in the middle of the grid, not at its ends (**Supplementary Fig. 1c**). Raising the SLC panel from 81 to 186 lifts XP-$nS_L$ from 0.399 to 0.686 at $f_0 = 0.05$, while the $\pi$ ratio gains nothing, 0.706 against 0.694, because it compares diversity rather than haplotype length. By $f_0 = 0.37$

the advantage is gone for every statistic, XP-$nS_L$ at 0.002 against 0.004. The two deletions therefore fall in very different places on this surface (**Supplementary Table 7**). The 83-bp deletion is standing variation at $f_0 = 0.37$ in SP, and under selection at the origin of SLC it also takes the oldest sweep age we tested, putting it in the worst region of both axes at once. The 52-bp deletion is a hard sweep in SLL against SLC at the much younger re-domestication event, where power is substantially higher (**Supplementary Fig. 1d**). At $s = 0.05$ the CSHL panel recovers it at 0.955 by $\pi$, 0.946 by the $\pi$ ratio and 0.998 by XP-$nS_L$ immediately after fixation, falling to 0.858 and 0.474 for the two cross-population statistics by the re-domestication age of 711 generations. The larger Li et al., (2026)[36] panel raises Tajima's $D$ from 0.294 to 0.988 and XP-$nS_L$ at that age from 0.474 to 0.846. A negative result is much more informative for the 52-bp deletion than for the 83 bp.

Resolving the end-frequency and sweep-age axes more finely shows where the signal is lost (**Supplementary Fig. 2**). Power requires the allele to rise to fairly high frequency. For example, from a single copy, XP-$nS_L$ power is 0.060 at a final frequency of 0.05 and 0.150 at 0.20, reaching 0.920 only by 0.50 (**Supplementary Fig. 2a**). Power then decays sharply with time since selection ceased. A completed hard sweep retains power of 0.928 after 2,500 generations but falls to 0.138 by 10,000 generations and 0.060 at $T_2$ (**Supplementary Fig. 2b**). A sweep that stops before fixation behaves the same way once selection ceases: an allele that rose only to 0.50 is recovered with power 0.950 immediately after selection ceased, but only 0.042 after 10,000 generations (**Supplementary Fig. 2c**). Across this finer grid — every final frequency, every age, both selection coefficients and both migration settings — no cell starting from $f_0 = 0.37$ exceeded 0.080, so neither a lower final frequency nor a younger sweep rescues the 83-bp case.

**Simulations of the 83-bp deletion at *SP5G***

Importantly, the 83-bp deletion is the least favorable case for the entire simulation grid. This deletion likely segregated at about 37% in SP before domestication, rising to about 98% in landraces and reaching fixation in modern SLL. Sweeps starting from $f_0 = 0.37$ give distributions of $\pi$, Tajima's $D$, $nS_L$ and XP-$nS_L$ that overlap the neutral distributions almost entirely, and power runs from 0.000 to 0.116 across both panels and all three selection coefficients. Model 5 includes secondary contact, so these are the values with migration on. Turning migration off raises the ceiling only to 0.278. Thus, if a sweep has occurred at this locus, our simulations indicate we will have poor power to detect it, at least with these datasets and tests.

**Target loci**

When we applied these tests to our four target loci, we found that most statistics were unremarkable across most genes (**Supplementary Tables 8 and 9**). Excluding tests in SP alone, which are of less relevance and where we had fewer samples, only five of 64 scored statistics exceeded the 95th percentile genome-wide. For example, XP-$nS_L$ for SLC relative to SP is high at *SP6A* (P98.2) and *FTL1a/b* (P97.1) in the Li et al., (2026)[36] dataset. Notably, the causal variant at *FTL1a/b* arose in cherry tomato rather than in SP; Song et al., (2020)[41] report it as homozygous in more than 99% of cultivars and 85% of cherry tomato accessions, while the wild species carry the ancestral allele. A rise from absence to 0.85 falls well inside the range our simulations recover. The apparent causal allele at *SP6A* is likewise absent from SP, so both are consistent with hard sweeps during domestication.

By contrast, *SFT* is high by XP-$nS_L$ in the direction opposite to that expected for a domestication sweep, for SP relative to SLC [P99.7 in the Li et al., (2026)[36] dataset]. This gene also shows elevated $F_{ST}$ [P97.5 in the CSHL dataset and P95.7 in the Li et al., (2026)[36] dataset]. *SFT* lies in a pericentromeric region, however, where markedly reduced recombination rates may contribute to a spurious signal of a sweep. Notably, *SP5G* is not an outlier by any statistic in either dataset. Its highest value is Weir–Cockerham $F_{ST}$, at P90.4 in the CSHL dataset and P84.1 in the Li et al., (2026)[36] dataset.

We also compared SLL with SP and with SLC, the case in which the 52-bp deletion segregates and the one our simulations identify as best powered (**Supplementary Tables 8 and 10**). The pattern matches the SP–SLC result. Of 169 rankable values, those reaching the 95th percentile fall almost entirely at *SP6A*, *FTL1a/b* and *SFT*: XP-$nS_L$ for SLL relative to SP is high at *FTL1a/b* (P99.5) and *SP6A* (P99.4) in the Li et al., (2026)[36] dataset, as is the $\pi$ ratio at *FTL1a/b* (P99.8 against SP, P99.9 against SLC), while *SFT* is again elevated by $F_{ST}$ [P99.5 in the CSHL dataset, P97.5 in the Li et al., (2026)[36] dataset]. *SP5G* is an outlier by one statistic in one panel: Tajima's $D$ within SLL in the CSHL dataset is −0.75 (P4.6), in the direction expected under a recent sweep. That panel contains ten accessions, however, and the same test in the much larger Li et al., (2026)[36]. SLL panel gives −0.45 (P25.8). No other *SP5G* value in any SLL comparison exceeds P86.

**Known domestication loci**

Similarly, the eight known domestication loci give few extreme values and little agreement between tests (**Supplementary Tables 13 and 14**). Eight of 154 values exceed the 95th percentile, spread over six of the eight loci, with *OVATE* and *LIN5* accounting for two each. No locus is flagged by the same test in both datasets.

*LIN5* carries a missense variant at a frequency of 0.022 in SP which rises to 0.787 in SLC[108]. Accordingly, it is elevated by XP-$nS_L$ for SLC relative to SP [P98.6, Li et al., (2026)[36]] and by $nS_L$ within SLC [P95.5, Li et al., (2026)[36]. Notably, a sweep from that starting frequency to that final frequency does fall in the detectable part of our XP-nSL power surface (**Supplementary Fig. 2a**). The deletions observed at *EID1* and *LNK2* are absent from all 38 SP accessions previously genotyped[109], so both are consistent with hard or low-initial-frequency soft sweeps, but these sweeps appear to have completed at strikingly different times. *EID1* is already at a frequency of 0.85 in SLC and near fixation in the Peruvian *cerasiforme* accessions, whereas *LNK2* remains at 0.40 in SLC and reaches fixation only in the populations that arose after tomato reached Europe[109]. We detect *LNK2* within SLC ($nS_L$ P99.3, CSHL) but not *EID1*, as predicted from **Supplementary Fig. 2b**.

*OVATE* also scores highly, by $nS_L$ within SLC (P99.0) and by XP-$nS_L$ for SLC relative to SP (P97.0), both in the Li et al., (2026)[36] dataset, as does *lc*/*SlWUS* by XP-$nS_L$ (P98.2, CSHL). The only published frequencies for these two loci come from a collection assembled by fruit shape and are reported as carrier counts rather than allele frequencies[110], and of the seven wild accessions in that collection only two are SP. The two genes also lie 1.2-Mb apart, a distance over which linkage disequilibrium in SLC is barely reduced[35], and the *OVATE* window contains several annotated genes, so the two signals may not be independent. *fw2.2*/*CNR* is detected only in the SLL comparisons below. *FAS*/*SlCLV3* is the one locus that no test identifies.

Extending these tests to the SLL comparisons produces similar results (**Supplementary Table 15**). Of 300 rankable values, 18 reach the 95th percentile, spread over seven of the eight loci and showing no more consistency than before: *OVATE* is high by XP-$nS_L$ for SLL relative to SP [P99.96 in the Li et al., (2026)[36] dataset] and by $nS_L$ (P99.5), and *LNK2*, *LIN5*, *GAME12*, *lc*/*SlWUS*, *EID1* and *fw2.2*/*CNR* each reach it by one or two tests. Here, only 6% of SLL values and 5% of SP–SLC values exceed the 95th percentile, against the 5% expected by construction. At the target loci the corresponding rates are 8% and 9%.

Together with our simulations, these observations suggest that, given our sample sizes, sample composition, and the demographic model for these groups, we are near the limits of detection for most domestication sweeps. Whether each sweep is detected depends heavily on its particularly evolutionary history, including selection strength, age and starting frequency, as well as its genomic position. Detection likely also depends on technical features such as sequencing depth, SNP density, and window placement relative to causal alleles.

**Brinjal eggplant**

We repeated the analysis in brinjal eggplant (*S. melongena*), using a single dataset of 324 accessions and the same five target loci. Here we treat *FTL1a* and *FTL1b* separately, because in this assembly they do not fall within a common window.

For eggplant, we considered three groupings of the same accessions. Our primary analysis is scenario C, which compares Indian accessions (IND, $n$ = 47) with Middle Eastern and European accessions (MEU, $n$ = 121), excluding Indian *S. insanum*; this grouping most closely matches the reported origin and distribution of the causal coding deletion. Scenario A instead compares IND and MEU combined ($n$ = 168) with all remaining *S. melongena* accessions (COMP, $n$ = 156). Scenario B pools all 324 accessions and therefore supports only within-population statistics.

Under scenario C, no statistic at any of the five loci exceeded the 95th percentile genome-wide (**Supplementary Tables 11 and 12**). The largest values were Tajima's $D$ in MEU at *FTL1b* (P91.0) and *FTL1a* (P90.7), and $nS_L$ in IND at *SP5G* (P89.6). As in tomato, *SP5G* was not an outlier by any statistic, and its differentiation between IND and MEU was close to the genome-wide median ($F_{ST}$ P20.5).

The other two groupings give a broadly similar picture. Scenario B yielded no value above the 95th percentile, its largest again being Tajima's $D$ at *FTL1a* (P90.7). Scenario A yielded seven of 55 scored statistics above that threshold, but they were concentrated at *SP6A* — XP-$nS_L$ for IND+MEU relative to COMP (P99.5), Weir–Cockerham $F_{ST}$ (P99.1) and Hudson $F_{ST}$ (P98.5) — together with Tajima's $D$ in COMP at *SP6A* (P96.6), *FTL1b* (P95.5) and *FTL1a* (P95.4), and $\pi$ in COMP at *SP6A* (P95.1). These shifts largely track the genome-wide background rather than the loci themselves: the median $F_{ST}$ between groups is three times higher under scenario C than under scenario A (0.072 against 0.023), so the same absolute value falls at a very different percentile. *SP5G* reached the 95th percentile under none of the three grouping.

### Supplementary Tables

**Supplementary Table 1.** Accessions analyzed, by population and dataset.

| | CSHL | | Li et al. | |
|---|---|---|---|---|
| | original | analyzed | original | analyzed |
| SP | 15 | 15 | 34 | 30 |
| SLC | 84 | 81 | 222 | 186 |
| SLL | 11 | 10 | 299 | 236 |
| Total | 110 | 106 | 555 | 452 |

**Supplementary Table 2.** Gene coordinates and focal windows.

| | Locus | Gene ID | Chr. | Gene span | Focal window[a] |
|---|---|---|---|---|---|
| CSHL PAS014479_MAS2.0 target loci | *SP5G* | Solyc05g053850 | 05 | 66,577,343–66,579,537 | 66,530,001–66,630,000 |
| | *SP6A* | Solyc05g055660 | 05 | 67,909,212–67,911,152 | 67,860,001–67,960,000 |
| | *SFT* | Solyc03g063100 | 03 | 29,894,026–29,897,862 | 29,850,001–29,950,000 |
| | *FTL1a* | Solyc11g008650 | 11 | 2,862,517–2,862,740 | 2,800,001–2,900,000 |
| | *FTL1b* | Solyc11g008640 | 11 | 2,848,285–2,850,649 | 2,800,001–2,900,000 |
| CSHL PAS014479_MAS2.0 known domestication loci | *EID1* | Solyc09g075080 | 09 | 65,168,316–65,171,100 | 65,120,001–65,220,000 |
| | *LNK2* | Solyc01g068560 | 01 | 73,525,568–73,539,374 | 73,480,001–73,580,000 |
| | *lc*/*SlWUS* | Solyc02g083950 | 02 | 45,712,924–45,714,360 | 45,660,001–45,760,000 |
| | *OVATE* | Solyc02g085500 | 02 | 46,873,736–46,875,468 | 46,820,001–46,920,000 |
| | *fw2.2*/*CNR* | Solyc02g090730 | 02 | 50,829,171–50,829,965 | 50,780,001–50,880,000 |
| | *LIN5* | Solyc09g010080 | 09 | 3,513,231–3,517,348 | 3,470,001–3,570,000 |
| | *FAS*/*SlCLV3* | Solyc11g071380 | 11 | 52,957,700–52,958,260 | 52,910,001–53,010,000 |
| | *GAME12* | Solyc12g006470 | 12 | 975,445–981,593 | 930,001–1,030,000 |
| Li et al. BGV1.3 target loci | *SP5G* | Solyc05g053850 | 05 | 64,963,662–64,965,436 | 64,910,001–65,010,000 |
| | *SP6A* | Solyc05g055660 | 05 | 66,276,035–66,277,974 | 66,230,001–66,330,000 |
| | *SFT* | Solyc03g063100 | 03 | 29,450,608–29,454,988 | 29,400,001–29,500,000 |
| | *FTL1a* | Solyc11g008650 | 11 | — | 2,820,001–2,920,000 |
| | *FTL1b* | Solyc11g008640 | 11 | 2,871,487–2,873,877 | 2,820,001–2,920,000 |
| Li et al. BGV1.3 known domestication loci | *EID1* | Solyc09g075080 | 09 | 64,509,004–64,512,203 | 64,460,001–64,560,000 |
| | *LNK2* | Solyc01g068560 | — | — | — |
| | *lc*/*SlWUS* | Solyc02g083950 | 02 | 44,641,381–44,644,944 | 44,590,001–44,690,000 |
| | *OVATE* | Solyc02g085500 | 02 | 45,827,412–45,828,828 | 45,780,001–45,880,000 |
| | *fw2.2*/*CNR* | Solyc02g090730 | 02 | 49,657,477–49,660,008 | 49,610,001–49,710,000 |
| | *LIN5* | Solyc09g010080 | 09 | 3,587,956–3,592,095 | 3,540,001–3,640,000 |
| | *FAS*/*SlCLV3* | Solyc11g071380 | — | — | — |
| | *GAME12* | Solyc12g006470 | 12 | 979,223–985,323 | 930,001–1,030,000 |

—, no directly validated gene model in the assembly; no coordinate was inferred.

[a] 100-kb window whose center lies nearest the gene center.

**Supplementary Table 3.** Published sweep evidence at known domestication loci.

| Locus | Gene ID | Coordinates (BGV1.3) | Trait | Sweep evidence | References |
|---|---|---|---|---|---|
| *EID1* | Solyc09g075080 | 9:65214817–65217601 | Circadian rhythm | High $\pi_{SP}/\pi_{SLC}$ | [35, 111] |
| *LNK2* | Solyc01g068560 | 1:71067359–71081167 | Circadian rhythm | High $\pi_{SP}/\pi_{SLC}$ | [109] |
| *lc/SlWUS* | Solyc02g083950 | 2:46830023–46831448 | Fruit size | High $\pi_{SP}/\pi_{SLC}$ | [35, 112] |
| *OVATE* | Solyc02g085500 | 2:48027328–48029060 | Fruit shape | CLR, mean $F_{ST}$, high $\pi_{SP}/\pi_{SLC}$ | [34, 113] |
| *fw2.2/CNR* | Solyc02g090730 | 2:51859918–51860703 | Fruit size | High $\pi_{SP}/\pi_{SLC}$, negative Tajima's *D*, XP-CLR | [114, 115] |
| *LIN5* | Solyc09g010080 | 9:3620212–3624362 | Sucrose metabolism | CLR, mean $F_{ST}$, high $\pi_{SP}/\pi_{SLC}$ | [34, 116] |
| *FAS/SlCLV3* | Solyc11g071380 | 11:52977003–52977557 | Fruit size, locule number | High $\pi_{SP}/\pi_{SLC}$, XP-CLR | [34, 117] |
| *GAME12* | Solyc12g006470 | 12:963842–969905 | Alkaloid biosynthesis | CLR, mean $F_{ST}$, high $\pi_{SP}/\pi_{SLC}$ | [34, 118] |

**Supplementary Table 4.** Simulation parameters.

| Parameter | 83-bp deletion origin of SLC (model 5) | 52-bp deletion redomestication (model 3) |
|---|---|---|
| *Demography* | | |
| Reference size $N_{\mathrm{ref}}$ | 42,560 | 5,694 |
| Split time | 78,148 | 6,967 |
| Onset of secondary contact $T_2$ | 45,956 | 711 |
| Sweeping population, $N_e$ | SLC 11,800[a] | SLL 1,981 |
| Reference population, $N_e$ | SP 6,405 | SLC 28,432 |
| Ancestral sizes | 2,549 and 4,133 | no size change |
| Migration into sweeping pop. | $9.47 \times 10^{-6}$ | $8.66 \times 10^{-5}$ |
| Migration out of sweeping pop. | $3.99 \times 10^{-6}$ | $2.04 \times 10^{-4}$ |
| *Sweep grid* | | |
| Starting frequency $f_0$ | hard, 0.001 to 0.37[b] | hard only |
| Sweep age (generations) | 0 and 45,956 | 0, 500 and 711 |
| Panels (haplotypes) | 15 × 81, 30 × 186 | 10 × 81, 236 × 186 |
| Selection coefficient $s$ | 0.01, 0.05, 0.1 | |
| Final frequency | 0.99 | |
| *Simulation and scoring* | | |
| Generation time | 1 year | |
| Mutation rate $\mu$ | $1 \times 10^{-8}$ per bp per generation | |
| Recombination rate $r$ | $9.5 \times 10^{-10}$ primary[c] | |
| Window size | 100 kb | |
| Simulated region | 500 kb, scored over the central window | |
| Replicates per cell | up to 2,000 msprime, 500 discoal[d] | |
| Neutral null | 2,000 windows, 5,000 for the 10-kb calibration | |
| Detection threshold | matched neutral 5% tail | |

Sizes are diploid individuals, times are generations.

[a] Recalibrated. The fitted value, 211,649, describes SLC and SLL pooled.

[b] hard, 0.001, 0.005, 0.01, 0.02, 0.05, 0.10, 0.37.

[c] Effective rate under 95% selfing.

[d] Fewer contribute where a statistic is undefined.

**Supplementary Table 5.** Detection power, within-population statistics, within SLC.

| | $f_0$[a] | $\pi$ | Tajima's $D$ | $nS_L$ |
|---|---|---|---|---|
| matched to CSHL SLC ($n$ = 81) | hard sweep | **0.9995** | **0.9910** | 0.4887 |
| | 0.001 | **0.7900** | 0.2055 | **0.5124** |
| | 0.005 | 0.2375 | 0.0020 | 0.2140 |
| | 0.01 | 0.1255 | 0.0035 | 0.1260 |
| | 0.02 | 0.0735 | 0.0045 | 0.0720 |
| | 0.05 | 0.0710 | 0.0145 | 0.0420 |
| | 0.10 | 0.0410 | 0.0215 | 0.0720 |
| | 0.37 | 0.0540 | 0.0360 | 0.0360 |
| matched to Li et al. SLC ($n$ = 186) | hard sweep | **1.0000** | **0.9965** | **0.5522** |
| | 0.001 | **0.8025** | 0.2000 | 0.4640 |
| | 0.005 | 0.2170 | 0.0000 | 0.2400 |
| | 0.01 | 0.1035 | 0.0020 | 0.1620 |
| | 0.02 | 0.0990 | 0.0040 | 0.1200 |
| | 0.05 | 0.0560 | 0.0100 | 0.0560 |
| | 0.10 | 0.0485 | 0.0115 | 0.0560 |
| | 0.37 | 0.0590 | 0.0385 | 0.0440 |

Fraction of sweep replicates beyond the 5% tail of a matched neutral set. $s = 0.05$, 100-kb window, $r_{eff} = 9.5 \times 10^{-10}$ (95% selfing); 2,000 replicates for $\pi$ and Tajima's $D$, 500 for $nS_L$. **Bold**, power above 0.5.

Single-population statistics; migration does not apply.

[a] Starting frequency; *hard sweep* is a single copy. The 83-bp deletion is at 0.37 in SP.

**Supplementary Table 6**. Detection power, cross-population statistics, SLC against SP.

| | $f_0$[a] | $\pi$ ratio | | XP-$nS_L$ | |
|---|---|---|---|---|---|
| | | $\tau = 0$ | $\tau = 45{,}956$[b] | $\tau = 0$ | $\tau = 45{,}956$[b] |
| matched to CSHL SLC ($n = 81$) vs. SP ($n = 15$) | hard sweep | **0.9660** | 0.0520 | **1.0000** | 0.0600 |
| | 0.001 | **0.9580** | 0.0680 | **1.0000** | 0.0580 |
| | 0.005 | **0.9400** | 0.0600 | **0.9495** | 0.0520 |
| | 0.01 | **0.9100** | 0.0600 | **0.8793** | 0.0700 |
| | 0.02 | **0.8320** | 0.0440 | **0.7280** | 0.0460 |
| | 0.05 | **0.7060** | 0.0600 | 0.3988 | 0.0500 |
| | 0.10 | 0.4940 | 0.0740 | 0.1002 | 0.0840 |
| | 0.37 | 0.1040 | 0.0520 | 0.0020 | 0.0560 |
| matched to Li et al. SLC ($n = 186$) vs. SP ($n = 30$) | hard sweep | **0.9700** | 0.0640 | **1.0000** | 0.0380 |
| | 0.001 | **0.9620** | 0.0460 | **1.0000** | 0.0540 |
| | 0.005 | **0.9720** | 0.0680 | **0.9899** | 0.0320 |
| | 0.01 | **0.9180** | 0.0440 | **0.9758** | 0.0640 |
| | 0.02 | **0.8600** | 0.0400 | **0.9069** | 0.0560 |
| | 0.05 | **0.6940** | 0.0740 | **0.6861** | 0.0320 |
| | 0.10 | 0.4580 | 0.0560 | 0.3212 | 0.0540 |
| | 0.37 | 0.0960 | 0.0540 | 0.0040 | 0.0500 |

Fraction of sweep replicates beyond the 5% tail of a matched neutral set. $s = 0.05$, 100-kb window, $r_{\text{eff}} = 9.5 \times 10^{-10}$ (95% selfing); 500 replicates per cell. **Bold**, power above 0.5.

Migration on; both settings were run.

[a] Starting frequency; *hard sweep* is a single copy. The 83-bp deletion is at 0.37 in SP.

[b] Sweep age in generations; $T_2 = 45{,}956$ is the age of a sweep at the origin of SLC.

**Supplementary Table 7.** Detection power for both deletions.

| | $f_0$ | age | $\pi$ | Tajima's *D* | $nS_L$ | $\pi$ ratio | XP-$nS_L$ |
|---|---|---|---|---|---|---|---|
| *83 bp deletion, SLC against SP, migration on* | | | | | | | |
| CSHL SLC ($n$ = 81) vs. SP ($n$ = 15) | hard | 0 | **0.9995** | **0.9910** | 0.4887 | **0.9660** | **1.0000** |
| | hard | 45,956 | **0.9995** | **0.9910** | 0.4887 | 0.0520 | 0.0600 |
| | 0.37 | 0 | 0.0540 | 0.0360 | 0.0360 | 0.1040 | 0.0020 |
| | 0.37 | 45,956 | 0.0540 | 0.0360 | 0.0360 | 0.0520 | 0.0560 |
| Li et al. SLC ($n$ = 186) vs. SP ($n$ = 30) | hard | 0 | **1.0000** | **0.9965** | **0.5522** | **0.9700** | **1.0000** |
| | hard | 45,956 | **1.0000** | **0.9965** | **0.5522** | 0.0640 | 0.0380 |
| | 0.37 | 0 | 0.0590 | 0.0385 | 0.0440 | 0.0960 | 0.0040 |
| | 0.37 | 45,956 | 0.0590 | 0.0385 | 0.0440 | 0.0540 | 0.0500 |
| *52 bp deletion, SLL against SLC, no migration* | | | | | | | |
| CSHL SLL ($n$ = 10) vs. SLC ($n$ = 81) | hard | 0 | **0.9545** | 0.2936 | n/a | **0.9460** | **0.9980** |
| | hard | 711 | **0.9545** | 0.2936 | n/a | **0.8580** | 0.4740 |
| Li et al. SLL ($n$ = 236) vs. SLC ($n$ = 186) | hard | 0 | **0.9960** | **0.9880** | 0.0843 | **0.9720** | **1.0000** |
| | hard | 711 | **0.9960** | **0.9880** | 0.0843 | **0.8880** | **0.8460** |

$s$ = 0.05, 100-kb windows, 95% selfing. **Bold**, power above 0.5.

n/a, $nS_L$ not computable for the SLL panel of ten.

**Supplementary Table 8.** Within-population statistics at target loci.

| | Locus | $\pi$ (× $10^{-3}$) | Tajima's $D$[a] | $nS_L$[b] |
|---|---|---|---|---|
| CSHL SP ($n$ = 15) | *SP5G* | 3.597 (45.2) | 0.30 (35.6) | 0.0176 (23.3) |
| | *SP6A* | 3.299 (36.3) | 0.94 (71.8) | 0.0459 (65.3) |
| | *FTL1a/b* | 3.933 (54.7) | 0.93 (71.3) | **0.1260** (**96.0**) |
| | *SFT* | 1.213 ( 2.0) | **−0.84** ( **3.5**) | 0.0584 (75.8) |
| CSHL SLC ($n$ = 81) | *SP5G* | 1.527 (24.0) | −1.31 (10.8) | 0.0026 ( 3.9) |
| | *SP6A* | 2.134 (46.8) | −1.67 ( 5.5) | 0.0194 (39.6) |
| | *FTL1a/b* | 2.338 (53.6) | 0.01 (60.7) | 0.0177 (36.2) |
| | *SFT* | 2.271 (51.5) | 1.03 (90.3) | 0.0240 (48.1) |
| CSHL SLL ($n$ = 10) | *SP5G* | 1.118 (80.2) | **−0.75** ( **4.6**) | 0.0254 (56.5) |
| | *SP6A* | 1.986 (94.3) | 0.70 (41.0) | 0.0211 (48.8) |
| | *FTL1a/b* | 0.326 (17.2) | −0.04 (14.2) | 0.0000 ( 9.1) |
| | *SFT* | 0.368 (21.3) | 0.52 (31.7) | 0.0132 (32.5) |
| Li et al. SP ($n$ = 30) | *SP5G* | 0.695 (13.2) | 2.25 (62.7) | 0.0135 (39.3) |
| | *SP6A* | 0.928 (25.9) | **3.55** (**96.5**) | 0.0439 (73.3) |
| | *FTL1a/b* | 1.147 (38.7) | 2.54 (72.6) | **0.1707** (**96.1**) |
| | *SFT* | 0.518 ( 6.3) | **−0.76** ( **2.4**) | 0.0000 ( 8.4) |
| Li et al. SLC ($n$ = 186) | *SP5G* | 0.502 ( 8.6) | 1.45 (39.4) | 0.0816 (86.6) |
| | *SP6A* | 0.605 (13.7) | 1.44 (39.2) | 0.0640 (82.4) |
| | *FTL1a/b* | 0.700 (18.8) | 1.20 (30.4) | 0.1111 (90.8) |
| | *SFT* | 1.267 (47.3) | 2.60 (78.1) | 0.0324 (66.0) |
| Li et al. SLL ($n$ = 236) | *SP5G* | 0.224 (10.8) | −0.45 (25.8) | — |
| | *SP6A* | 0.124 ( 4.3) | **−2.17** ( **2.5**) | — |
| | *FTL1a/b* | 0.030 ( 0.5) | **−2.54** ( **0.4**) | — |
| | *SFT* | 0.599 (43.9) | 0.34 (39.2) | 0.0179 (51.5) |

Value (genome-wide percentile); 100-kb windows, 10-kb step. **Bold**, percentile above 95, or below 5 for Tajima's $D$. —, focal window not rankable.

[a] [100]. Negative values indicate an excess of low-frequency variants, as expected under a recent sweep.

[b] [101, 103]. Fraction of steps with |normalized $nS_L$| > 2; background excludes chromosome 5. For SLL, sites come from the SLC× SLL pairing.

**Supplementary Table 9.** Cross-population statistics at target loci, SLC against SP.

| | Locus | $F_{ST}$[a] Weir–Cockerham | Hudson | $\pi$ ratio[b] | XP-$nS_L$[c] longer in SLC | longer in SP |
|---|---|---|---|---|---|---|
| CSHL SP ($n$ = 15) vs. SLC ($n$ = 81) | *SP5G* | 0.5349 (90.4) | 0.4548 (88.3) | 2.356 (82.4) | 0.0245 (77.2) | 0.0000 ( 6.2) |
| | *SP6A* | 0.2746 (28.7) | 0.2492 (29.5) | 1.546 (44.0) | 0.0198 (74.8) | 0.0008 (14.8) |
| | *FTL1a/b* | 0.3952 (59.3) | 0.3555 (61.6) | 1.682 (54.0) | 0.0262 (77.9) | 0.0025 (21.5) |
| | *SFT* | 0.4759 (79.0) | **0.5319** (**97.5**) | 0.534 ( 0.4) | 0.0000 (18.0) | 0.0861 (92.0) |
| Li et al. SP ($n$ = 30) vs. SLC ($n$ = 186) | *SP5G* | 0.5487 (84.1) | 0.5197 (83.0) | 1.384 (67.4) | 0.0707 (86.8) | 0.0303 (66.1) |
| | *SP6A* | 0.4336 (59.1) | 0.3958 (53.0) | 1.534 (75.7) | **0.4526** (**98.2**) | 0.0000 (16.3) |
| | *FTL1a/b* | 0.5411 (83.1) | 0.4949 (78.2) | 1.640 (80.9) | **0.3553** (**97.1**) | 0.0000 (16.3) |
| | *SFT* | 0.5592 (85.6) | **0.6255** (**95.7**) | 0.409 ( 1.5) | 0.0000 (36.5) | **0.3321** (**99.7**) |

Value (genome-wide percentile); 100-kb windows, 10-kb step. **Bold**, percentile above 95.

[a] Weir–Cockerham [97]; Hudson [98].

[b] $\pi_{SP}/\pi_{SLC}$.

[c] [103, 119]. One run per contrast with the more derived population as query, reported as its two tails; background excludes chromosome 5.

**Supplementary Table 10.** Cross-population statistics at target loci, SLL against SP and SLC.

| | Locus | $F_{ST}$[a] | | $\pi$ ratio[b] | XP-$nS_L$[c] | |
|---|---|---|---|---|---|---|
| | | Weir–Cockerham | Hudson | | longer in SLL | longer in the other |
| CSHL SP ($n$ = 15) vs. SLL ($n$ = 10) | *SP5G* | 0.4622 (53.4) | 0.5110 (52.7) | 3.204 (20.4) | 0.0031 (58.3) | 0.0046 (26.0) |
| | *SP6A* | 0.2206 ( 1.7) | 0.2590 ( 2.0) | 1.668 ( 3.6) | 0.0000 (20.4) | 0.0082 (34.7) |
| | *FTL1a/b* | 0.5245 (76.4) | 0.5858 (79.8) | 12.217 (84.1) | 0.0000 (20.4) | 0.0008 (13.6) |
| | *SFT* | **0.8100 (99.5)** | **0.8300 (99.5)** | 3.297 (21.6) | 0.0000 (20.4) | 0.0753 (89.1) |
| CSHL SLC ($n$ = 81) vs. SLL ($n$ = 10) | *SP5G* | 0.0278 (14.1) | 0.0601 ( 9.3) | 1.366 ( 5.1) | 0.0000 (29.0) | 0.0106 (48.4) |
| | *SP6A* | 0.0116 ( 3.5) | 0.0352 ( 3.5) | 1.077 ( 1.1) | 0.0000 (29.0) | **0.2151 (98.0)** |
| | *FTL1a/b* | 0.0511 (31.6) | 0.1404 (55.4) | 7.168 (86.1) | 0.0000 (29.0) | 0.0057 (36.1) |
| | *SFT* | 0.1104 (77.7) | 0.2083 (81.5) | 6.179 (81.3) | 0.0000 (29.0) | 0.0024 (24.0) |
| Li et al. SP ($n$ = 30) vs. SLL ($n$ = 236) | *SP5G* | 0.7602 (77.6) | 0.6563 (82.3) | 3.110 (69.5) | 0.0200 (85.0) | 0.0000 (20.5) |
| | *SP6A* | 0.7957 (85.3) | 0.6118 (71.4) | 7.500 (91.0) | **0.7500 (99.4)** | 0.0000 (20.5) |
| | *FTL1a/b* | **0.9009 (98.8)** | 0.7046 (90.8) | **38.880 (99.8)** | **0.7872 (99.5)** | 0.0000 (20.5) |
| | *SFT* | 0.7882 (83.7) | **0.7970 (97.5)** | 0.865 (15.8) | 0.0000 (40.6) | **0.2705 (98.8)** |
| Li et al. SLC ($n$ = 186) vs. SLL ($n$ = 236) | *SP5G* | 0.0672 (49.9) | 0.0657 (49.6) | 2.246 (69.3) | 0.0000 (39.4) | 0.0000 (36.2) |
| | *SP6A* | 0.1342 (83.9) | 0.1266 (83.3) | 4.888 (91.5) | **0.3721 (96.8)** | 0.0000 (36.2) |
| | *FTL1a/b* | 0.1554 (89.8) | 0.1426 (88.4) | **23.708 (99.9)** | 0.0000 (39.4) | 0.0000 (36.2) |
| | *SFT* | 0.0856 (61.4) | 0.0827 (61.1) | 2.115 (66.2) | 0.0000 (39.4) | 0.0000 (36.2) |

Value (genome-wide percentile); 100-kb windows, 10-kb step. **Bold**, percentile above 95.

[a] Weir–Cockerham [97]; Hudson [98].

[b] $\pi$ of the named population over $\pi$ of the query, the less derived population in the numerator.

[c] [103, 119]. One run per contrast with the more derived population as query, reported as its two tails; background excludes chromosome 5.

**Supplementary Table 11.** Within-population statistics at target loci, eggplant.

| | Locus | $\pi$ (× $10^{-3}$) | Tajima's $D$[a] | $nS_L$[b] |
|---|---|---|---|---|
| IND ($n$ = 47) | *SP5G* | 1.612 (78.8) | −0.17 (69.8) | 0.0906 (89.6) |
| | *SP6A* | 0.918 (60.7) | −1.92 (22.1) | 0.0042 (41.5) |
| | *FTL1a* | 1.412 (75.2) | 0.84 (86.8) | 0.0702 (86.0) |
| | *FTL1b* | 1.468 (76.3) | 0.82 (86.6) | 0.0784 (87.7) |
| | *SFT* | 0.649 (49.2) | −0.39 (63.9) | 0.0185 (61.4) |
| MEU ($n$ = 121) | *SP5G* | 1.325 (85.0) | −0.35 (83.6) | 0.0124 (41.4) |
| | *SP6A* | 0.784 (71.1) | −1.82 (55.6) | 0.0249 (54.7) |
| | *FTL1a* | 1.245 (83.6) | 0.42 (90.7) | 0.0000 (13.0) |
| | *FTL1b* | 1.322 (84.9) | 0.46 (91.0) | 0.0129 (42.0) |
| | *SFT* | 0.396 (46.9) | −1.68 (59.4) | 0.0000 (13.0) |

Value (genome-wide percentile); 100-kb windows, 10-kb step. **Bold**, percentile above 95, or below 5 for Tajima's *D*.

[a] [100]. Negative values indicate an excess of low-frequency variants, as expected under a recent sweep.

[b] [101, 103]. Fraction of steps with |normalized $nS_L$| > 2.

**Supplementary Table 12.** Cross-population statistics at target loci, eggplant.

| Locus | $F_{ST}$ | | $\pi$ ratio[b] | XP-$nS_L$[c] | |
|---|---|---|---|---|---|
| | Weir–Cockerham | Hudson[a] | | longer in IND | longer in MEU |
| *SP5G* | 0.0338 (20.5) | 0.0380 (20.5) | 1.216 (37.7) | 0.0000 (31.8) | 0.0080 (71.9) |
| *SP6A* | 0.0039 ( 1.7) | 0.0080 ( 1.4) | 1.172 (35.6) | 0.0023 (75.9) | 0.0000 (29.6) |
| *FTL1a* | 0.0114 ( 4.3) | 0.0166 ( 4.4) | 1.135 (33.8) | 0.0000 (31.8) | 0.0037 (67.3) |
| *FTL1b* | 0.0106 ( 4.0) | 0.0161 ( 4.2) | 1.111 (32.6) | 0.0000 (31.8) | 0.0036 (67.0) |
| *SFT* | 0.1259 (69.3) | 0.1168 (70.2) | 1.640 (58.0) | 0.0000 (31.8) | 0.0089 (72.6) |

IND ($n$ = 47) against MEU ($n$ = 121). Value (genome-wide percentile); 100-kb windows, 10-kb step. **Bold**, percentile above 95.

[a] Weir–Cockerham [97]; Hudson [98].

[b] $\pi_{IND}/\pi_{MEU}$.

[c] [103, 119]. One run, reported as its two tails.

**Supplementary Table 13.** Within-population statistics at known domestication loci.

| | Locus | $\pi$ (× $10^{-3}$) | Tajima's $D$[a] | $nS_L$[b] |
|---|---|---|---|---|
| CSHL SP ($n$ = 15) | *EID1* | 4.821 (73.4) | 0.57 (50.0) | **0.1215** (**95.5**) |
| | *LNK2* | 2.623 (17.9) | 0.54 (48.3) | 0.0146 (17.9) |
| | *lc*/*SlWUS* | 2.958 (26.1) | 0.71 (58.7) | 0.0644 (79.5) |
| | *OVATE* | 2.210 (10.5) | 0.74 (60.2) | 0.0252 (37.3) |
| | *fw2.2*/*CNR* | 2.786 (21.6) | 0.95 (72.3) | 0.0177 (23.5) |
| | *LIN5* | 3.795 (51.0) | 0.68 (56.6) | 0.0341 (51.2) |
| | *FAS*/*SlCLV3* | 4.581 (69.1) | 0.63 (53.2) | 0.0564 (74.4) |
| | *GAME12* | 5.273 (79.6) | 1.28 (88.3) | 0.0582 (75.7) |
| CSHL SLC ($n$ = 81) | *EID1* | 2.537 (59.7) | −0.28 (47.2) | 0.0190 (38.7) |
| | *LNK2* | 1.012 ( 8.2) | −1.06 (16.5) | **0.3431** (**99.3**) |
| | *lc*/*SlWUS* | 1.403 (19.6) | −0.10 (55.7) | 0.0026 ( 3.9) |
| | *OVATE* | 1.287 (15.8) | −0.20 (50.8) | 0.0090 (16.3) |
| | *fw2.2*/*CNR* | 1.894 (38.0) | 0.45 (76.1) | 0.0216 (44.0) |
| | *LIN5* | 1.713 (31.1) | −0.65 (30.3) | 0.0088 (15.9) |
| | *FAS*/*SlCLV3* | 2.884 (69.1) | 0.59 (80.5) | 0.0116 (22.4) |
| | *GAME12* | 2.950 (70.6) | 1.23 (93.7) | 0.0501 (76.2) |
| CSHL SLL ($n$ = 10) | *EID1* | 0.472 (31.8) | 0.19 (19.8) | 0.0123 (30.9) |
| | *LNK2* | 0.107 ( 1.7) | 0.18 (19.4) | — |
| | *lc*/*SlWUS* | 0.181 ( 5.7) | 0.98 (58.9) | 0.0625 (86.1) |
| | *OVATE* | 0.435 (28.0) | **2.17** (**95.9**) | 0.0545 (83.2) |
| | *fw2.2*/*CNR* | 0.164 ( 4.6) | 1.66 (89.5) | **0.1429** (**95.3**) |
| | *LIN5* | 0.237 ( 9.7) | 0.34 (24.5) | 0.0222 (50.9) |
| | *FAS*/*SlCLV3* | 0.862 (67.0) | 1.52 (86.1) | 0.0820 (90.3) |
| | *GAME12* | 1.497 (89.8) | 0.48 (29.9) | **0.2351** (**97.9**) |
| Li et al. SP ($n$ = 30) | *EID1* | 1.397 (51.0) | 2.46 (69.6) | 0.0187 (48.1) |
| | *LNK2* | — | — | — |
| | *lc*/*SlWUS* | 1.152 (39.0) | 2.39 (67.2) | 0.0000 ( 8.4) |
| | *OVATE* | 0.996 (29.9) | 2.66 (76.8) | 0.1439 (94.4) |
| | *fw2.2*/*CNR* | 0.999 (30.2) | 3.11 (90.0) | 0.0598 (80.5) |
| | *LIN5* | 1.368 (49.8) | 2.52 (71.9) | 0.0256 (58.1) |
| | *FAS*/*SlCLV3* | — | — | — |
| | *GAME12* | 1.801 (65.9) | **3.67** (**97.5**) | 0.0000 ( 8.4) |
| | *EID1* | 1.148 (41.6) | 2.19 (64.1) | 0.0543 (79.2) |

| | | | | |
|---|---|---|---|---|
| Li et al. SLC ($n$ = 186) | *LNK2* | — | — | — |
| | *lc*/*SlWUS* | 0.830 (25.8) | 1.70 (48.5) | 0.1075 (90.4) |
| | *OVATE* | 0.635 (15.4) | 1.41 (38.0) | **0.3984** (**99.0**) |
| | *fw2.2*/*CNR* | 0.802 (24.3) | 2.52 (75.7) | 0.0392 (71.2) |
| | *LIN5* | 0.810 (24.7) | 1.06 (26.6) | **0.1823** (**95.5**) |
| | *FAS*/*SlCLV3* | — | — | — |
| | *GAME12* | 1.432 (55.1) | 3.26 (92.1) | 0.0000 ( 6.7) |
| Li et al. SLL ($n$ = 236) | *EID1* | 0.420 (28.7) | −0.24 (29.4) | — |
| | *LNK2* | — | — | — |
| | *lc*/*SlWUS* | 0.313 (18.6) | −0.71 (21.3) | — |
| | *OVATE* | 0.216 (10.3) | −1.43 (10.8) | — |
| | *fw2.2*/*CNR* | 0.253 (13.2) | −0.41 (26.5) | 0.0000 (15.1) |
| | *LIN5* | 0.364 (23.4) | −1.25 (13.2) | 0.1000 (92.7) |
| | *FAS*/*SlCLV3* | — | — | — |
| | *GAME12* | 0.351 (22.2) | −1.41 (11.1) | — |

Value (genome-wide percentile); 100-kb windows, 10-kb step. **Bold**, percentile above 95, or below 5 for Tajima's *D*. —, focal window not rankable.

[a] [100]. Negative values indicate an excess of low-frequency variants, as expected under a recent sweep.

[b] [101, 103]. Fraction of steps with |normalized $nS_L$| > 2; background excludes chromosome 5. For SLL, sites come from the SLC× SLL pairing.

**Supplementary Table 14.** Cross-population statistics at known domestication loci, SLC against SP.

| | Locus | $F_{ST}$[a] | | π ratio[b] | XP-$nS_L$[c] | |
|---|---|---|---|---|---|---|
| | | Weir–Cockerham | Hudson | | longer in SLC | longer in SP |
| CSHL SP ($n$ = 15) vs. SLC ($n$ = 81) | *EID1* | 0.3309 (42.3) | 0.2830 (38.8) | 1.900 (67.6) | 0.0128 (70.2) | 0.0070 (34.7) |
| | *LNK2* | 0.4446 (70.7) | 0.3595 (62.7) | 2.591 (86.0) | 0.0276 (78.5) | 0.0000 ( 6.2) |
| | *lc/SlWUS* | 0.3738 (53.5) | 0.3139 (47.9) | 2.108 (76.4) | **0.2747** (**98.2**) | 0.0000 ( 6.2) |
| | *OVATE* | 0.3535 (47.9) | 0.3145 (48.1) | 1.718 (56.6) | 0.0099 (67.3) | 0.0000 ( 6.2) |
| | *fw2.2/CNR* | 0.3521 (47.5) | 0.3271 (52.1) | 1.471 (38.4) | 0.0000 (18.0) | 0.0094 (40.4) |
| | *LIN5* | 0.4291 (67.1) | 0.3597 (62.8) | 2.216 (79.5) | 0.1003 (92.0) | 0.0000 ( 6.2) |
| | *FAS/SlCLV3* | 0.2731 (28.3) | 0.2439 (27.7) | 1.589 (47.1) | 0.0223 (76.1) | 0.0084 (38.2) |
| | *GAME12* | 0.3638 (50.7) | 0.3199 (49.8) | 1.788 (61.2) | 0.0013 (46.0) | 0.0000 ( 6.2) |
| Li et al. SP ($n$ = 30) vs. SLC ($n$ = 186) | *EID1* | 0.3261 (34.2) | 0.3124 (30.7) | 1.217 (57.0) | 0.2000 (93.5) | 0.0077 (45.2) |
| | *LNK2* | — | — | — | — | — |
| | *lc/SlWUS* | 0.3904 (48.6) | 0.3639 (43.8) | 1.389 (67.6) | 0.1543 (91.7) | 0.0000 (16.3) |
| | *OVATE* | 0.4988 (75.3) | 0.4576 (69.7) | 1.569 (77.6) | **0.3510** (**97.0**) | 0.0000 (16.3) |
| | *fw2.2/CNR* | 0.3160 (32.0) | 0.3024 (28.5) | 1.245 (59.0) | 0.0000 (36.5) | 0.0000 (16.3) |
| | *LIN5* | 0.4762 (69.9) | 0.4272 (61.9) | 1.688 (82.6) | **0.5023** (**98.6**) | 0.0000 (16.3) |
| | *FAS/SlCLV3* | — | — | — | — | — |
| | *GAME12* | 0.2275 (15.4) | 0.2170 (13.7) | 1.258 (59.8) | 0.0381 (83.7) | 0.0000 (16.3) |

Value (genome-wide percentile); 100-kb windows, 10kb step. **Bold**, percentile above 95. —, focal window not rankable.

[a] Weir–Cockerham [97]; Hudson [98].

[b] $\pi_{SP}/\pi_{SLC}$.

[c] [103, 119]. One run per contrast with the more derived population as query, reported as its two tails; background excludes chromosome 5.

**Supplementary Table 15.** Cross-population statistics at known domestication loci, SLL against SP and SLC.

| | Locus | $F_{ST}$[a] | | $\pi$ ratio[b] | XP-$nS_L$[c] | |
|---|---|---|---|---|---|---|
| | | Weir–Cockerham | Hudson | | longer in SLL | longer in the other |
| CSHL SP ($n$ = 15) vs. SLL ($n$ = 10) | *EID1* | 0.4233 (41.2) | 0.4858 (44.6) | 10.295 (78.5) | 0.0000 (20.4) | 0.0137 (45.5) |
| | *LNK2* | 0.4596 (52.6) | 0.5280 (58.5) | **23.262** (**96.7**) | 0.0012 (48.4) | 0.0000 ( 5.9) |
| | *lc*/*SlWUS* | 0.4352 (44.9) | 0.5042 (50.4) | 16.557 (91.8) | 0.0319 (88.7) | 0.0000 ( 5.9) |
| | *OVATE* | 0.4623 (53.4) | 0.5190 (55.4) | 5.073 (44.2) | 0.0000 (20.4) | 0.0095 (37.5) |
| | *fw2.2*/*CNR* | 0.5422 (82.3) | 0.6030 (85.0) | 16.956 (92.2) | 0.0000 (20.4) | 0.0000 ( 5.9) |
| | *LIN5* | 0.4718 (56.7) | 0.5359 (61.4) | 16.043 (91.2) | **0.1023** (**96.9**) | 0.0008 (13.3) |
| | *FAS*/*SlCLV3* | 0.4084 (36.5) | 0.4587 (35.6) | 5.328 (47.0) | 0.0191 (82.7) | 0.0024 (19.4) |
| | *GAME12* | 0.4040 (35.0) | 0.4576 (35.2) | 3.512 (24.4) | 0.0000 (20.4) | 0.0007 (12.8) |
| CSHL SLC ($n$ = 81) vs. SLL ($n$ = 10) | *EID1* | 0.0841 (65.9) | 0.1705 (71.9) | 5.365 (75.9) | 0.0011 (62.3) | 0.0064 (38.1) |
| | *LNK2* | 0.0274 (13.8) | 0.1096 (36.4) | 9.023 (91.6) | 0.0000 (29.0) | 0.0149 (56.5) |
| | *lc*/*SlWUS* | 0.0524 (32.9) | 0.1423 (56.6) | 7.738 (88.1) | 0.0000 (29.0) | 0.0000 ( 7.0) |
| | *OVATE* | 0.0767 (60.2) | 0.1490 (61.0) | 2.958 (45.3) | 0.0000 (29.0) | **0.2718** (**98.7**) |
| | *fw2.2*/*CNR* | 0.1205 (80.9) | 0.2350 (86.7) | **11.574** (**95.6**) | 0.0211 (87.3) | 0.0000 ( 7.0) |
| | *LIN5* | 0.0489 (29.4) | 0.1339 (51.1) | 7.218 (86.3) | 0.0000 (29.0) | 0.0000 ( 7.0) |
| | *FAS*/*SlCLV3* | 0.0715 (55.5) | 0.1366 (52.8) | 3.347 (53.1) | 0.0000 (29.0) | 0.0045 (32.3) |
| | *GAME12* | 0.0289 (14.7) | 0.0773 (16.9) | 1.972 (20.3) | 0.0000 (29.0) | 0.0048 (33.3) |
| Li et al. SP ($n$ = 30) vs. SLL ($n$ = 236) | *EID1* | 0.6605 (56.7) | 0.5298 (48.6) | 3.325 (71.9) | **0.3556** (**96.4**) | 0.0000 (20.5) |
| | *LNK2* | — | — | — | — | — |
| | *lc*/*SlWUS* | 0.6931 (63.3) | 0.5563 (55.8) | 3.676 (75.4) | 0.1000 (90.5) | 0.0000 (20.5) |
| | *OVATE* | 0.7673 (79.3) | 0.6230 (74.4) | 4.602 (81.8) | **1.0000** (**99.96**) | 0.0000 (20.5) |
| | *fw2.2*/*CNR* | 0.6929 (63.2) | 0.5481 (53.5) | 3.953 (77.7) | 0.0238 (85.5) | 0.0000 (20.5) |
| | *LIN5* | 0.7040 (65.4) | 0.5666 (58.7) | 3.757 (76.1) | **0.5610** (**98.4**) | 0.0000 (20.5) |
| | *FAS*/*SlCLV3* | — | — | — | — | — |
| | *GAME12* | 0.6636 (57.4) | 0.4836 (36.3) | 5.128 (84.4) | 0.0513 (88.0) | 0.0000 (20.5) |
| Li et al. SLC ($n$ = 186) vs. SLL ($n$ = 236) | *EID1* | 0.1026 (70.8) | 0.0982 (70.4) | 2.733 (77.9) | 0.1538 (93.5) | 0.0000 (36.2) |
| | *LNK2* | — | — | — | — | — |
| | *lc*/*SlWUS* | 0.0849 (61.0) | 0.0824 (60.9) | 2.647 (76.7) | 0.0000 (39.4) | 0.0000 (36.2) |
| | *OVATE* | 0.0812 (58.8) | 0.0785 (58.5) | 2.933 (80.4) | 0.0000 (39.4) | 0.0000 (36.2) |
| | *fw2.2*/*CNR* | 0.1376 (85.0) | 0.1318 (85.2) | 3.174 (82.8) | 0.0541 (89.4) | 0.0000 (36.2) |
| | *LIN5* | 0.0604 (44.7) | 0.0594 (44.6) | 2.225 (68.8) | 0.0299 (87.0) | 0.0000 (36.2) |
| | *FAS*/*SlCLV3* | — | — | — | — | — |
| | *GAME12* | 0.1799 (94.1) | 0.1706 (94.1) | 4.077 (88.4) | 0.1613 (93.6) | 0.0000 (36.2) |

Value (genome-wide percentile); 100-kb windows, 10-kb step. **Bold**, percentile above 95. —, focal window not rankable.

[a] Weir–Cockerham [97]; Hudson [98].

[b] $\pi$ of the named population over $\pi$ of the query, the less derived population in the numerator.

[c] [103, 119]. One run per contrast with the more derived population as query, reported as its two tails; background excludes chromosome 5.

### Supplementary Figures

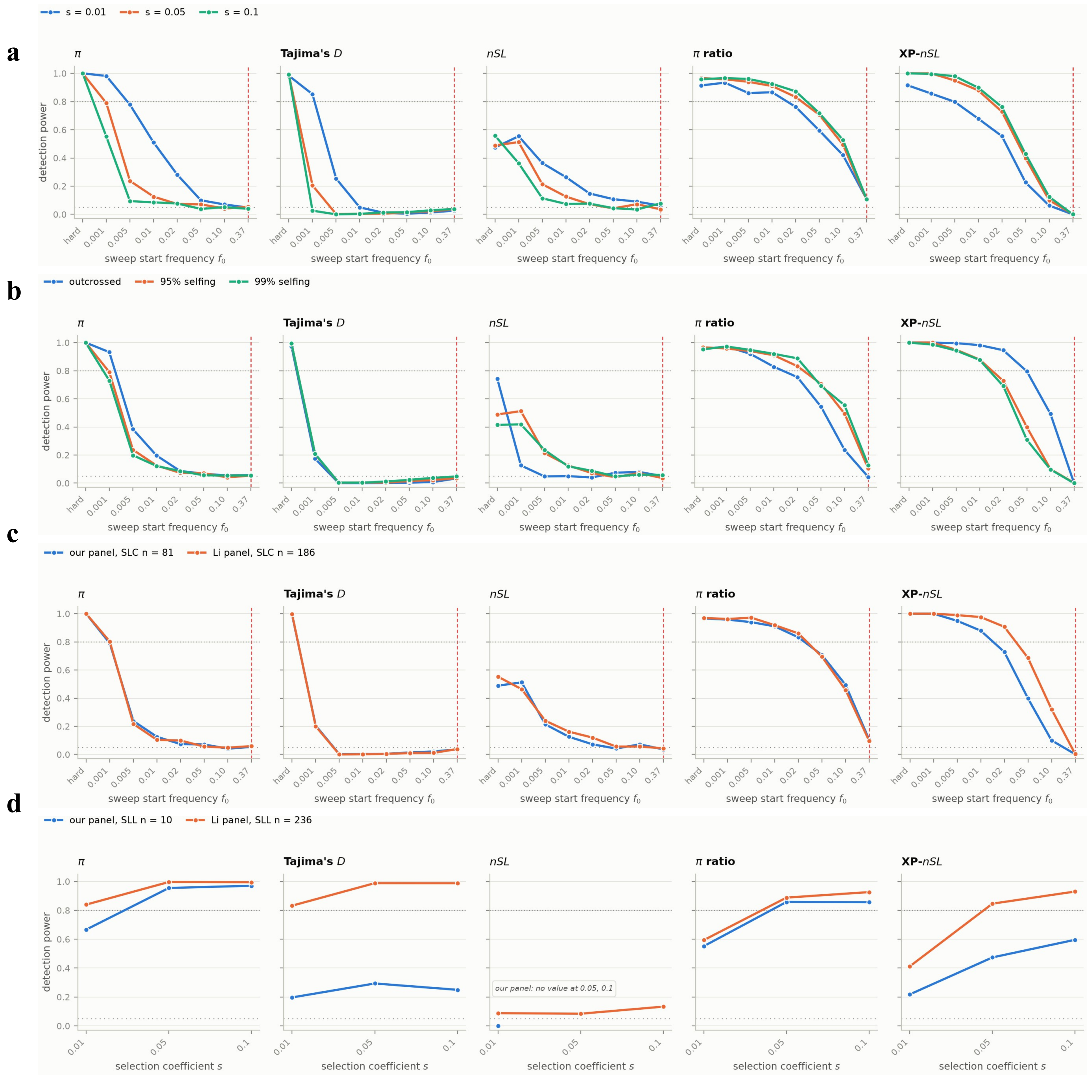


**Supplementary Figure 1. Detection power.** Power is the fraction of sweep replicates beyond the 5% tail of matched neutral simulations, in 100-kb windows. Dashed red marks $f_0 = 0.37$ and dotted gray 80% power. **(a)** Against starting frequency, at 95% selfing, SP 15 × SLC 81. Lines are selection coefficients. (**b)** As (a), at $s = 0.05$, with lines for the outcrossed rate and the effective rates under 95% and 99% selfing. **(c)**, As (a), at $s = 0.05$, with lines for the CSHL panel (SLC $n = 81$) and the Li et al. panel (SLC $n = 186$). **(d)** The 52-bp deletion, SLL against SLC. The sweep is hard, so power is shown against the selection coefficient. The $\pi$ ratio and XP-$nS_L$ are at the re-domestication age of 711 generations. Lines are SLL $n = 10$ against SLC $n = 81$ and SLL $n = 236$ against SLC $n = 186$.

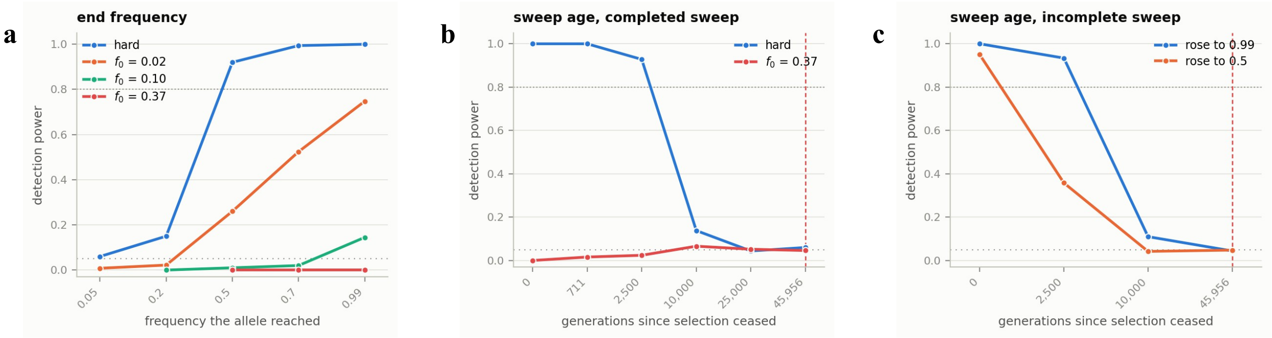


**Supplementary Figure 2. Detection power against end frequency and sweep age.** Power is the fraction of sweep replicates beyond the 5% tail of matched neutral simulations. XP-$nS_L$ only, in the SP–SLC comparison, at $s$ = 0.05, 95% selfing, migration on, 100-kb windows and the CSHL panel. Dotted gray lines mark 80% power and the 5% false-positive rate. **(a)** Power against the frequency reached by the selected allele, scored immediately after selection ceased. Lines are starting frequencies, and each curve begins where the end frequency first exceeds the starting frequency. (**b**) Power against time since selection ceased for a completed sweep, with end frequency 0.99. **(c)** Power against time since selection ceased from a single-copy start, comparing a sweep stopped at 0.50 with one that reached 0.99. Dashed red lines in **b** and **c** mark $T_2 = 45{,}956$ generations, the age a sweep at the origin of SLC would have.

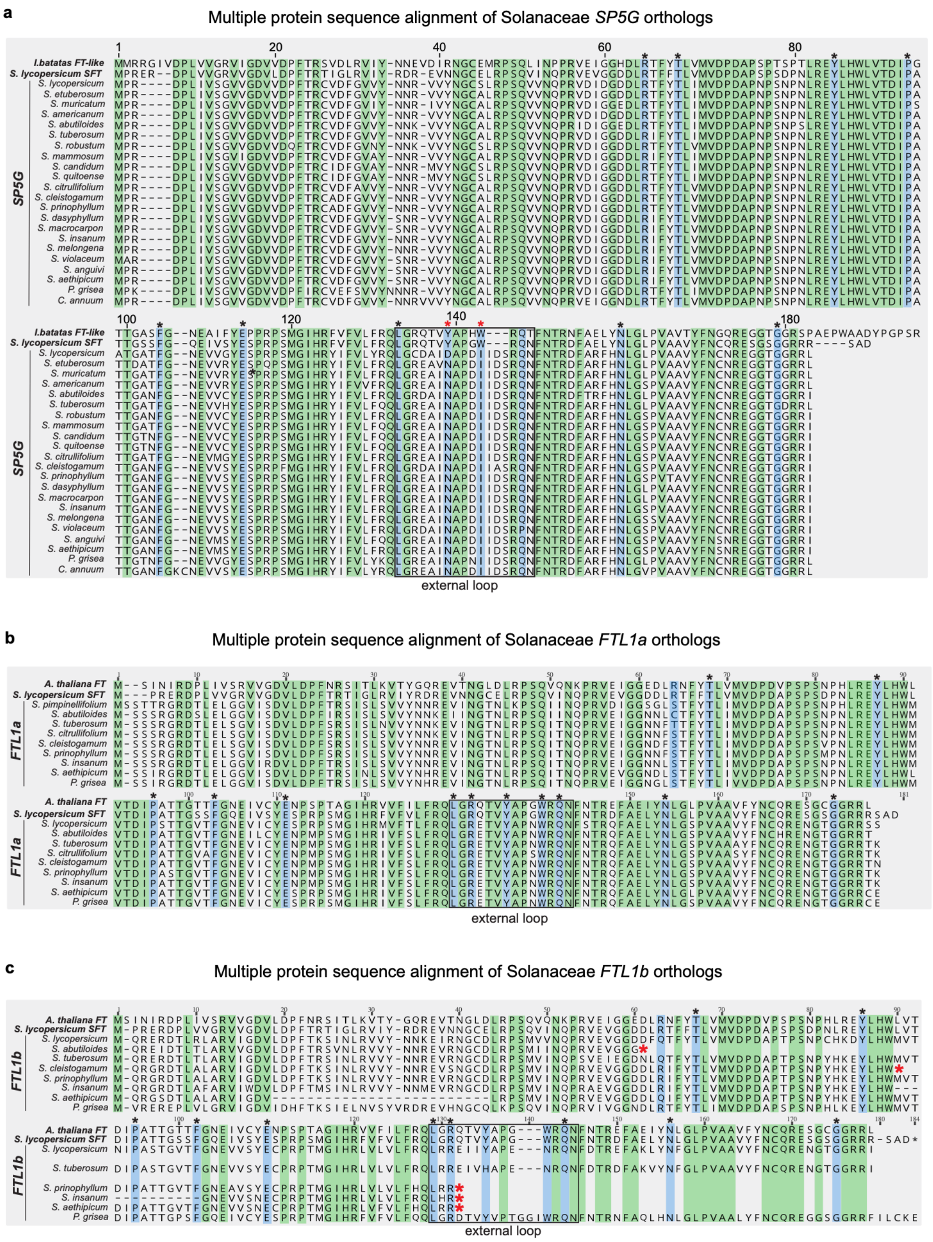


**Supplementary Figure 3. Key florigenic residues are largely conserved across Solanaceae FT-like paralogues. (a-c)** Full protein alignments of Solanaceae *SP5G* (a), *FTL1a* (b), and *FTL1b* (c) orthologues. The tomato FT (SFT) and the closest sweet potato (*Ipomoea batatas*) homolog retain the canonical *florigen* motif in the external loop domain. Asterisks and blue color mark key florigenic amino acid residues with red asterisks marking critical changes relative to florigens. Green color marks identical residues. Red asterisks in (c) mark premature stop codons.

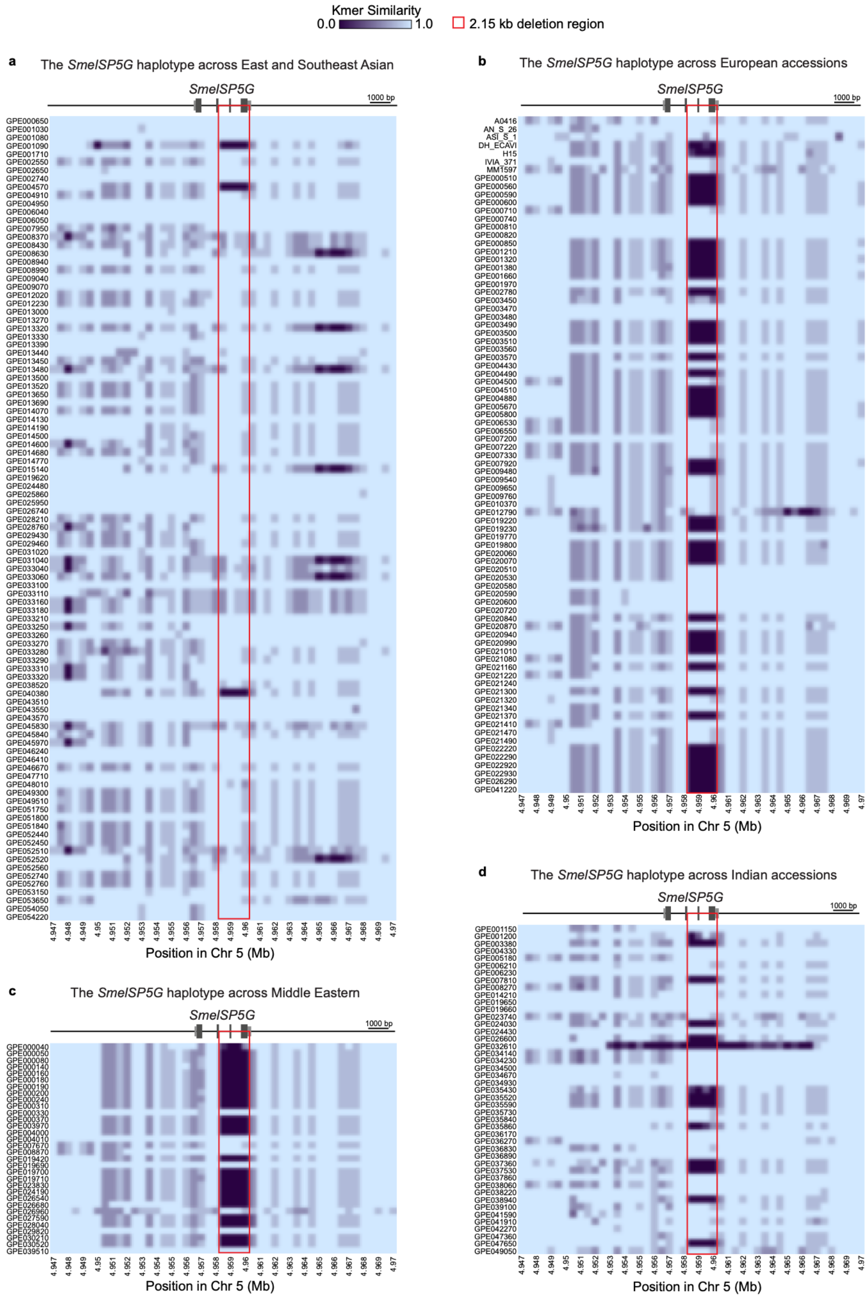


**Supplementary Figure 4. The frequency of the *SmelSP5G* coding deletion allele (*Smelsp5g$^{del}$*) is widespread in European and Middle Eastern accessions. (a-d)** K-mer-based *Panagram* visualization of the *SmelSP5G* haplotype in 250 Brinjal eggplants accessions from East and Southeast Asia (a), Europe (b), Middle East (c), and India (d). Haplotype sequences were represented using binned k-mer similarity values, with similarity displayed as the degree of sequence divergence (purple) from the reference accession GPE001970[46] in 500-bp bins. Geographical origins were assigned based on passport data reported by Gaccione et al., (2025)[46], summarized in **Supplementary Table 5**.

**Supplementary Discussion**

Our findings prompted us to ask whether the similar evolutionary patterns of paralogue neofunctionalization and subsequent phenotypic changes reflect a broader rule of adaptive significance, extending beyond flowering time and Solanaceae evolution. We therefore searched for examples across biological systems that share three fundamental features:
(i) the emergence of a paralogue following gene duplication;
(ii) neofunctionalization through acquisition of an opposing function relative to the ancestral paralogue; and
(iii) Contribution of the neofunctionalized paralogue in subsequent phenotypic change.

Our search was not exhaustive, focusing on identifying a range of taxa that differ substantially in developmental contexts and phylogenetic distances to reveal conceptual parallels. Below, we highlight three divergent examples spanning plant morphology, insect wing development, and human brain evolution.

***Leaf complexity in plants***
A more ancient example resembling *SP5G* is found in the evolution of the *KNOTTED-like* homeobox (*KNOX*) transcription factor genes controlling leaf complexity in land plant[120]. Leaf complexity refers to the degree to which leaves are subdivided into lobes or distinct leaflets, ranging from simple undivided leaves to highly compound architectures such as those found in tomato[121]. Evolutionary changes in leaf complexity often arise through modulation of developmental programs controlling the duration of meristematic activity during leaf formation[122]. An ancient duplication generated the *KNOX1* and *KNOX2* clades, the latter of which diverged into an antagonistic developmental node[120]; *KNOX1* clade members promote meristematic activity and leaflet formation, whereas *KNOX2* genes promote differentiation and repress meristematic state, thereby reducing leaf complexity. This antagonistic configuration expands developmental plasticity and facilitates morphological diversification across land plants[123]. Importantly, repeated evolutionary changes in leaf complexity across flowering plants, including tomato, legumes, and *Cardamine*, have been associated with recurrent redeployment and modulation of *KNOX* regulatory activity[124-127]. Thus, similarly to *SP5G* within the Solanaceae *florigen* network, the emergence of ancient antagonistic function between *KNOX* paralogues was repeatedly targeted during plant morphological evolution.

***Wing polyphenism in insects***
A remarkably similar pattern of antagonistic neofunctionalization has been described in wing polyphenism of true insects (Hemiptera) through the emergence and neofunctionalization of paralogous insulin receptor genes (InRs)[128]. Wing polyphenism refers to the environmentally or genetically regulated production of distinct wing morphs within the same species, typically involving a tradeoff between long-winged dispersal forms and short-winged or wing-reduced forms optimized for reproduction or local adaptation[129]. These alternative wing strategies allow insects to balance dispersal capacity with reproductive investment in response to ecological conditions, thereby contributing to adaptive flexibility across environments. The ancestral InR1 protein retained canonical insulin-signaling activity promoting long-wing morph

development, whereas InR2 evolved antagonistic regulatory activity that suppresses InR1 signaling through direct receptor interaction, thereby promoting the short-wing morph[130]. This antagonistic insulin perception presents a recurrent regulatory mechanism underlying wing polyphenism across multiple species[128,130]. Thus, similarly to *SP5G* in the Solanaceae, duplication and the subsequent emergence of antagonistic receptors within the insulin signaling pathway repeatedly contributed to adaptive phenotypic plasticity in insects.

***Human neocortical development***

A final example appeared in primates with the human-specific duplication of the neuronal signaling regulator *SRGAP2*, a membrane-associated protein controlling neuronal migration and synaptic maturation[131]. Following partial duplication of the ancestral *SRGAP2A* gene, the derived paralogue *SRGAP2C* evolved as a truncated protein that antagonizes the activity of its ancestral counterpart through heterodimerization[132]. Whereas *SRGAP2A* promotes neuronal maturation and limits dendritic spine density across mammals, *SRGAP2C* delays human synaptic maturation and enhances spine density and plasticity, features associated with the evolution of neocortical development[131]. This prolonged period of neuronal immaturity, sometimes described as developmental neoteny, is thought to extend cortical plasticity and increase the capacity for circuit refinement during human brain development[133]. Experimental studies further demonstrated that expression of *SRGAP2C* induces cortical neuronal phenotypes resembling aspects of human-specific brain development[131], highlighting how recent duplication and antagonistic functional divergence can remodel conserved developmental programs during evolution.
